# Detecting rhythmic spiking through the power spectra of point process model residuals

**DOI:** 10.1101/2023.09.08.556120

**Authors:** Karin M. Cox, Daisuke Kase, Taieb Znati, Robert S. Turner

## Abstract

**Objective:** Oscillations figure prominently as neurological disease hallmarks and neuromodulation targets. To detect oscillations in a neuron’s spiking, one might attempt to seek peaks in the spike train’s power spectral density (PSD) which exceed a flat baseline. Yet for a non-oscillating neuron, the PSD is not flat: The recovery period (“RP”, the post-spike drop in spike probability, starting with the refractory period) introduces global spectral distortion. An established “shuffling” procedure corrects for RP distortion by removing the spectral component explained by the inter-spike interval (ISI) distribution. However, this procedure sacrifices oscillation-related information present in the ISIs, and therefore in the PSD. We asked whether point process models (PPMs) might achieve more selective RP distortion removal, thereby enabling improved oscillation detection.

**Approach:** In a novel “residuals” method, we first estimate the RP duration (*n_r_*) from the ISI distribution. We then fit the spike train with a PPM that predicts spike likelihood based on the time elapsed since the most recent of any spikes falling within the preceding *n_r_* milliseconds. Finally, we compute the PSD of the model’s residuals.

**Main results:** We compared the residuals and shuffling methods’ ability to enable accurate oscillation detection with flat baseline-assuming tests. Over synthetic data, the residuals method generally outperformed the shuffling method in classification of true-versus false-positive oscillatory power, principally due to enhanced sensitivity in sparse spike trains. In single-unit data from the internal globus pallidus (GPi) and ventrolateral anterior thalamus (VLa) of a parkinsonian monkey -- in which alpha-beta oscillations (8-30 Hz) were anticipated -- the residuals method reported the greatest incidence of significant alpha-beta power, with low firing rates predicting residuals-selective oscillation detection.

**Significance:** These results encourage continued development of the residuals approach, to support more accurate oscillation detection. Improved identification of oscillations could promote improved disease models and therapeutic technologies.

## Introduction

To estimate oscillations in neural activity, it is conventional to compute the power spectral density (PSD) of measures of extracellular fields (e.g., local field potentials (LFPs), or electrocorticograms (ECoG)). Such signals reflect a complex mixture of contributions from neuronal populations [2], and are therefore not optimal when the objective is to investigate oscillations at the level of individual neurons. In these situations, we can attempt to apply spectral analysis to single-unit spike trains. Ideally, the resulting PSD output – along with post-processing and statistics -- should enable the inference of oscillations in the rate function that generated the observed spikes [3]. We adopt the working definition of an oscillation as an approximately sinusoidal, narrowband modulation in action potential (“spike”) occurrence.

To detect oscillatory components in a spike-derived PSD function accurately, we must first address a challenge: These spectra exhibit a distortion pattern that is common to phenomena subject to measurement “dead time” (i.e., an interval after an event, when the occurrence or detection of a subsequent event is impeded [4]). In the context of single-unit spike trains, this dead time is typically attributed to the neuronal refractory period, which may be absolute (*p*(spike) = 0) or relative (gradually recovering *p*(spike)). The simplest example of this distortion occurs in the context of a hypothetical spike train arising from a Poisson process interrupted by an absolute refractory period. In this case, the resulting PSD departs from the flat spectrum anticipated for an unmodified Poisson process, and is instead characterized by a trough over a range of low frequencies, followed by elevated power (often accompanied by visible ringing) over the high frequencies [5, 6]. The precise form of the distortion differs modestly for relative refractory periods (see Fig. 1 for a synthetic example), and is exacerbated by high firing rates (FR) and long refractory period durations [1].

**Fig. 1.**
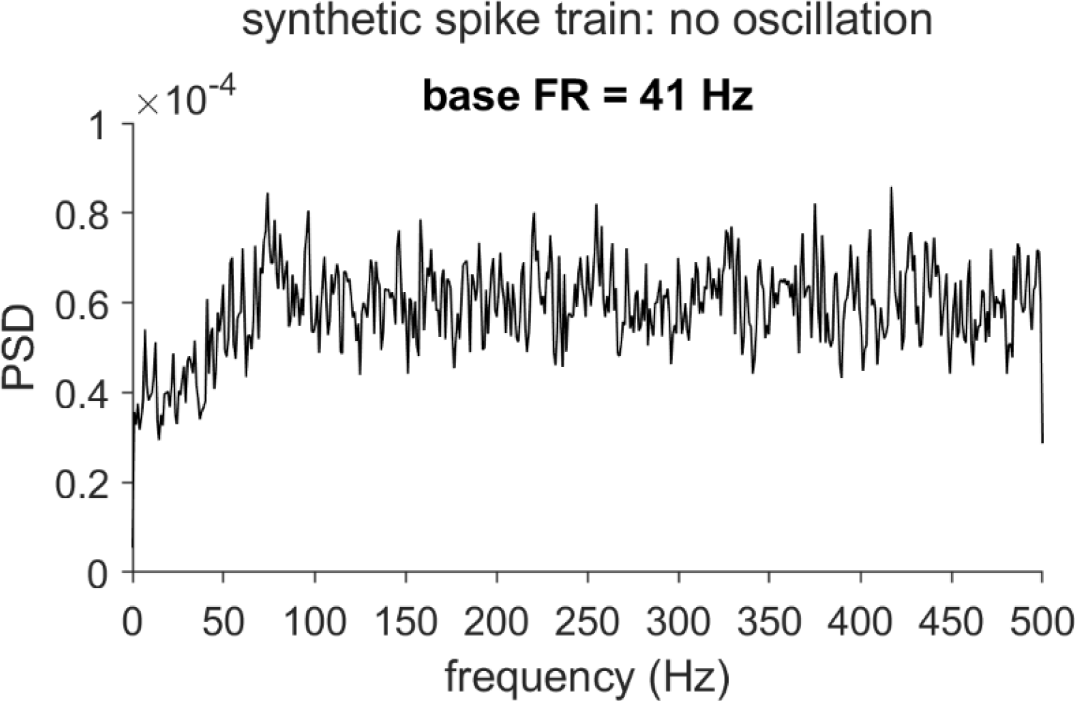
Recovery period distortion of the power spectrum for a synthetic spike train with no oscillation. Power spectral density (PSD) of a spike train simulated under the assumption of Poisson spiking interrupted by a roughly exponential recovery period (RP, the post-spike reduction in spike probability, which can extend beyond the refractory period). Relative to the flat spectrum of a Poisson process, this RP-distorted PSD exhibits a characteristic trough over the lowest frequencies, and relative elevation and ringing over the higher frequencies. The PSD was generated according to the simulation framework from [1] (see Methods; baseline firing rate (FR) = 41 Hz; RP duration n_r_ = 9 ms and steepness k = 0.7; simulation duration T = 60 × 1024 ms).

The present research examines two methods – including a commonly-used approach (the “shuffling method”), and a novel alternative (the “residuals method”) – that aim to remove this dead time-induced distortion from spike train spectra, and ultimately facilitate more sensitive and selective detection of the periodic effects of interest. Beyond these two options (and an analytical variant on the shuffling approach [7]), we are aware of no additional methods specifically designed to correct distortion in neuronal spike train spectra. Therefore, we limit our comparison to these two methods.

Before describing the correction methods, it is necessary to clarify a terminology choice: Instead of attributing the dead time to the refractory period, we will use a mechanism-neutral term: “recovery period” (see [8] for similar usage). In data from our lab (including the empirically-sampled units we present here), we have observed several examples of units that exhibit a dead time (i.e., sharp post-spike drops in the hazard function) that far exceeds the typical duration of the channel dynamics responsible for refractoriness (5-10 ms [9]). Such long post-spike pauses might be generated by either circuit-level or intrinsic mechanisms (such as those underlying pacemaker patterns [10]). Since neither of the methods that we discuss attempt to selectively correct for the biological refractory period *per se*, we adopt the generic recovery period (RP) terminology.

When a spike train features an oscillation, the corresponding peak in the PSD is superimposed on the recovery period-associated distortion pattern (see Fig. 2(a) for synthetic examples). The distortion complicates standard statistical procedures that aim to distinguish such oscillatory PSD peaks from baseline by using a null hypothesis model that is estimated from the spectrum itself. In particular, detection problems arise when the baseline does not account for the RP contribution.

**Fig. 2.**
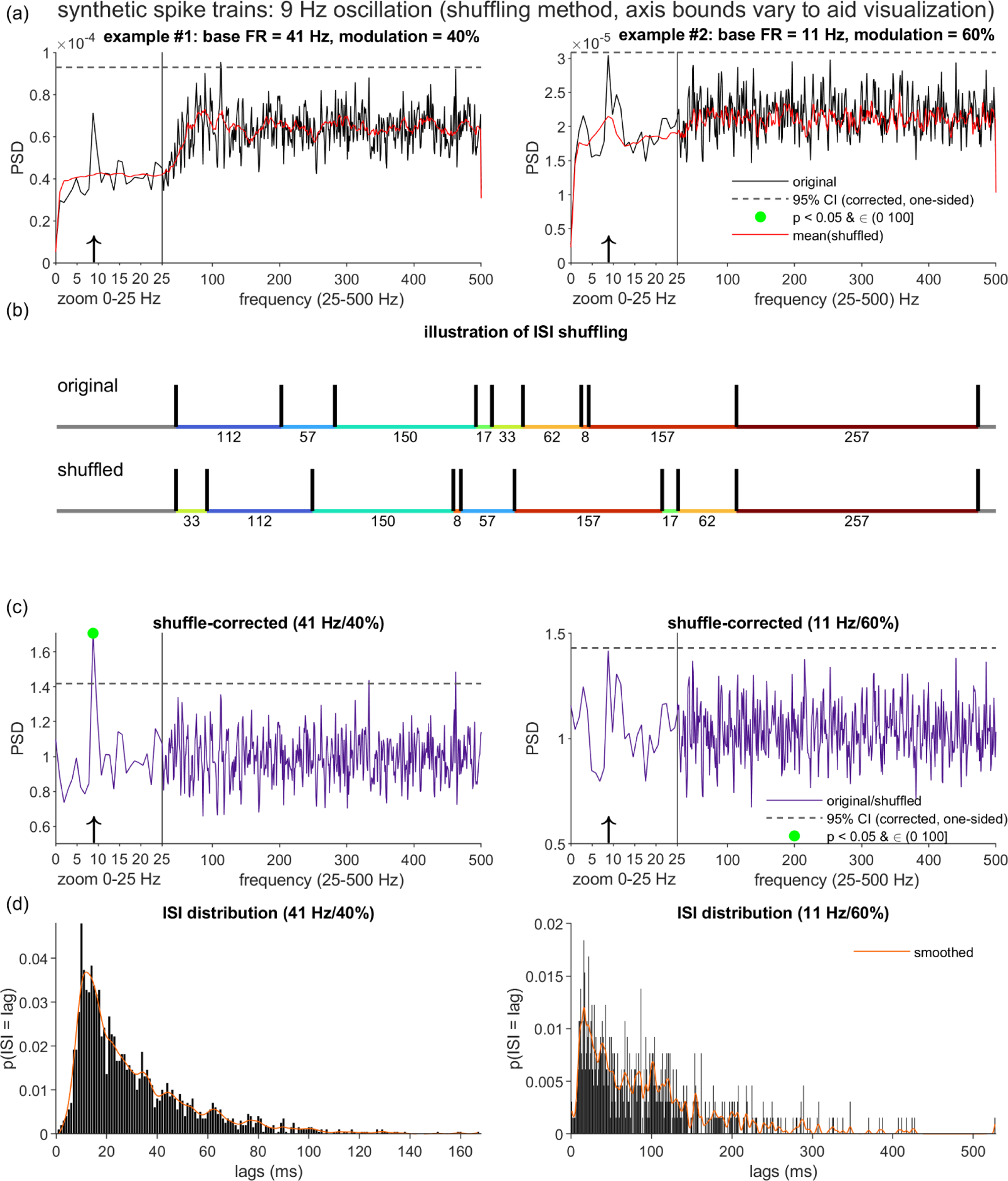
Recovery period distortion of the PSDs for synthetic spike trains with oscillations, and the limitations of inter-spike interval-based correction. (a) PSDs (black lines) for two synthetic spike trains, generated using the same RP parameters and simulation durations as Fig. 1, and added oscillation terms of matching oscillation frequency (*f_osc_* = 9 Hz). The spike trains differed with respect to the oscillation modulation strength (*m*) and the baseline firing rate (base FR; see column titles). Dashed lines mark the corrected 95% one-sided confidence interval (CI) constructed under an approximation of a null model of Poisson spiking (see Methods). Green dots (absent in (a)) indicate PSD points that both cross the CI bound and fall within an a priori search range of (0, 100] Hz. Red lines plot the spectral component that the ISI shuffling algorithm [1] estimates to be attributable to the structure present in the inter-spike intervals (ISIs). (b) Illustration of one iteration of ISI shuffling on a truncated 1000 ms dataset (actual implementation applied global shuffling to the full spike train duration). The average PSD of several shuffled spike trains forms the estimated ISI-attributable spectrum. (c) The shuffling-corrected PSDs, equal to the ratio of the original and shuffling-estimated PSDs. (d) ISI probability distributions for the two example spike trains. Smoothing spline fits (orange lines) highlight the more visible shaping of the ISI distribution by the 9 Hz oscillation in the lower-FR case. Note that the concurrent shaping of the ISIs by a high frequency rhythm, which is readily visible in both the higher- and lower-FR examples, is attributable to the 9 ms relative recovery period.

An especially detailed treatment of RP-associated distortion [1] -- which also introduced the established shuffling method -- illustrated the oscillation detection challenges with a straightforward statistical test that assumes a Poisson process (in the time domain) and corresponding white noise (in the frequency domain) as the baseline. The authors observed that, over the highest frequencies (beyond the frequency bands typically prioritized by neuroscientists), the PSD functions of neuronal spike trains tended to approach those of matched-FR Poisson processes, thereby motivating the construction of confidence intervals (CI) based on the mean and variance over these high-frequency points. Points in the PSD function that cross the CI bound (see Fig. 2(a), dashed lines, for CI examples) are labeled as significant, with the possible added requirement that the points also fall within an *a priori* search range (set at (0, 100] Hz in the figures). Because this statistical test does not account for the RP distortion, it is prone to miss oscillatory peaks that fall within the low frequency trough (see the two examples in Fig. 2(a)), and incorrectly label as significant points at higher frequencies, beyond the trough, which exhibit elevated power as a consequence of the distortion effect alone [1]. Although the examples in Fig. 2(a) do not show false positives within the (0, 100] Hz search range, the PSD in the left panel does include a peak that crosses the CI threshold at ≈112 Hz. Note that this general pattern of false negatives and positives also appears when the CIs for the Poisson null hypothesis are estimated using an alternative analytical procedure (see [1] and [11] for details).

A switch to a different, but still RP-neglecting baseline model is insufficient to resolve these issues. For example, consider the common practice of fitting a “1/f-like” function to the spectra of field potential measures, in order to separate an assumed aperiodic baseline from periodic effects [12]. Although this practice can be effective for such population-level signals – on which individual units’ RPs will not have a discernable impact – it is unclear how a 1/f-like function could be successfully fit to a raw spike train spectrum.

To account for the RP distortion, the correction method proposed by [1] builds upon the null hypothesis that the spike train emerges from a renewal process with a matched inter-spike interval (ISI) distribution. Recall that a renewal process is a sequence of events (here, spikes) for which the waiting times (the ISIs) are independently sampled from an identical distribution [13]. Since we define the recovery period as an influence solely on the waiting time between two successive events, and not on any higher order structure (e.g., correlations between successive waiting times), the renewal null hypothesis does capture the temporal structure attributable to the RP.

The ISI shuffling method described by [1] estimates the renewal equivalent spectrum for a spike train. According to the simplest, global version of this scheme, several control spike trains are generated by randomly permuting the ISIs of the original. The PSDs computed from these control spike trains are then averaged to estimate the renewal equivalent spectrum. An analytical alternative to this Monte Carlo procedure does also exist, which leverages an established method to compute the renewal process spectrum from an ISI distribution [7, 14]. Although this analytical approach is fast and non-stochastic, it does require the use of rectangular spectral analysis windows. For the present work, we focus on the shuffling approach, given its flexibility to tapered windows (e.g., Hamming windows, or the Slepian sequences used by the multitaper method [15]).

Regardless of whether shuffling or the analytical method is used, the resulting control spectrum is then divided out from the original spectrum to obtain the corrected PSD. Given a corrected spectrum, it is customary to proceed with the aforementioned procedure of constructing CIs based on the high frequencies, under the assumption that the null hypothesis of a flat spectrum is now appropriate.

Shuffling correction continues to see common use in single-unit studies, especially in research on the motor control circuitry (e.g., [16–20]), but also in other domains [21, 22]. The method is a well-justified option when researchers are willing to forgo the temporal information present in the ISI distribution. However, as noted by [1], the ISIs can carry information about the rhythmic occurrence of individual spikes, which the shuffling method will remove. A simple, rigid example of this occurs in the case of precisely timed pacemaking activity – that is, single spikes separated by a relatively consistent period *P*, plus or minus a modest amount of jitter (i.e., an approximation of a Dirac comb [23]). In such cases, the pacemaking interval is indistinguishable from a recovery period, implying that such patterns should be detected through means other than RP-corrected spectra (e.g., by seeking instances of extremely low ISI variance). A more subtle example occurs when rhythmic spiking appears to be driven by an approximately sinusoidal function, but the mean FR is so low that spikes often occur only once per oscillation cycle, and may at times skip cycles altogether. To detect and statistically validate such cases, spectral analysis may prove more helpful, but how to effectively address RP distortion while preserving the sparse information regarding the underlying rhythmicity remains unresolved.

Here we explore a new “residuals” method which aims to selectively remove RP-associated variance from a spike train, while leaving intact the information present in the long-lag ISIs. The specific method we describe should be seen as one instantiation of a general, mixed time- and frequency-domain framework that can be adapted flexibly to accommodate analysis goals that are more complex than those prioritized here. We also note that the method represents the combination of a number of existing ideas and techniques from prior work that has sought to estimate spike rhythms exclusively in the time domain (e.g., [24–26]).

The residuals method consists of three stages. First, similar to previous works [24, 27], we use the ISI distribution to inform an estimate of the RP duration (*n̂_r_*) for the input spike train. Second, we construct a point process model (PPM) that predicts the time-varying spike probability (*p_spk_(t)*) as a function of the time elapsed since the most recent spike, but only when a spike was observed within the last *n̂*_*r*_ ms; outside of this RP interval, the prediction relies on the default constant term alone. In other words, this “bounded last-spike” model estimates the component of the data that can be explained by the base firing rate and the first *n̂*_*r*_ ms of the estimated post-spike hazard function.

The fitted PPM is assumed to account for the variance associated with the RP, thereby implying that the raw residuals (actual - predicted values) should represent a “corrected” time series. Therefore, as the third step, we submit the residuals to spectral analysis. The resulting PSD is subjected to the same statistical procedures as were described for the shuffling-corrected spectra.

To our knowledge, no prior work has applied such an approach to the analysis of spike train oscillations. However, we do note some conceptual relatedness to a two-step PPM fitting procedure reported by [24], which was used to account for slow fluctuations in spike rate prior to seeking oscillations in the time domain.

We present a comparison of the shuffling and residuals methods on both synthetic and real neuronal spike trains. In the synthetic data – for which ground truth is certain -- we report a general advantage for the residuals method in the accurate classification of true and false oscillatory points in the corrected spectra. In follow-up analyses, we find that the residuals method is particularly advantageous in the detection of true oscillations, especially when these are of a moderately strong amplitude, and arise from units that fire at rates only modestly greater than the frequency of the oscillation to be detected.

The neuronal data were acquired from the internal globus pallidus (GPi) and ventrolateral anterior thalamus (VLa) of a parkinsonian monkey, thereby presenting a dataset for which the *a priori* expectation of pathological oscillations is high (specifically in the 12-30 Hz beta- and 8-12 Hz alpha-frequency ranges [28, 29]), and for which spiking may tend to be sparse (particularly within the VLa, [30]). As expected for biological data, the findings are complex. We do find several instances of candidate alpha-beta oscillations that are detected by the residuals method exclusively, especially amongst the slowest-spiking units. At the same time, we also find a small number of alpha-beta oscillations flagged exclusively by the shuffling method. We additionally observe aperiodic spectral components (1/f-like trends and probable bursting) that residuals but not shuffling correction leave intact, and propose options for expansion upon the residuals method that could account for these additional phenomena.

## Methods

### Ethics Statement

The single nonhuman primate (NHP) dataset analyzed in this study (see “Experimental Data Methods”) was acquired under a protocol approved by the Institutional Animal Care and Use Committee of the University of Pittsburgh (#18093682). Experimental procedures and animal care complied with the National Institutes of Health Guide for the Care and Use of Laboratory Animals, the PHS Policy on the Humane Care and Use of Laboratory Animals, and the “Guiding Principles for Research Involving Animals and Human Beings” of The American Physiological Society. The animal was housed with a single cage mate in a climate-controlled room, received regular enrichments, and consumed a diet that included fresh fruits and vegetables daily. Surgical procedures were conducted under general anesthesia, with analgesics administered post-operatively.

### Methods Overview

In the subsections that follow, we first describe the procedures for comparing the two correction methods (shuffling and residuals) on synthetic spike trains. These include the steps for generating the spike trains, calculating the original and corrected PSDs, and evaluating the methods’ accuracy in detecting known oscillations. Subsequently, we describe the experimental procedures used in acquiring the NHP data, and also the few details of PSD analysis that differed between the NHP data and the synthetic spike trains.

All simulations and the majority of analyses were implemented in Matlab (R2022a, The Mathworks, Inc., Natwick, MA; RRID:SCR_001622). A limited number of analyses were implemented in R (R Core Team, 2021; RRID:SCR_001905). Matlab and R code sufficient to reproduce the reported results is available on Github (https://github.com/kc13/residuals_spectral_analysis; 10.5281/zenodo.10867519, [31]). We make use of conventional Matlab notation for referring to the ranges spanned by / increments separating vector elements ([min:step:max]).

Data files are available in both a Zenodo repository (DOI: 10.5281/zenodo.8313070, [32]) and the Github repository (see also the Data Availability statement). The original synthetic datasets and NHP spike train data are available on Zenodo. The Github repository stores copies of the NHP spike train data, and also synthetic datasets that are represented in a postprocessed state (the subsampling and pROC output, see “Comparison of ROC curves for the Correction Methods”; note the original synthetic datasets exceed Github file size limits).

### Synthetic Data Methods

#### Spike Train Simulation

We generated synthetic spike trains through the procedure described in [1]. For each time bin *t* (Δt = 1 ms), the spike probability *p_spk_(t)* was determined according to the discrete-time approximation of an inhomogeneous Poisson process:

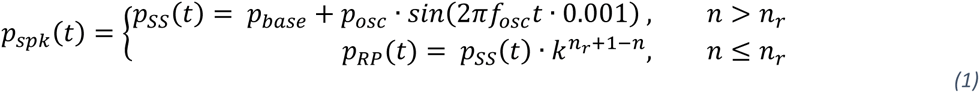

Here, *n* indexes the millisecond latency since the last spike occurrence, and *n_r_* denotes the duration of the recovery period (RP). Therefore, the spike probability at *t* depends on whether the simulated unit is in the recovery period (RP) or has recovered the steady state (SS) firing pattern. During the steady state, *p_spk_(t)* reflects the sum of two components: (1) *p*_base_, corresponding to the mean steady state firing rate (FR), and (2) an oscillation term, in which 0 ≤ *p_osc_* ≤ *p*_base_ governs the oscillation amplitude, and *f_osc_* (specified in Hz) indicates the oscillation frequency. Following from [3], we set *p*_*osc*_ indirectly through variation of the modulation index, *m = p_osc_*/*p_base_*; i.e., the ratio of the peak and mean steady state FRs, respectively. The above *p*_osc_ constraint implies that 0 ≤ *m* ≤ 1. For all simulations, *p_base_* << .5, thereby implying that *p_spk_(t)* remained bounded in [0,1].

During the recovery period, *p_spk_(t)* is set equal to the steady state probability following modulation by an exponential recovery term, where 0 ≤ *k* < 1. Following from [1], our simulations adopted default RP parameter settings of *k* = 0.7 and *n_r_* = 9; a set of follow-up tests considered the use of other values (see “Evaluation of the PSD Correction Methods”).

The duration of the synthetic spike trains (*T*) was constrained to multiples of 1024 ms (the segment size used for spectral analysis). The settings used for *T* and three other free parameters (*p_base_*, *m, f_osc_*) are described in further detail in “Evaluation of the PSD Correction Methods”. The output of each simulation run was a *T* × 1 binary vector (a “delta vector”), indicating time bins of spike occurrence at a resolution of 1 kHz.

#### Uncorrected PSD

To obtain the initial, uncorrected PSD estimates, we submitted the delta vectors (demeaned segment-wise) to Welch’s method (Matlab *pwelch*, Hamming window length = NFFT = 1024 ms, Fs = 1 kHz, zero segment overlap). PSD functions were represented as 513-element vectors spanning the frequency range [0, 500] Hz, implying a frequency resolution of *Δf* ≈ 0.9766 Hz.

Within each PSD function, we sought points of statistically significant power in the range (0,100] Hz. Significance was determined using the CI-construction procedure from [1] which was described in the Introduction. For our analyses, points in the (0, 100] Hz range of a spectrum were labeled as significant if they exceeded the upper limit of a one-sided, [100*(1-α_c_)]% confidence interval formed using the mean and SD over the [250, 500] Hz range of the same spectrum (α_c_ = Bonferroni-corrected significance level, with respect to the 102 points in the (0,100] range). All figures of uncorrected PSDs were generated with α_c_ = .05.

For the shuffling and residuals methods detailed below, all spectral analysis and statistical testing steps made use of the same routines as were applied to the uncorrected PSDs, and the same default α_c_ = .05 for figure generation; α_c_ was varied away from this default for the ROC analyses described in “Evaluation of the PSD Correction Methods”.

#### Shuffling-corrected PSD

To obtain the shuffling-corrected PSD for a given spike train, we first generated 100 surrogate, ISI-shuffled versions of the original delta vector (Fig. 2(b)). We implemented the global shuffling method described by [1] (i.e., random permutation of all ISIs across the entire vector). We chose this approach over local shuffling (permutation of ISIs within nonoverlapping segments) since our simulations emphasized relatively low firing rates (necessitating a broad shuffling scope to admit a sufficient number of ISIs) and did not use time-varying simulation parameters (eliminating the need to capture local variations). Additionally, using the same primary evaluation that we describe for the shuffling and residuals methods (see “Evaluation of the PSD Correction Methods”), we verified superior performance for global as compared to local shuffling. Code sufficient to reproduce this observation is available in the Github repository.

Each of the 100 globally shuffled spike trains was submitted to spectral analysis, and the mean of the resulting PSDs was divided from the original uncorrected PSD, producing the final shuffling-corrected PSD.

#### Residuals-corrected PSD

The residuals method entails three steps: (1) estimation of the recovery period duration, (2) fitting a point process model (PPM) to the spike train, which estimates the effect of the time elapsed since the most recent spike, up to a history bound corresponding to the estimated RP duration, and (3) applying spectral analysis to the residuals of the PPM fit.

### Estimation of Recovery Period Duration

We define the recovery period as a post-spike reduction in a unit’s spike probability, terminating upon return to a steady state firing rate. For our synthetic spike trains (excluding a series of follow-up datasets), the ground truth duration of the recovery period was *n_r_* = 9 ms.

Our estimation approach builds upon a general logic that was described in [27], although note that these authors did not propose an explicit estimation algorithm. We expand upon these authors’ ideas by detailing a specific algorithm that operates on the probability density function (PDF) of the inter-spike intervals (ISI). To introduce our approach, we use a simple synthetic spike train for which *m* = 0 (i.e., no oscillation present, such that firing is shaped by only *p*_base_ and the RP); however, subsequent analyses will show that the algorithm performs reasonably when *m* > 0.

The ISI PDF for an example spike train generated with *m* = 0 is shown in Fig. S1(a). Recall that, in the case of Poisson firing (i.e., *n_r_* = 0, with an intensity λ = *p*_base_), the observed ISIs are sampled from an exponential PDF [33]: *P(ISI = x) = λe^−λx^*. Because a spike train with an RP is not Poisson, an exponential curve fit to such a spike train’s ISI PDF will overestimate the probability mass assigned to the ISI lags ≤ *n*_r_. Now, consider what we should observe if we progressively shift the exponential curve fit rightward. Fig S1(a) illustrates this process for a series of example lags *L* assigned to the *x* = 1 position of the exponential fit. When *L* ≤ *n*_r_, the exponential curve continues to overestimate the probability mass for the lowest lags. However, when *L* = *n_r_* + 1, the exponential model of this left-cropped ISI distribution is appropriate, as firing outside of the RP is Poisson. Our strategy for identifying the boundary between the end of the RP and the start of the steady state relies on this expected transition. We additionally leverage the observation that statistical evidence in favor of the exponential fit (i.e., the extent to which the addition of an exponential curve explains variance beyond that explained by a constant term alone) tends to be stronger when *L* = *n_r_* + 1 than when it is incremented to the immediately neighboring lags to the right.

The estimation algorithm proceeded as follows:

1. Given a delta vector, compute the ISI histogram, with bins spanning the ms lags in [1, max(ISI)].
2. For each *L* in *lags*:

a. Extract the histogram values over the range [L, max(ISI)].
b. Normalize the extracted histogram to unit area to form a PDF.
c. Fit a single scaled exponential to the PDF using the generalized linear model (GLM) of the form 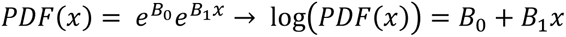. See the next subsection for more detail on this form of GLM.
d. Compute the deviance difference, which is a standard statistic for assessing the improvement in goodness-of-fit contributed by a full GLM (“model 1”, here set to the constant term plus the exponential lag effect), compared to a GLM restricted to the constant *B_0_* term (“model 0”). The difference statistic is Δ*D* = *D*_0_ − *D*_1_ = −2[log *L̂*_0_ − log *L̂*_1_]; i.e., the log likelihood ratio scaled by −2 [33, 34].
e. Upon confirming the first instance of a *ΔD* value that is a local maximum (*ΔD(L-1) < ΔD(L) > ΔD(L+1*)), break out of the loop iterating over *lags*.
3. The L associated with the local *ΔD* maximum marks the start of the steady state region; the estimated recovery period duration, *n̂*_*r*_, is set to L-1 ms.

For the synthetic unit with *m* = 0, this process of finding the lag *L* associated with the local *ΔD* maximum is illustrated in Fig. S1(b). Note that in this case, the duration of the recovery period is accurately estimated as L-1 = 9 ms.

Figures 3(a)-(b) illustrate the same process as applied to the two synthetic oscillatory spike trains introduced in Fig. 2. Although the presence of non-zero oscillation (i.e., *m* > 0) implies that the steady-state firing is not Poisson, we observe that the curve-fitting algorithm is robust to this departure and again correctly identifies the ground truth RP duration of 9 ms. Evaluation of the RP estimation accuracy over a full dataset of synthetic spike trains is presented in the Results.

**Fig. 3.**
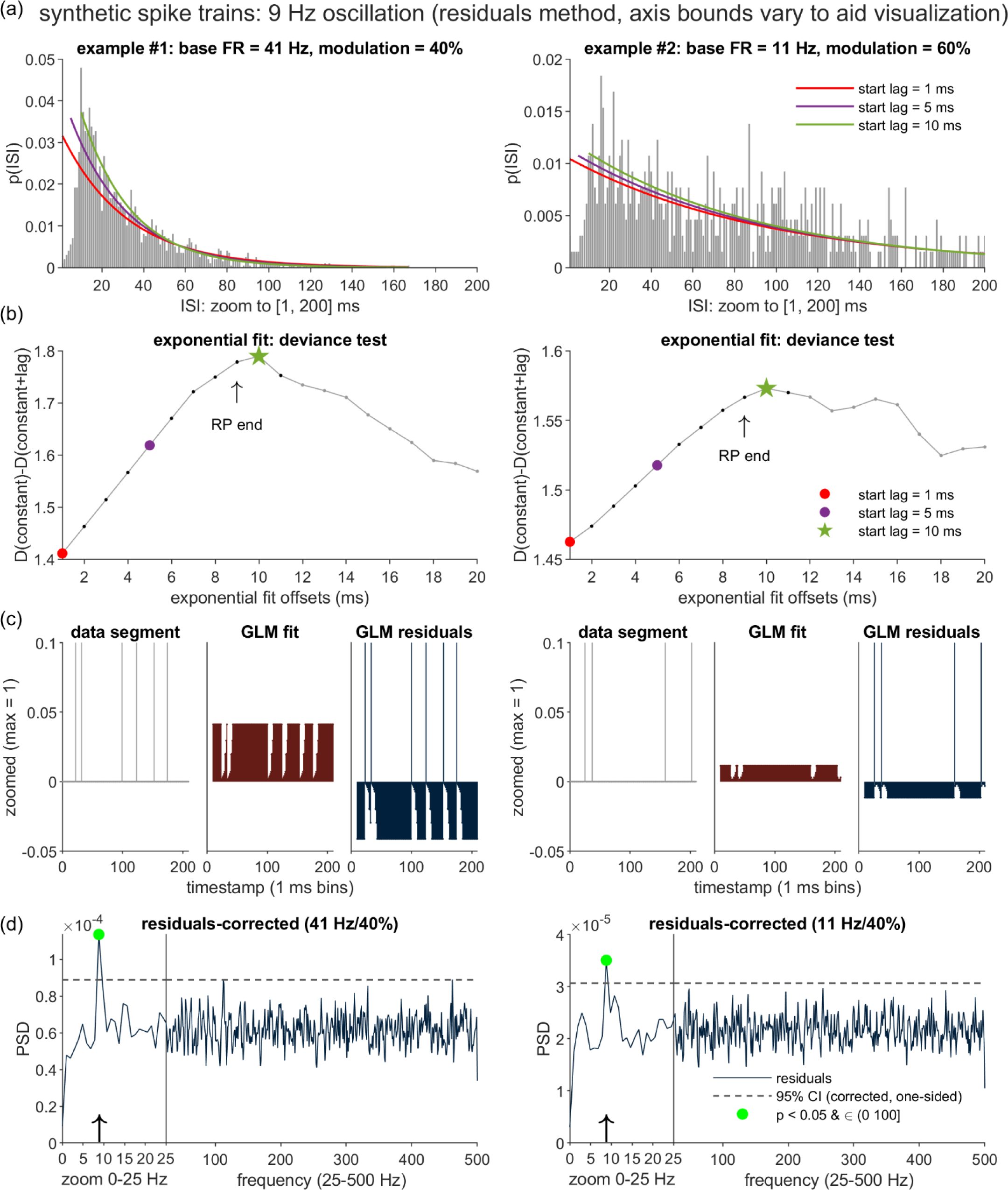
Distortion correction by the proposed residuals method. (a) Illustration of the procedure for obtaining an estimate of the RP duration (*n̂*_*r*_), as applied to the two example spike trains from Fig. 2. A series of right-shifted exponential curves are fit to each ISI distribution, left-anchored to starting positions advanced in 1 ms steps (with 3 sample iterations highlighted in the figures). (b) Plots of the deviance difference statistic, ΔD, as a function of the first 20 starting positions of the exponential fits. D(constant), D(constant+lag) = deviance measures for the intercept-only and intercept+exponential curve models, respectively. ΔD tracks the goodness of fit contributed by the exponential curve. Each *n̂*_*r*_ estimate is set equal to the post-spike lag immediately preceding the first local maximum in the corresponding ΔD plot. (c) Illustration of the Poisson Generalized Linear Model (GLM) accounting for the effects on spike likelihood of the time elapsed since the most recent of any spikes occurring within the preceding *n̂*_*r*_ ms. The three panels (which highlight the first 209 ms of data) depict the GLM input (left panels), model fit (middle panels), and the model residuals (right panels). (d) PSDs of the GLM residuals. Plotting and statistical conventions follow those from Fig. 2.

Note that steps 2(e)-3 of the above algorithm can be equivalently implemented by (1) applying significance testing to each Δ*D* value (i.e., a χ^2^ test, with a single degree of freedom, reflecting the difference in the number of parameters between model 0 and model 1; [33]), and (2) seeking the first local minimum in the resulting *p* values.

### Bounded Last-Spike Point Process Model

Once the RP duration is estimated, the next step is to estimate the RP shape. For this purpose, we fit the spike train with a point process model (PPM).

With reference to [8, 24, 33, 35, 36], we will briefly review the fundamental concepts and terminology for point process models in general, and for our implemented Poisson GLM in particular. A PPM seeks to predict the likelihood that a discrete event (e.g., a spike) will occur at time *t*, given a set of covariates. For a stochastic point process represented in continuous time (e.g., a spike train expressed as a series of spike timestamps), the PPM models the conditional intensity function (CIF):

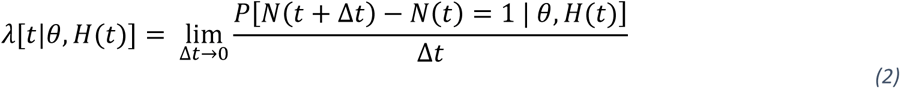

where *P*[*N*(*t* + Δ*t*) − *N*(*t*) = 1 | *θ*, *H*(*t*)] is the conditional probability of a discrete event instance (e.g., a spike) occurring in the time interval (*t*, *t* + Δ*t*], given a covariate history *H*(*t*) and model parameters *θ*. For our discrete-time spike train representation (the delta vectors), this continuous formulation may be approximated using the assumptions that for *Δt* ≤ 1 ms, *P*(spike in (*t*, t+ *Δt*]) ≈ *λ*[*θ*, *H*(*t*)]Δ*t*. For the remainder of this text, as a shorthand, we will follow the convention of using *t* to index the ms-width bins of the discrete-time representation (as opposed to continuous time).

Given this discrete-time representation, a Poisson GLM (i.e., a generalized linear model with a log link function, also known as Poisson regression) is one example of a PPM that may be used to predict spike probability. We implemented the bounded last-spike model as a Poisson GLM of the form

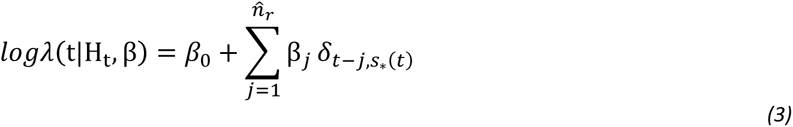

where *s*_∗_(*t*) denotes the time of the most recent spike [35], and *δ*_*t*−*j*,*s*∗(*t*)_ is the Kronecker delta function, returning 1 when *t-j* equals s_∗_(t) and 0 otherwise. In other words, the model estimates the variance in the logarithm of spike intensity that is explained by either the constant term alone (outside of the RP) or the constant plus the weighted contribution of the millisecond interval transpired since the last spike. Consequently, inside the RP, λ is modeled as the multiplicative contribution of the exponentiated constant and last-spike 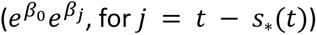, and as 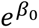 outside of the RP.

The model was fit to the spike train data using Matlab *fitglm,* which applies the conventional iteratively re-weighted least squares (IRLS) method to seek the log likelihood-maximizing parameters. Note that the log likelihood function for a model such as (3) is known to be concave [8], thereby implying that if a maximum exists, it must be unique.

### Spectral Analysis of Model Residuals

Given a set of parameter estimates *β̂*, the raw residuals of the modeled spike train (the observed - fitted values) are calculated as

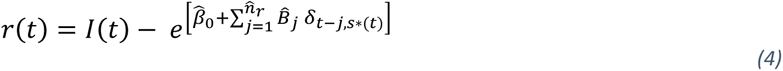

where *I(t)* represents the binary spike train input to the PPM. The final step of the residuals method is submission of *r(t)* to spectral analysis. Note that (4) only defines *r(t)* for *t* ≥ *n̂*_*r*_ + 1 ms. To maintain alignment with the original time series (and the divisibility by the 1024 ms segment size), the initial 1024 ms segment submitted to *pwelch* consisted of the initial 1024-*n̂*_*r*_ ms of computed residuals, demeaned and prepended with zero-padding of *n̂*_*r*_ ms.

### Clarifications and Relationship to Pre-existing PPM Approaches

We conclude this description of the residuals method with two points concerning the “bounded last-spike” PPM in particular. The first point concerns the use of the term “bounded”: Here, we intend to emphasize the bounded, *n̂*_*r*_-millisecond historical window within which the most recent spike must fall, to be eligible to contribute to the spike intensity prediction at time *t*. We make this point to prevent confusion with another common usage of “bounded”: For a discrete-time point process, the count of events that may occur in each sampled time bin is expected to be bounded in [0,1], thereby satisfying the conditions for “orderliness” [37]. Although our model was not named for this type of boundedness, we note that the orderliness expectation is met by the synthetic time series we are analyzing (given the constraints of our simulation framework) and is very likely to be met by our sample of empirically-acquired spike trains (given the typical assumption of a 1 ms lower bound on the neuronal refractory period [33]).

As a second point, we note that the bounded last-spike PPM draws upon multiple features of pre-existing PPMs in the spike modeling literature, although it is also important to clarify how our current implementation differs from these earlier approaches. For this purpose, two categories of pre-existing PPMs are relevant. The models in both categories aimed to account for spike activity relatively comprehensively (in contrast to our model, which aims to selectively capture the RP effect).

A principal representative of the first category is the Inhomogeneous Markov Interval (IMI) model of [35]. The IMI model predicts spike intensity as a function of the time elapsed since the most recent spike (*t* − *s*_∗_(*t*), with no bound imposed) as further modulated by a function of the current clock time (*t*); the two functions’ forms are estimated with regression splines. Therefore, our current model shares the IMI model’s incorporation of a *t* − *s*_∗_(*t*) term, but differs in its exclusion of the clock time term, and also of the *t* − *s*_∗_(*t*) information when *t* − *s*_∗_(*t*) > *n̂*_*r*_. Our PPM also differs in its modeling of the *t* − *s*_∗_(*t*) effect with a series of delta functions, as opposed to the splines used by [35]. We did test a version of the residuals method that replaced the delta functions with cubic splines (specifically, the modified cardinal splines from [36]), and that adjusted the number and location of the knots according to rules that depended on the magnitude of *n̂*_*r*_. The spline-based model was found to perform slightly worse than the delta function version, as demonstrated through the same evaluation procedures as were used to compare global and local shuffling. See the Github repository for code sufficient to reproduce this result.

The second category consists of autoregressive PPMs which predict spike intensity as a function of the spike counts recorded for each of a series of historical time bins (e.g., [25, 26]). Specifically, these models consider a long historical time interval (up to *t* - 150 ms in [25]), divide this interval into time bins of interest, and use the counts of spikes recorded in each bin as individual predictor variables in the PPM. Typically, a subset of narrow (e.g., 1 ms) bins positioned over the most recent historical timepoints (e.g,. −10 through −1 ms) are assumed to capture the combined effects of both the recovery period and any other short-lag history effects (e.g., bursting). Our present approach draws upon these PPMs’ use of indicator functions to model history effects limited within bounded historical intervals, but differs in its exclusive focus on the most recent *n̂*_*r*_ millseconds, and on the timing of the most recent spike that occurred within that historical window.

#### Evaluation of the PSD Correction Methods

Comparison of the residuals and shuffling methods entailed (1) simulation of synthetic spike trains over a range of physiologically-relevant parameters, (2) generation of the two methods’ corrected PSDs, with tabulation of hit and false alarm rates for these PSDs across a range of α_c_ levels, and (3) comparison of the methods’ classification of true and false oscillations, by way of ROC (Receiver Operating Characteristic) analysis (collapsed across parameter settings) and regression analysis of the residuals-shuffling differences in the hit and false alarm rates (to examine parameter effects).

### Synthetic Spike Train Test Sets

Method evaluation was first carried out on a primary synthetic test set; subsequently, we generated a number of additional test sets to address follow-up questions. For each test set, and for each of the unique combinations resulting from the crossing of the varied simulation parameters, we randomly generated one hundred spike trains.

For the primary test set, 100 spike trains were simulated for each of the 540 unique combinations that resulted from crossing the following four varied parameters:

(1) Spike train duration (*T*): {30,60,120} × 1024 ms windows
(2) Oscillation frequency (*f_osc_*): {7,9,12,20,32} Hz
(3) Base firing rate offset (*p_base_offset_*, such that *p_base_* = *f_osc_* + *p_base_offset_*): +2^[0:1:5]^ Hz
(4) Modulation index (*m = p_osc_/p_base_*): [0:0.2:1]

These parameters emphasize relatively low duration and firing rate settings, with the short durations consistent with the limitations that can be posed by real experimental data, and the low FRs in line with the motivation for developing the residuals method. At the same time, the setting of *p*_base_ as a positive offset from *f*_osc_ ensured that, in most cases, the synthetic unit averaged > 1 spike per oscillation cycle, although the recovery period implied that the average dropped to slightly < 1 in some of the *f_osc_* + 1 cases. For *m*, note that the inclusion of 0 allowed for tracking of false positive detections of significant power when no oscillatory effect was introduced into the simulation.

Three kinds of variants on this primary dataset were created to address follow-up questions. First, we generated one dataset in which *p_base_* and *p_base_offset_* were directly manipulated, leaving *f_osc_* = *p_base_* - *p-_base_offset_* as the indirectly manipulated variable. This dataset allowed for tests to determine whether the methods’ relative performance was significantly influenced by the magnitude of the *p_base_ - f_osc_* difference, when controlling for *p_base_* itself. For this dataset, the simulations used the same *T* and *m* settings as were used for the primary dataset. We crossed these factors with two levels of *p_base_* (15 Hz, 20 Hz) and 4 levels of *p_base_offset_* ([2:2:8] Hz), resulting in 144 unique parameter combinations.

The second and third variations involved the use of higher FRs, and alternatives to the default RP parameters, respectively. For the high-FR simulations, the set of sampled *p_base_offset_* values was changed to [35:10:85] Hz; these were added to the *f_osc_* values used for the primary dataset ((2) in the list above) to determine the *p_base_* settings. For the altered-RP simulations, we tested both shorter and longer RP durations (*n_r_*) and varying exponential steepness terms (*k*); full details are described in the Results.

### Labeling of Hits and False Alarms

For each of the 540 × 100 (or 144 × 100) synthetic spike trains in a test set, the shuffling and residuals methods were both applied to produce corrected PSDs. Given a corrected PSD and an α_c_ setting, statistical testing returned a binary vector indicating whether oscillatory power exceeded the CI bound for each of the 102 frequency bins of width *Δf* = 0.9766 Hz in (0,100] Hz. Frequency bins are labeled by their lower bound (0.9766, 1.9531, 2.9297…). Let **f** denote the frequency labels and s(f) denote the binary significance output. We classified a PSD as containing a “hit” (i.e., a true positive detection of oscillatory power) if (1) m > 0 and (2) any s(f) = 1 for those labels *f*_i_ ∊ **f** that were the 3 nearest neighbors of *f*_osc_ (e.g., *f*_i_ ∊ {10.7422,11.7188,12.6953} for *f*_osc_ = 12 Hz). We classified a PSD as containing a false alarm (FA) if either of two cases held true: (1) *m* = 0, s(f) = 1 for any label *f*_i_ ∊ **f**, or (2) *m* > 0, s(f) = 1 for *f_i_* ∊ |*f*_i_ - *f*_osc_| > 5 Hz (equivalent to those *f_i_* outside the 10 nearest neighbors of *f_osc_*).

Note that the output of this classification procedure was a pair of binary labels for each PSD, indicating whether each contained ≥ 1 hit, and ≥ 1 FA, respectively. For the ROC analysis described below, counts of hits and FAs refer to the counts of these PSD-level labels (as opposed to the total counts of individual hit- or FA-classified frequency bins).

### Comparison of ROC curves for the Correction Methods

ROC analysis of the shuffling and residuals methods consisted of three steps: (1) random selection of subsamples of the PSDs, to be used for a bootstrapping procedure for statistical testing, (2) calculation of partial ROC curves and corresponding partial area under the curve (pAUC) values for the subsampled PSDs, and (3) statistical comparison of the pAUC values yielded by the shuffling and residuals methods. Note that the ROC analyses were limited to the *m* > 0 cases, thereby implying that, for those datasets that originally consisted of 540 unique parameter conditions, 450 (unique parameter conditions) *x* 100 (*n* spike trains per condition) were available for ROC analysis.

The subsampling procedure was iterated 1000 times. For each subsampling iteration, we randomly sampled, without replacement, 20 of the 100 synthetic spike trains generated within each of the 450 parameter combinations. The full resulting subsample of 450 × 20 = 9000 spike trains, and the corresponding hit and FA classifications for the shuffling- and residuals-corrected PSDs, were used to generate separate shuffling and residuals ROC curves for that iteration.

For a given subsampling iteration and correction method, we formed an ROC curve as the function relating hits to FAs over a range of α_c_ values. The full set of α_c_ values consisted of the sorted vector of the values 1 and 5 multiplied by a series of negative powers of 10 ([1 5] ^T^ * [10^[−8:1:-1^]) and 1. The curves were formed by collapsing across the simulation parameter combinations, such that a point on an ROC curve represented the fraction of the 9000 corrected PSDs that were associated with ≥ 1 hit and ≥ 1 FA, respectively, at a given *α_c_* level.

For each analyzed dataset, the above procedure resulted in 1000 pairs of shuffling- and residuals-generated ROC curves. Note that the FA and HR ranges for these curves could vary slightly, and fall short of spanning [0,1]. Curves that did not reach 100% HR or FA rates (even at α_c_ = 1) were possible due to the use of a one-sided significance test: If all points in either the hit or FA zones of a PSD were numerically less than the mean of the high frequencies-based CI, then these points could never pass even the most lax significance threshold. Curves that did not drop to 0% HR or FA rates reflected PSDs with prominent hit- or FA-eligible points that survived even an α_c_ = 1 × 10^−8^ threshold (which is much more stringent than the criteria that would be typically applied in practice; e.g., [1]).

The variability in the curves’ spanned ranges precluded analyses that compared the areas under the entirety of the ROC curves. Therefore, we instead adopted the partial area under the curve (pAUC) approach [38], such that we exclusively considered the area under an ROC region that was shared by all 2000 curves. Specifically, the raw curves were cropped (using linear interpolation as needed) to span the intersection of all 2000 curves’ FA ranges. For each cropped curve, we used the trapezoidal rule (Matlab *trapz*) to compute the area under this bounded curve region. Due to the bootstrapping process, the paired differences in the shuffling and residuals methods’ pAUC estimates (D_pAUC_(i) = pAUC_res_(i) – pAUC_shuf_(i)) were assumed to approximate a normal distribution; therefore, a two-tailed paired *t*-test was used to assess the null hypothesis of D_pAUC_ = 0.

### Analysis of Simulation Parameter Effects on Relative Hit and False Alarm Rates

To examine the simulation parameter effects, we focused on a fixed α_c_ = .05 threshold, and applied regression analyses to the residuals-shuffling differences in HRs (D_HR_) and FAs (D_FA_) separately. To avoid specific distributional assumptions regarding D_HR_ and D_FA_, we conducted these analyses using the same 1000 subsamples as were used for the pROC tests. We considered all simulation parameter combinations, with the exception of those for which *m* = 0 (to omit cases when D_HR_ = 0 - 0 by definition, and with this same exception applied to the D_FA_ analyses for consistency). Consequently, 450 × 1000 (or 120 × 1000) unique D_HR_ and D_FA_ estimates, respectively, contributed to the two analyses, for each dataset on which they were performed.

Separate general linear models for the two dependent variables (DVs), D_HR_ and D_FA_, were constructed using five independent variables (IVs): the mean-centered equivalents of the four directly manipulated simulation parameters (*T*, *f*_osc_, *p_base_offset_*, and *m* for most datasets, with *f_osc_* replaced with *p_base_* when *p_base_* was directly manipulated), and also the square of the mean-centered modulation strength parameter ((*m*-*m̅*)^2^), given trends in the data visualizations that were suggestive of quadratic effects of the *m* factor (Fig. 4(a-b), Fig. S2). For each DV, the full GLM consisted of the intercept term plus the sum of the main effect, 2-way interaction, and 3-way interaction terms formed from the five IVs.

**Fig. 4.**
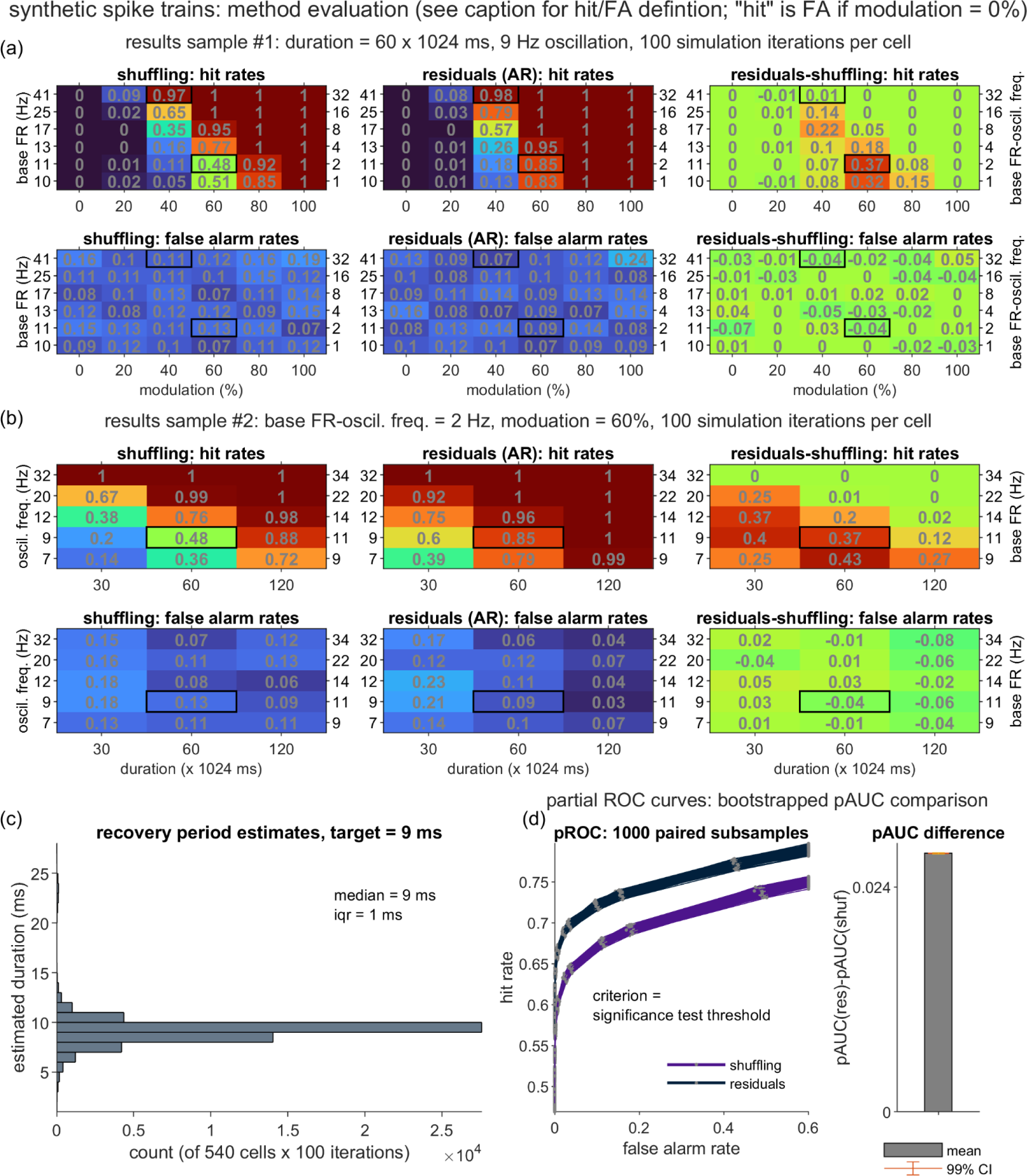
Performance of the shuffling and residuals methods over a diverse synthetic dataset. One hundred synthetic spike trains were generated for each of 540 unique parameter setting combinations (resulting from the crossing of the varied duration (*T*), modulation strength (*m*), oscillation frequency (*f_osc_*), and [base FR - oscillation frequency] offset (*p_base_offset_*) parameters; see Methods). For each spike train, corrected PSDs were generated by the two correction methods. When *m* > 0, a PSD represented a “hit” if it contained ≥ 1 point of significant power within the three nearest neighbors of the ground truth oscillation frequency (= *f*_osc_, when *m* > 0) on the sampled frequency axis (resolution = 0.9766 Hz). A false alarm (FA)-containing PSD included ≥ 1 point of significant power at frequency points beyond the 10 nearest neighbors of *f_osc_* (when *m* > 0) or at any frequency point (when *m* = 0). (a) A subset of the methods’ hit and FA rates, shown for all combinations of *m* and *p_base_offset_*, for a fixed *T* = 60 × 1024 ms and *f_osc_* = 9 Hz. Bold outlines highlight the parameter combinations used for the two example spike trains depicted in Figs. 2-3. (b) Hit and FA rates, shown for all combinations of *T* and *f_osc_*, for a fixed *p_base_offset_* = 2 Hz and *m* = 60%. Bold outlines highlight the parameter combinations used for the second, lower FR spike train depicted in Figs. 2-3. (c) Histogram of the residuals method’s estimates of the recovery period duration (ground truth *n*_r_ = 9 ms), collapsed over all of the synthetic spike trains. (d) Partial receiver operator characteristic (pROC) analysis of the residuals (res) and shuffling (shuf) methods’ performance, restricted to the 450 *m* > 0 conditions (for which a hit rate may be defined). Left: Hit versus FA rates are plotted over varying statistical test thresholds; individual curves represent spike train subsamples (matched for *res* and *shuf*) that were uniformly drawn from the 450 conditions (to obtain bootstrapped estimates of variability; see Methods). The “partial” designation refers to the truncating of individual curves to the FA range bounded by the minimum and maximum FAs observed over all curves. Individual curves may fail to reach 100% hit and FA rates due to the use of a one-sided statistical test, and may remain above 0% due to highly elevated PSD values that surpass even the most stringent *α_c_* level (1 × 10^−8^). Right: Mean and 99% CI corresponding to the *t-*test (df = 999) contrasting the residuals-shuffling partial area under the curve (pAUC) difference against zero.

### Post-Hoc Analyses of False Alarms in Spike Trains with High Firing Rates

In addition to the above ROC and regression analyses, we applied an additional set of post-hoc analyses to the synthetic dataset that had been generated with high FR settings (see “Synthetic Spike Train Test Sets” above). These follow-up assessments were motivated by the observation of unexpectedly high FA rates, especially for spike trains generated with long *T* and high *m*, and either low *f_osc_* (residuals method) or high *f_osc_* (shuffling method; see Fig. S8-S10). We report three analyses, all of which summarized over the 1000 spike train subsamples, and focused on the *T* = 120 × 1024 ms and *m* = 1.0 conditions.

The first analysis was a linear regression consisting of two IVs (*f_osc,_ p_base__*_offset_) and the DV (the subsampled D_FA_ scores). Consequently, 30 (= number of *f_osc_* × *p_base__*_offset_ combinations) × 1000 (= *n* subsamples) unique D_FA_ estimates contributed to this analysis. The regression model included both main effects and the *f_osc_* × *p_base_offset_* interaction term.

The second and third analyses entailed the calculation of descriptive statistics. These statistics were computed separately for the two correction methods. For one set of descriptive statistics, we report the mean absolute FA rate for each of the two methods and the two most extreme *f*_osc_ cases (7 Hz, 32 Hz); this value simply corresponded to the grand average of the FA rates, collapsed over the 1000 subsamples and six *p_base_offset_* settings. For the other set, we report summaries of the peak power (and associated frequencies) for the PSDs’ false alarm points, expressed as a ratio over the peak power for the hit points. For each correction method, we iterated over the 1000 subsamples of spike trains, and for each subsample, over the 30 unique *f_osc_* × *p_base_*__offset_ setting combinations. Within each subsample, each unique *f_osc_* × *p_base_*__offset_ setting combination is represented by 20 spike trains (given the constraint that *T* = 120 × 1024 ms and *m* = 1.0). For the correction method under consideration, we identified the FA and hit points present within the 20 respective corrected PSDs. For each PSD, the peak (i.e., maximum) power estimates were identified over any identified FA and hit points, respectively. For a PSD that included both FA and hit points (as was common in the *T* = 120 × 1024 ms, *m* = 1.0 cases), we computed the peak_FA_ / peak_Hit_ power ratio. These peak ratios and the associated peak FA frequencies were then averaged over the PSDs (for which they were defined), and then averaged over the 30 parameter setting combinations and 1000 subsamples.

### Experimental Data Methods

#### Subject

We analyzed data from a single rhesus monkey (monkey G, 7.1 kg, female) which was one of two animals studied in an investigation of MPTP (1-methyl-4-phenyl-1,2,3,6-tetrahydropyridine)-induced parkinsonism. Here, we selectively analyzed data from the post-MPTP state. A prior report based on the pre-MPTP data [30] provides detailed explanations of several components of the experimental protocol that are also applicable to the post-MPTP dataset; for these components, we briefly recap the main points below.

#### Task

Single-unit extracellular signals were collected during performance of a visually-cued reaching task, on which monkey G had been trained to a high degree of proficiency. Food rewards were delivered for successfully performed trials. Full task details are available in [30]. Here, the most relevant task component is the pre-reach delay period that spanned a variable duration interval (1.01 - 3.50 s for the trials included in our analyses) from trial start to go-cue onset. During this delay period, the left hand rested at a start position (a metal bar) fixed at waist height to the animal’s left. The onsets/offsets of hand contact with this start position were detected with an infrared sensor.

#### Surgery

Devices were implanted surgically to allow access to the right thalamus and globus pallidus via a parasagittal approach. The surgery was performed under sterile conditions with ketamine induction followed by Isoflurane anesthesia. Vital signs (i.e., pulse rate, blood pressure, respiration, end-tidal pCO2, and EKG) were monitored throughout the surgery to ensure a proper level of anesthesia. Analgesics and antibiotics were administered prophylactically.

On protocols.io, we have posted two step-by-step protocols relevant to the surgical procedures (craniotomy https://www.protocols.io/private/8791490F352811EEAFC70A58A9FEAC02 and head fixation post and recording chamber implantation https://www.protocols.io/private/A317A58E352711EEAFC70A58A9FEAC02).

#### MPTP Administration

The animal was rendered hemiparkinsonian by injection of MPTP into the right internal carotid artery (0.5-mg/kg, [39]). This model of parkinsonism was chosen to facilitate care of the animal and to increase the likelihood that the animal would perform the behavioral task following intoxication [40]. The MPTP administration procedure was performed under general anesthesia (1-3% Isoflurane) and prophylactic antibiotics and analgesics were administered post-surgically. The animal developed stable signs of parkinsonism contralateral to the infusion (i.e., on the left side of the body).

On protocols.io, we have posted a protocol that includes a description of the procedures for internal carotid artery administration of MPTP (https://www.protocols.io/private/7BED34B9490A11EEACE60A58A9FEAC02).

#### Data Acquisition and Spike Sorting

Regions of GPi and VLa to be targeted for single-unit recording were located using standard neurophysiologic mapping methods [30]. The extracellular spiking activity of neurons in GPi and VLa was collected using multiple glass-insulated tungsten microelectrodes (0.5–1.5MΩ, Alpha Omega Co.) or 16-contact linear probes (0.5–1.0MΩ, V-probe, Plexon Inc.). Data were amplified (4×, 2Hz–7.5kHz), digitized at 24 kHz (16-bit resolution; Tucker Davis Technologies), and saved to disk as continuous data. The stored neuronal data were high-pass filtered (Fpass: 300Hz, Matlab *firpm*) and thresholded, and candidate action potentials were sorted into clusters in principal components space (Off-line Sorter, Plexon Inc.; RRID:SCR_000012). Clusters were accepted as well-isolated single-units only if the unit’s action potentials were of a consistent shape and could be separated reliably from the waveforms of other neurons as well as from background noise throughout the period of recording. Times of spike occurrence were saved at millisecond accuracy.

On protocols.io, we have posted protocols relevant to electrophysiological recording (https://www.protocols.io/private/7CF970AD3BE111EE882A0A58A9FEAC02) and recording chamber maintenance (https://www.protocols.io/private/F56F59C63BDD11EE882A0A58A9FEAC02).

#### Task-Aligned Analysis Windows

To focus on intervals of the monkey G spike trains when oscillatory activity was expected to be high, we restricted our analyses to task windows spanning the first second of each trial. These windows sampled data from the pre-reach delay periods; such intervals of enforced hand stillness have been historically associated with elevated beta power [41]. The 1000 ms window was chosen instead of the PSD segment size (1024 ms) because a small fraction of the go-cue onsets occurred at post-start latencies in (1000, 1024] ms. This selection step jointly served a second purpose of reducing potential sources of substantial non-stationarity in the analyzed spike data (along with a firing rate screening step, described further below).

#### Unit and Trial Selection Criteria

The process of selecting units and trials to include in analysis proceeded in three stages. As a first step, we excluded units recorded from the GPi and VLa when these failed to meet basic quality checks (e.g., poor isolation during spike sorting, or highly anomalous spike waveform features) or exhibited outlier FRs (as measured over the full recording period) of > 106.48 Hz (GPi) or > 39.50 Hz (VLa). Units with FRs < 1 Hz were also excluded.

A second, trial selection step was prompted by the observation of units for which the “hold” (i.e., pre-reach delay) period-windowed data exhibited visible inhomogeneity in firing rates over trials, often appearing as blocks of trials during which spiking was either absent or very sparse, preceded or followed by blocks for which spiking was more regular. As this pattern was suggestive of considerable non-stationarity remaining in the hold period-restricted data, we implemented a procedure to detect and remove windows associated with trials with aberrant firing rates. This procedure itself entailed multiple steps:

1. Prepare an *nW* × 1 vector **S** indicating the number of spikes in each 1000 s window.
2. Remove those rows from **S** reporting < 2 spikes.
3. Use a Gaussian Mixture Model (GMM) to cluster the remaining entries of **S**, using a forward search procedure to set the number of Gaussians (*k*) parameter:

a. For each **k** in 1 to length(**S**):

i. Fit *k* Gaussians to the data in **S** (Matlab *fitgmdist*) using initial centers (*µ*), variance (*σ^2^*), and mixing proportion (*p*) settings as determined by the Variance Partitioning method [42].
ii. Record the Bayesian Information Criterion (BIC) value for this *k*. The BIC is a negative log-likelihood-based measure that is penalized by the magnitude of *k*; lower BIC values are preferable.
iii. Break out of the loop when either:

1. BIC(k) > BIC(k-1)
2. A failure to converge is reported for iteration *k*.
b. Upon exiting the loop, return the GMM with the number of Gaussians set to *k*-*1*.
4. Assign each row in **S** to one of the *k* Gaussians (hard clustering with Matlab *cluster*, which assigns the cluster labels that maximize the probability of observing the data, given the GMM parameters).
5. Find the Gaussian **G** with the maximum number of assigned windows. Retain only those windows from **S** that were assigned to **G**.
6. From the windows assigned to **G,** remove those windows for which the number of spikes *nSpk* is an outlier, given the mean and standard deviation parameters estimated for **G**. A window is defined as an outlier when |*nSpk*-*µ_g_*| > 3* *σ_g_*).

For each unit, the output of this trial (i.e., window) selection step was assumed to represent a set of 1000 ms segments for which the firing rates were relatively homogeneous. As a final step, we discarded those units for which < 30 windows survived the trial selection procedure, or for which the mean firing rate over the surviving windows was < 1 Hz.

#### Windowed Data: Uncorrected PSD

Note that Welch’s method processes time series segments independently (i.e., the final estimated PSDs are the average of the individual PSD estimates for the Hamming-windowed 1024 ms data segments). Prior to calling *pwelch*, we prepared the hold period data by demeaning and adding 24 ms of zero padding to each 1 s hold period vector, and then concatenating these 1 s vectors into a single time series. This vector was then submitted to identical PSD analysis procedures as were described for the synthetic data.

#### Windowed Data: Shuffling-corrected PSD

Similar to the synthetic dataset, many of the recorded units exhibited low FRs, such that local shuffling was not a practical option. We extended the global shuffling option from [1] (which was only defined for unbroken time series) to temporally separated data segments with the following algorithm (all indices assume indexing starts at 1):

1. Input: **dMat** (“delta matrix”), an *nD* × *T* matrix, consisting of the original *nD* delta vectors of length *T* msec (here, *T* = 1000; *nD* = number of trials).
2. Create a list, **sISI**, consisting of the concatenation of all ISIs encountered in **dMat**, sorted in descending order of size. (Note that **sISI** will effectively function like a max-oriented priority queue.)
3. Initialize an *nD* × *T* matrix **wMat** (“working matrix”) with zero entries.
4. Initialize an *nD* × 1 vector **timeLeft** with entries of the value *T*-1 (here, 999), corresponding to the size of the largest ISI that might be inserted into each row of **wMat**. (Note that 1 ms must be reserved for insertion of the first spike for the first ISI).
5. While **sISI** is not empty:

a. **isi** ← top (max duration) entry, removed from **sISI**
b. **openRows** ← indices of rows in which **timeLeft** ≥ **isi**
c. **targetRow** ← randomly chosen row from **openRows**
d. Insert **isi** into **wMat[targetRow]**:

i. If this is the first ISI entered into targetRow, place “spikes” (1 entries) at **wMat[targetRow]**[1] and **wMat[targetRow][1+isi]**
ii. Otherwise, place a spike at **wMat[targetRow][i+isi]**, where **i** is the index of the rightmost spike present in the row
e. timeLeft[targetRow] **←** timeLeft[targetRow] - isi
6. For each row in **wMat:**

a. If > 2 spikes are present in a row, randomly re-sequence the ISIs (as in the standard shuffling procedure)
b. If **timeLeft[row]** > 0: Right shift the spikes (insert a leading series of 0s) by a random integer drawn from [0,**timeLeft[row]**]
7. **shufMat** (“shuffled matrix”) ← **wMat**

To summarize, the algorithm implements global ISI shuffling by (1) initializing an empty data matrix, and a pool of ISIs to be inserted into it, (2) randomly inserting ISIs into the new matrix, proceeding from the longest to shortest ISIs, and packing the rows in a left-aligned fashion, and (3) randomly re-sequencing and right-shifting the ISIs in each row, to eliminate the structure introduced by step (2).

Each matrix of shuffled spikes was submitted to PSD analysis through an identical manner as was described for the uncorrected PSDs, and the standard division of the original PSD by the mean of the shuffling-generated PSDs was then used to produce the corrected PSD.

#### Windowed Data: Residuals-corrected PSD

Recall that, in the case of an unbroken time series, the residuals output, *r(t)*, is not defined for the first *n̂*_*r*_ ms. In the case of the windowed data, the initial *n̂*_*r*_ ms must be dropped for the residuals output corresponding to each trial. Therefore, the delta vectors corresponding to each individual trial were zero-padded by 24+*n̂*_*r*_ ms, to both account for the initial RP-dependent lag, and to match the expected 1024 ms segment size. All other aspects of the residuals method remain as described in the context of the unbroken synthetic data.

#### Reference Synthetic Data for Comparison with Bursting Units

We conducted follow-up examination of two VLa spike trains, which exhibited spiking timing patterns and corrected PSDs that were distinct from those generally observed in our synthetic datasets. To better characterize the distinctive features of these spike trains, which included suspected burst-firing behavior, we generated additional sets of reference spike trains, which did not include a bursting component, but did roughly match the two VLa units with respect to other key parameters. All simulations used the same framework as described above. For the synthetic data matched to the first VLa unit, *T* = 118 × 1024 ms, *f_osc_* = 13 Hz, and *p_base_* = 15 Hz. For the simulations matched to the second VLa unit, *T* = 51 × 1024 ms, *f_osc_* = 14 Hz, and *p_base_* = 21 Hz. For both sets of simulations, *m* = 0.6, *n_r_* = 1, and *k* = 0, and *n* = 100 synthetic spike trains were generated.

#### Population-Level Comparisons of Oscillation Detection Rates

We used three analyses to compare the counts of significant oscillations reported by the two methods, for specific frequency bands of interest. All three analyses were performed over the pooled GPi and VLa data. The first analysis compared the proportions of PSDs for which the residuals and shuffling methods reported at least one point of significant power within the 8-30 Hz alpha-beta band. Proportions were compared with McNemar’s χ^2^ test, with continuity correction (R *mcnemar.test*). The second analysis focused specifically on those PSDs for which at least one method reported an alpha-beta oscillation (as defined above), and used logistic regression (Matlab *glmfit*, logit link) to ask whether the corresponding unit’s mean FR inversely predicted residuals-specific identification of alpha-beta incidence. The dependent measure was a binary coding of the PSDs (1 = alpha-beta detected by residuals method alone; 0 = detection by either both methods or shuffling only). The third analysis used McNemar’s χ^2^ to compare the proportions of PSDs for which the residuals and shuffling methods reported at least two consecutive points of significant power in the (0,4] Hz delta range. This delta band measure was intended to serve as a substitute for more traditional forms of 1/f-like trend assessment (which involve curve fits that do not accommodate possible RP distortion [12]).

## Results

### Results Overview

In the following, we first consider the shuffling and residuals methods in the context of synthetic spike trains. We will highlight two example cases that aid in illustrating the procedures and outputs of the two correction methods, and then present the results of a systematic comparison of the methods’ performance over simulations that encompass diverse parameter settings. Subsequently, we present the shuffling- and residuals-corrected output for the empirical single-unit data collected from the parkinsonian monkey, and highlight cases in which the divergence of the two methods’ output is particularly notable.

### Synthetic Data Results

#### Shuffling Method: Examples of Detected and Non-Detected Oscillations

Stochastic spike trains were generated according to the framework described in [2, 7]; full details are provided in the Methods. To briefly recap, spike arrivals were sampled from 1 kHz rate functions, *p*_spk_(t), which could be varied with respect to the parameters that governed the exponentially rising recovery period (steepness *k*; duration *n_r_*), the oscillation (frequency *f_osc_*; modulation strength *m*), the baseline FR (*p_base_*), and the simulation duration (*T*). We considered a variety of parameter settings, although with an initial priority on relatively low *p_base_* values, given the challenges posed by cases in which oscillation detection may be particularly reliant on the information present in the ISI distribution.

Figure 2 presents two illustrative cases generated from these simulation procedures (spike train duration *T* = 60 *x* 1024 ms, *f*_osc_ = 9 Hz; for all synthetic data, *n_r_* = 9 and *k* = 0.7, unless otherwise specified). The two cases are similar in that standard statistical assessment of the uncorrected spike trains fails to detect the underlying 9 Hz oscillatory component, but differ in the factors that likely contribute to this failure, and in the shuffling method’s ability to recover this spectral feature.

Figure 2(a) shows the uncorrected spectra. As compared to the second (Fig. 2(a) *right*) example, the first spike train was generated with a relatively high base FR (41 Hz), but with a somewhat low amplitude modulation factor (*m* = 40%). As noted in the Introduction, high FRs lead to more dramatic RP-related distortions of the associated spectra [1]. Given these combined limitations (low *m*, greater distortion), the 9 Hz peak fails to cross the upper bound of the 95% corrected CI. The second spike train was generated with a stronger modulation setting (*m* = 60%), and a lower base FR (11 Hz), which results in less dramatic RP distortion; however, these advantages are offset by the reduced oscillation signal-to-noise ratio (SNR) that is also a known consequence of low FRs [3].

To both of these spike trains, we applied the standard global shuffling procedure (Fig. 2(b)), yielding estimates of the renewal-equivalent power spectra (Fig. 2(a), red lines) and the shuffling-corrected PSDs. For the high FR, low *m* example, the mean shuffled data PSD follows the RP distortion trend while avoiding the implanted oscillation. Consequently, division by this shuffling-generated PSD achieves its intended outcome: The corrected spectrum (Fig 2(c) *left*) is generally flat, apart from the 9 Hz peak, which does surpass the upper frequency-derived CI threshold.

However, such successful peak recovery is not observed for the low FR, high *m* example (Fig. 2(c) *right*). In this case, the shuffled PSD appears to capture both the distortion trend, and, to a modest extent, the 9 Hz peak. Correction reduces the relative height of this spectral feature, thereby compounding the challenges to detecting the spectral peak that were already present due to the low SNR. Even with the RP distortion removed, the 9 Hz peak falls just short of the significance threshold.

Figure 2(d), which plots the ISI PDFs for the two cases, provides a time domain perspective on these different shuffling outcomes, by highlighting the spike timing information that shuffling correction removes. In the higher FR, lower *m* case (Fig. 2(d) *left*), > 50% of the ISIs were ≤ 23 ms, thereby limiting the opportunity for the somewhat modest amplitude, 9 Hz oscillatory trend (period ≈ 111.11 ms) to substantially influence the shape of the ISI distribution. Qualitatively, the PDF resembles the form expected for a Poisson process modulated by a recovery period alone (i.e., an initial dip followed by an exponential distribution). For the lower FR, higher *m* case (Fig. 2(d) *right*), 50% of the ISIs exceeded 73 ms (≈65.7% of the 9 Hz cycle length), and > 31% were 112 ms or longer (i.e., the spiking skipped entire cycles). These long survival times afforded greater opportunity for the moderately strong oscillation term to more visibly shape the ISI distribution, with relative peaks and valleys appearing in rough correspondence to that expected for the 9 Hz rhythm.

To summarize, the lower FR/higher *m* example illustrates an instance relevant to the general caveat raised by [1]: If much of the rhythmicity of interest is present in the ISI distribution, then caution is warranted when applying the shuffling method (or the analytical alternative), as such correction will remove any oscillatory information present in the first-order ISI statistics. What our example here helps to clarify is that such scenarios emerge not only from precise pacemaker-like patterns, but also in spike trains in which a known sinusoidal drive is expressed through relatively sparse spiking. This observation motivated our interest in examining how the residuals method, which models and removes spike history effects restricted to a short history window, might perform in such scenarios.

#### Residuals Method: Procedure and Example Case Performance

Figure 3 illustrates the steps of the residuals method, and the outcomes for the two example synthetic spike trains introduced above. To reiterate, the main steps include (1) estimation of the RP duration, (2) fitting the spike train with the bounded last-spike point process model (PPM), with the bound set to the RP duration, and (3) applying spectral analysis to the residuals of the fitted PPM.

### RP Estimation Algorithm

The RP duration estimation method is illustrated in Fig. 3(a-b) (for the two example cases) and also Fig. S1 (for a spike train with no oscillation). To recap (see Methods for full details), we sought to translate the general strategy outlined by [27] into a specific algorithm that might perform with reasonable effectiveness on limited, sparse spiking data.

We specifically implemented a procedure that compares the variance explained by a series of right-shifted exponential curves fit to the ISI PDF. We pursued this approach after observing, on cases such as the Fig. 2-3 examples, that the curve goodness-of-fit often reaches its first local maximum when the exponential is anchored to the first post-RP time bin (i.e., when a unit with *m* = 0% should first exhibit ISIs consistent with Poisson spiking), even when noise and oscillatory trends create visible departures of the post-RP region from a clean exponential form. In Fig. 3(a) we highlight select iterations of the right-shifted curve fits, for the two examples. When the curves are left-anchored to the first possible start lag (1 ms), one observes visibly poor fits over the initial, RP-spanning time bins. As shown in Fig. 3(b), these mismatches correspond to relatively low values for our goodness of fit metric (the deviance difference, see Methods). Iterating through the subsequent RP-containing bins, the fit measure gradually improves, reaching local peaks (in both examples) at the first post-RP time bin of 10 ms. After detecting subsequent decreases in the exponential term’s explanatory power at 11 ms, we confirm the presence of initial local maxima, and return, for both spike trains, RP duration estimates (*n̂*_*r*_) that match the ground truth *n_r_* of 9 ms.

Note that more thorough evaluation of RP estimation performance is provided in the Methods Evaluation section, where we report on the systematic comparison of the shuffling and residuals methods.

### Bounded Last-Spike Point Process Model

The next step of the residuals method entails the estimation of the shape of the RP (i.e., the *t* − *s*_∗_(*t*) effect; see Methods) bounded within a historical interval of *n̂*_*r*_ millseconds. Fig. 3(c) illustrates the inputs and outputs of our current estimation procedure on the two example cases and with *n̂*_*r*_ = 9 ms. As in global shuffling, the input (leftmost panels, zoomed in on the first 209 ms) was the entire binary spike train vector.

As stated in the Introduction, we chose to use the point process modeling (PPM) framework to estimate the *t* − *s*_∗_(*t*) effect. In general, PPMs may operate on binary incidence vectors to estimate discrete-time approximations of conditional intensity functions (CIFs; see [8, 33] for additional background). When PPMs are used to predict future spikes on the basis of spike history, the CIF (*λ*[*t*|*θ*, *H*(*t*)]) represents the time varying probability that a spike will occur in time bin *t*, given the spiking history *H(t)* and the model parameters (ϴ). PPMs can be implemented in a variety of frameworks; we chose the well-established route of utilizing the Poisson Generalized Linear Model framework. The GLM took on the form 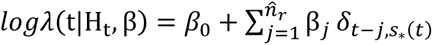, where β_0_ is the intercept term, the *δ*_*t*−*j*,*s*∗(*t*)_ terms indicate whether the most recent spike fell within the time bin corresponding to *j* ms in the past, and the *β*_*j*_ weight the additive contributions of these last-spike lags to the predicted logλ(t). Once the vector ^*β̂*^ of coefficient estimates is found, the linear expression is exponentiated to obtain the fit of the model to the original spike train (excluding the first *n̂*_*r*_ time bins for which no fit is defined). Due to the exponentiation step, the constant and the last-spike factors combine multiplicatively in the estimation of λ(t) [8].

Qualitatively, the GLM fits to the two example spike trains (Fig. 3(c), center panels) do appear to approximate the forms anticipated on the basis of the ground truth RP and *p_base_* parameters. When no spikes occurred in the prior *n̂*_*r*_ ms, the predicted λ(t) were approximately 0.041 and 0.011 for examples #1 and #2, respectively. When spikes did occur, the predicted λ(t) approximated the gradually rising RP trend.

### Spectral Analysis

Given a GLM fit, the residuals time series is generated as the actual-predicted time series (Fig. 3(c), rightmost panels; note the initial *n̂*_*r*_ time bins are filled with zero padding). The residuals are submitted to identical spectral analysis steps as are applied to the original time series. For the two example cases (Fig. 3(d)), the known 9 Hz peaks are successfully recovered, including for the lower FR spike train (Fig. 3(d), *right*) for which the shuffling method did not detect significant oscillation. This sensitivity gain – in tandem with the desired flattening of the RP distortion – is the result that we aimed to achieve by selectively removing bounded *t* − *s*_∗_(*t*) effects, and leaving intact the temporal structure expressed by the longer lags of the ISI distribution.

#### Methods Evaluation

##### Oscillation Detection Performance: Low-to-Moderate Spike Rates

So far, we have shown a pair of example spike trains, for which the residuals method either matched or exceeded the shuffling method’s detection of a known oscillation. To compare the two methods comprehensively, we ran a series of simulations and analyses that took into account a broad set of factors that may influence method performance, including the simulation parameter settings, and the corrected alpha level (α_c_) of the significance test. Evaluation began with a primary synthetic dataset that focused on low-to-moderate base firing rates (range 8-64 Hz), oscillations in the theta-low gamma ranges (7-32 Hz), and the default recovery period parameters; we also generated a series of secondary datasets to address follow-up questions.

For the primary dataset, we directly manipulated four simulation parameters (*T*, *f_osc_*, *m*, and *p_base_offset_* = *p_base_ - f_osc_*); the tables in Fig. 4(a)-(b) list the full set of parameter settings. For each unique setting combination, we generated 100 spike trains, computed the shuffling- and residuals-corrected PSDs, and tabulated the true and false positive oscillation detection rates for each (see the Fig. 4 caption for the hit rate (HR) and false alarm (FA) rate definitions).

To compare the classification accuracy of the two methods over a range of α_c_ thresholds, we conducted a pROC (partial Receiver Operating Characteristic) analysis (Fig. 4(d)). This analysis was implemented over a pooled dataset (i.e., collapsing over the parameter settings in Fig. 4(a)-(b), with the exclusion of the *m* = 0% cases). As in traditional ROC analysis, we plotted hit versus false alarm rates as the decision threshold (α_c_) was varied, with shuffling and residuals curves generated for each of 1000 subsamples of the full spike train dataset (with the subsampling performed to obtain bootstrapped estimates of variability). Since the original curves spanned variable FA and HR ranges, which did not fully span [0,1] (in part due to the use of a one-tailed significance test; see Methods), we truncated each curve to a partial ROC estimate that covered a restricted FA range (see Fig. 4 caption for details). The above steps enabled confirmation that the partial area under the ROC curve (pAUC) was reliably greater for the residuals method as compared against the shuffling method (*t*-test of pAUC_res_-pAUC_shuf_ : t(999) = 1.3048e+03, *p* << .001). This result indicates that the residuals method generally improves oscillation detection performance on this collapsed-parameter, primary synthetic dataset. To illustrate this improvement with an example, consider that at α_c_ = 0.05, the residuals and shuffling methods produced mean hit rates of 72.14% and 67.59%, respectively, and mean false alarm rates of 10.00% and 11.54%.

We have described a rationale for anticipating that the residuals method may be especially advantageous for the lowest-FR spike trains, and in particular for sensitivity to the detection of oscillations when the base FR is low relative to the frequency of the oscillation to be detected. This reasoning leads to the prediction of an inverse relationship between *p_base_offset_* and the magnitude of the residuals-associated improvement in hit rates. A qualitative trend consistent with this prediction is visible in the top right table of Fig. 4(a) (representing D_HR_ = the residuals-shuffling differences in HR, for the original, non-subsampled data). For the select simulation parameters reported in the table, the predicted inverse relationship between D_HR_ and *p_base_offset_* is apparent as a general tendency for higher D_HR_ values (and warmer heatmap colors) towards the lower rows (corresponding to lower *p*_base_offset_ settings) of the table. The Fig. 4(a) D_HR_ values also point to a general inverted U-shaped trend relating *m* to D_HR_, such that the residuals’ advantage appears to be strongest for moderate modulation strengths, as compared to the lowest *m* levels (when the oscillation is either non-existent or too weak for either method to detect) and highest *m* levels (when the oscillation is readily detected by both methods).

To quantify and statistically test these trends, and to assess the parametric effects on the methods’ relative sensitivity and specificity more broadly, we conducted separate regression analyses of D_HR_ and D_FA_ (FA_res_-FA_shuf_), with the significance threshold held fixed (α_c_ = .05), and the manipulated simulation parameters taken into consideration. As the regression analyses assume normally distributed noise in D_HR_ and D_FA_, we conducted these regressions using D_HR_ and D_FA_ values derived from the same subsampled HR and FA output as informed the pROC analysis. Fig. S2-S3 show D_HR_ and D_FA_ (summarized as the mean +/− standard error over subsamples) for all 540 unique combinations of the manipulated parameters. The D_HR_- and D_FA_-specific regression models included predictors corresponding to all four directly manipulated parameters (T,f_osc_, *p_base_offset_*, and *m*, with all *m* = 0 cases excluded), and a quadratic term for *m*.

The full regression results for the primary synthetic dataset are provided in Tables S1-S2; we highlight a subset of the significant effects here. The D_HR_ results (Table S1) statistically confirmed the predicted inverse relationship between D_HR_ and *p_base_offset_* (*p_base_offset_* term: *t* = −292.25, *p* << .001) and the observed inverted U-shaped relationship between D_HR_ and *m* (*m*^2^ term: *t* = −296.41, *p* << .001). As an additional test of the *p_base_offset_* effect – which was correlated with *p*_base_ in these simulations, thereby creating ambiguity in the interpretation of the effect as reflecting either base FR - *f_osc_* or the absolute base FR – we created a follow-up synthetic dataset in which a directly manipulated *p_base_* factor (levels = 15 or 20 Hz) was crossed with an independently varied *p*_base_offset_ factor (levels = [2:2:8] Hz) and the same *T* and *m* parameter sets as were described above. Regression analyses of the *D*_HR_ estimates from these simulations (Table S3; *D*_HR_ values are plotted in Fig. S4) replicated the expected negative *p_base_offset_* effect (*t* = −59.81, *p* << .001). This result supports the interpretation of the margin between the firing rate and to-be-detected oscillation frequency as independently influential on the residuals method’s enhanced sensitivity (beyond the effect of the absolute *p_base_*).

No specific predictions were made regarding the two methods’ relative false alarm rates, and for many of the parameter combinations represented in Fig. S3 (and Fig. S5 for the dataset with the direct *p_base_* manipulation), the D_FA_ values clustered near zero. Nevertheless, the very high *n* of subsamples considered resulted in high sensitivity to parametric effects, and significant main effects and interactions were found (Table S2; Table S4); however, their modest scale leaves uncertain the practical implications of these trends. As discussed below, more noticeable trends were found upon introducing higher FRs to the simulations.

##### Oscillation Detection Performance: High Firing Rates

The residuals method was motivated by a problem presented by sparse spike trains, but we might also ask how effectively it performs with more vigorously spiking units. To this end, we created a secondary dataset that differed from the primary dataset in its use of higher *p_base_offset_* values (Fig. S6-S8).

Three patterns stand out in the results. First, as anticipated, high FRs reduced the residuals method’s overall performance advantage: pAUC_res_-pAUC_shuf_ remained reliably positive (Fig. S6(d)), but with only a narrow margin of performance improvement.

Second, as also anticipated, this narrowed performance gap was mirrored by a similar narrowing in the residuals’ sensitivity advantage. D_HR_ values remained mostly positive but very modest (Fig. S7; Table S5).

Third, we observed a complex interaction pattern in the methods’ relative false alarm rates. Complete illustrations and statistical results are available in Fig. S8-S11 and Table S6. Here, we will highlight four principal observations (all based on quantities summarized over the 1000 subsamples; see Methods). (1) As duration *T* increased, and especially as the modulation strength *m* approached 100%, D_FA_ scores grew either very positive (for low oscillation frequencies) or negative (for high frequencies; Fig. S8). In other words, in the high *T*, high *m* regime, the residuals method switched from producing either more or fewer FAs than shuffling did, as *f_osc_* increased (*T × f_osc_ × m* interaction: *t* = −178.70, *p* << .001). (2) Although *f_osc_* is correlated with the *p_base_* parameter, the sign flip in D_FA_ scores cannot be easily attributed to increasing FRs. In the *T* = 120 × 1024 ms, *m* = 100% cases, the effect of *p*_base_offset_ on D_FA_ negatively interacted with *f_osc_* (*t_f_osc x_* _pbase_offset_ = −68.23; *p* << .001), reflecting a tendency for the sign of the FR offset effect to likewise flip. (3) Plots of the average absolute FAs for the lowest *f_osc_* setting (7 Hz, Fig. S9) show that in the high *T*, high *m* cases, the shuffling FA rates remained mostly low (mean over all subsample *x p_base_* combinations = 10.16%) in contrast to the elevated residuals FA rates (overall mean = 47.55%). For the highest *f_osc_* setting (32 Hz, Fig. S10), the high *T*, high *m* cases resulted in very elevated FA rates for shuffling (overall mean = 76.09%) and less dramatically elevated FA rates for the residuals method (overall mean = 31.73%). (4) Summarizing over all *T* = 120 × 1024 ms, *m* = 1.0 cases, both methods’ false detections were typically characterized by very low peak power, relative to that observed for hit points (mean power_FA_/power_Hit_ = 0.15 for residuals and 0.22 for shuffling) and high, gamma range frequencies (means = 83.10 Hz for residuals and 57.86 Hz for shuffling). Fig. S11 presents representative PSDs for each method, in addition to samples of the associated synthetic spike trains. Although an ideal solution would prevent the false detections, their consistent features could aid in the development of post-hoc rules for flagging suspected FA points in either method’s corrected PSDs.

##### Evaluation of Recovery Period Duration Estimation

Overall, the preceding analyses indicate that the residuals method achieves its primary objective of enhancing oscillation detection in sparse spiking scenarios, while maintaining performance comparable to that of the shuffling method when spiking is more vigorous (excluding the limited high-FA cases described above). Correspondingly, the first step of the method – the estimation of the RP duration – was also reasonably accurate. In the primary synthetic dataset, for the RP estimates collapsed across the full set of 540 × 100 spike trains (Fig. 4(c)), 51.07% identified the 9 ms *n_r_* setting precisely, 85.13% erred by ≤ 1 ms, and 94.78% erred by ≤ 2 ms. The follow-up, high FR dataset also demonstrated generally low error rates (Fig. S6(c): % of absolute errors, |*err*|, ≤ 0, 1, or 2 ms: 66.22%, 93.53%, 99.87%).

To determine whether this accurate performance might generalize to other RP durations and forms, we generated three additional secondary datasets, with the original RP parameters modified to produce either a long relative RP (*n_r_* = 18 ms, steepness parameter *k* = 0.7), a short, sharply-rising relative RP (*n_r_* = 4 ms, *k* = 0.4), or a short absolute RP (*n_r_* = 3 ms, *k* = 0). Estimates clustered closely around the ground truth *n_r_* for both the 3 ms absolute RP (Fig. S12(c): % |*err*| ≤ 0, 1, or 2 ms: 87.28%, 96.62%, 98.90%) and the 4 ms relative RP (Fig. S13(c): % |*err*| ≤ 0, 1, or 2 ms: 84.03%, 96.58%, 99.01%). When the relative RP with the default *k* = 0.7 steepness is doubled to 18 ms, absolute error rates are comparable to those for the original 9 ms duration if the deviations under consideration are doubled accordingly (Fig. S14(c): % |*err*| ≤ 0, 2, or 4 ms: 44.79%, 86.73%, 92.72%).

These datasets also afford the opportunity to ask whether the residuals method retains its performance advantage when confronted with variable RP features. As reported above for the default RP datasets, panels (a), (b), and (d) of Fig. S12-S14 show, for each RP setting and correction method, highlighted HR and FA outcomes for select parameter combinations, and the overall pROC and pAUC output. For all three RP configurations, residuals remains the better-performing oscillation detector (*t*-tests of pAUC_res_-pAUC_shuf_ : *t*s(999) > 1.0608e+03, *p*s << .001). At *α_C_* = 0.05, the difference in the mean residuals - shuffling hit rates ranged from 4.24% to 5.47%, and the difference in the mean residuals - shuffling FA rates ranged from −3.96% to −1.13%.

### Experimental Data Results

#### Motivation and Approach

Further comparison of the residuals and shuffling methods, conducted on real, biological single-unit data, served two purposes. First, any qualitative discrepancies between the neuronal and synthetic spectra provide useful information that can help inform the development of more accurate simulation frameworks in the future. Second, and most importantly for the present purposes, these evaluations can help experimentalists anticipate what might be encountered when applying either PSD correction method to their own data, and determine what further development, if any, might be needed to tailor the methods to the specific needs of real-world research questions.

In empirically-observed, neuronal spike trains, the ground truth is unknowable. That said, we prioritized sampling neural data acquired during circumstances for which the prior expectation of oscillations is high, based on evidence from sources that are distinct from spike trains. Towards that end, we evaluated the two correction methods’ output on single-unit data recorded from the internal globus pallidus (GPi) and GPi-targeted ventrolateral anterior thalamus (VLa) of a monkey with MPTP-induced parkinsonism, during the pre-reach delay (i.e., “hold”) periods of a cued reaching task. Across multiple species and motor-related brain regions, pathologically exaggerated oscillations in neural activity are a hallmark of parkinsonism, especially within the beta (12-30 Hz) and neighboring alpha (8-12 Hz) frequency bands (see [28, 29] for recent general reviews, and [43, 44] for examples specific to the thalamus and GPi, respectively). Moreover, alpha-beta oscillations are especially pronounced during periods of postural maintenance, such as the hold periods under consideration here [41]. Since the bulk of the evidence concerning these oscillations has been obtained from recordings of local field potentials (LFPs), our expectation of their occurrence in spike trains is based on a source that, while ultimately derived from the same underlying electrophysiological signal, is not affected by RP-related (and general point process-related) challenges to oscillation measurement.

The shuffling and residuals methods’ output were compared for 92 GPi units (analyzed recording duration min, median, max = 30, 80.5, 262 s) and 78 VLa units (31, 74.5, 191 s). As anticipated based on previous reports [30], firing rates for the analyzed rest periods were generally slower for the VLa than for the GPi (*p* << .001 by the Wilcoxon rank sum test; min, median, max FR for VLa = 3.09, 12.24, 41.02 Hz; for GPi = 3.13, 39.90, 114.96 Hz). Our simulation results allow us to predict the likely sensitivity of the two methods for detecting oscillations in data with durations and firing rates typical of this empirical dataset. The synthetic data hit rate results for a representative low beta oscillatory frequency (12 Hz), low base firing rate (13 Hz), and short spike train duration (30 × 1024 ms; Fig. S15) predict that, for an oscillation modulation strength of at least 60%, the residuals method at minimum will detect the oscillation > 65% of the time.

Application of the two methods to the empirical dataset revealed considerable variability in the resulting corrected PSDs, but some recurring patterns do stand out. Here, we will highlight two of these patterns with four representative units (Fig. 5, Fig S16). The full population results are summarized in Fig. 6. To highlight the (0,100] Hz region over which we sought significant power, all neural spectral plots are confined to the initial 100 Hz; however, note that full spectra may be generated using code from the Github repository that accompanies this report.

**Fig. 5.**
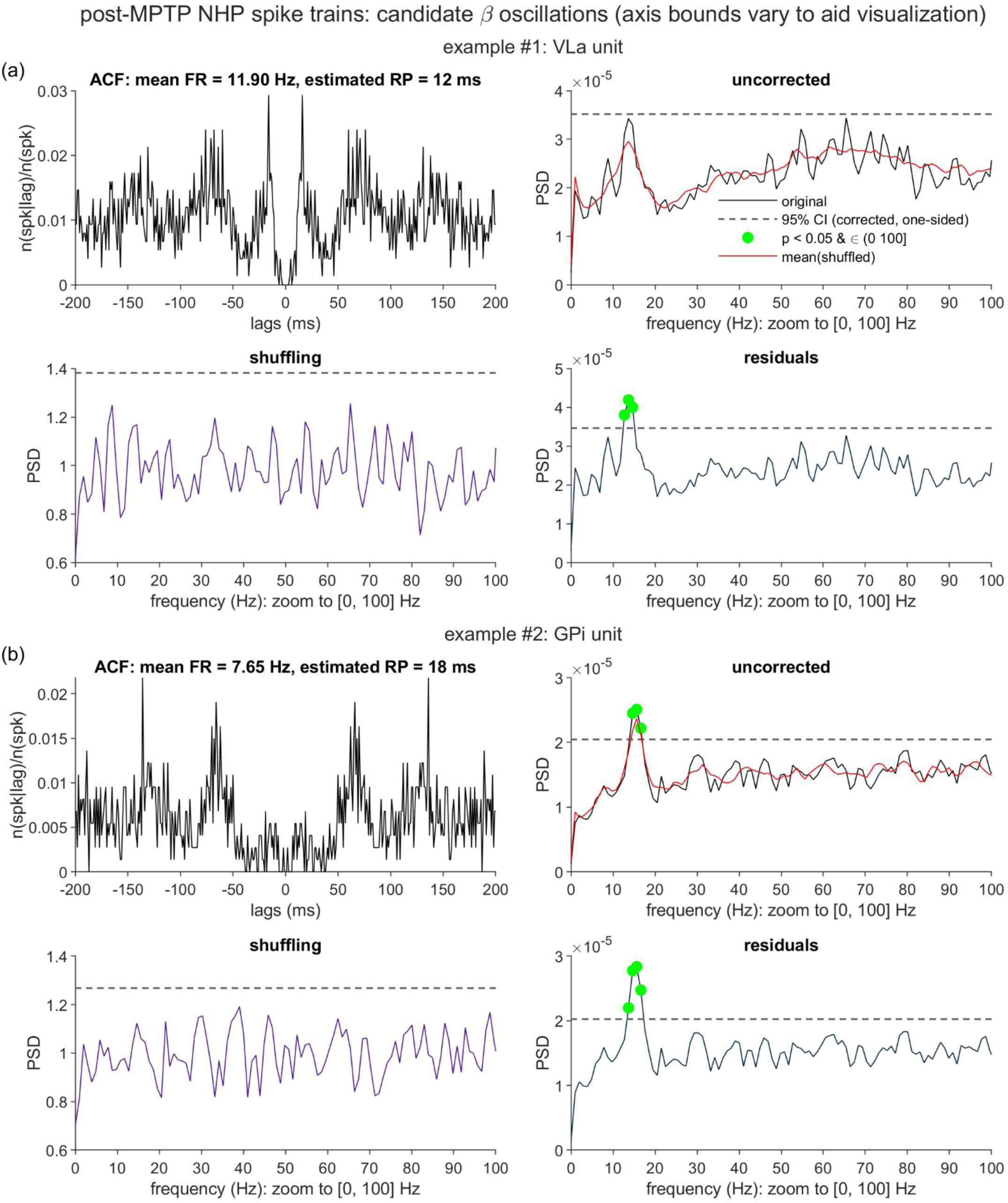
Comparison of the shuffling and residuals output for two highlighted units acquired from a parkinsonian non-human primate (NHP). Spike trains were sampled from one monkey that had been injected with the MPTP (1-methyl-4-phenyl-1,2,3,6-tetrahydropyridine) neurotoxin, and restricted to resting periods of a reaching task. Example units exhibit autocorrelation functions (ACF) consistent with a possible β (beta band; 12-30 Hz) oscillation. (a) Example unit from the ventrolateral anterior thalamus (VLa). Top, left: ACF, normalized by total spike count; spk = spike. Top right, bottom left, and bottom right: Uncorrected, shuffling-corrected, and residuals-corrected PSDs for the VLa unit, following the formatting and statistical conventions from Fig. 2-3. In the uncorrected spectrum panel, the ISI-attributed PSD (red line) was estimated via an adaptation of the global shuffling method to temporally separated task windows (see Methods). (b) The same sequence of plots as described in (a), as applied to a unit from the internal globus pallidus (GPi).

**Fig. 6.**
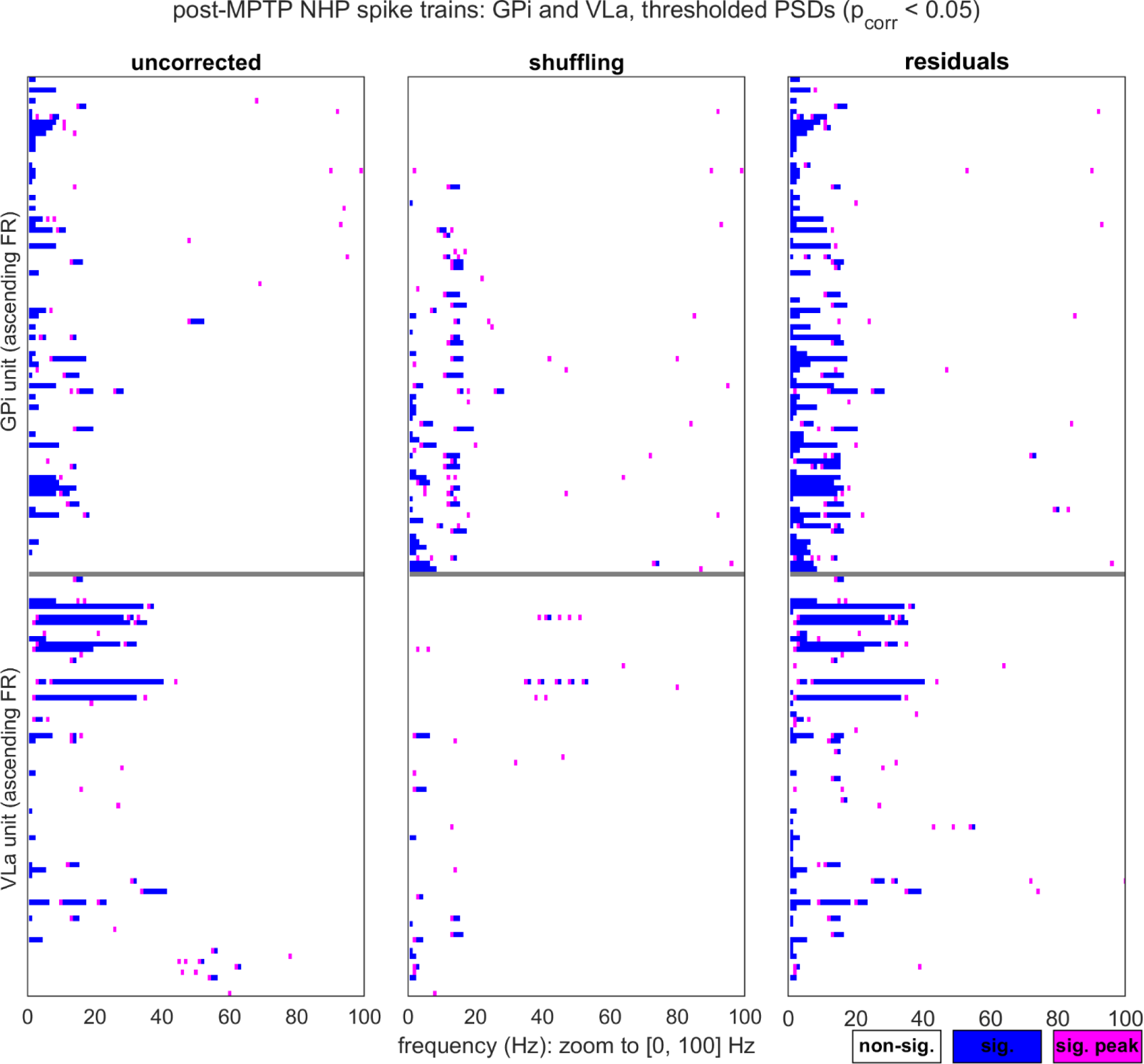
Points of significant spectral power identified by the shuffling and residuals methods across the population of analyzed spike trains acquired from the parkinsonian non-human primate (NHP). Spike data were sampled and statistically tested according to the same conventions as applied to the two example units in Fig. 5; *p_corr_* = corrected p value. Left panel: Points of statistically significant (sig.) power for the full set of GPi (top) and VLa (bottom) uncorrected power spectra. Each heatmap row represents a single power spectrum, bounded within [0, 100] Hz, with each significant point labeled as either a magenta peak (local maximum within a set of neighboring significant points) or a blue non-peak. For each region, rows are sorted in order of ascending firing rate (slowest FRs in the highest-positioned rows). Middle panel, right panel: Points of significant power found in the shuffling- and residuals-corrected PSDs.

#### Single Unit Examples

Fig. 5 presents two units that illustrate the first recurring pattern, which is characterized by its consistency with what the simulations would predict: beta (or alpha) peaks that were recovered by the residuals method exclusively, from sparse spike trains that share qualitative features with our synthetic data. Fig. 5(a) shows such an example from the VLa (mean FR = 11.90 Hz, analyzed duration = 63 s). The uncorrected PSD matches the canonical RP distortion form, with a candidate beta oscillation (peak = 13.67 Hz) sitting atop the depressed low-frequency region. The oscillation is flagged as significant in the residuals PSD, but not the uncorrected or shuffling PSD. In this case, the behavior of the shuffling method is similar to that illustrated in the right-most column of Fig. 2(a): The shuffling approximation of the renewal equivalent PSD captures not only the global distortion-consistent trend, but also much of the candidate beta peak, resulting in partial removal of this feature from the shuffling-corrected PSD. The 11.90 Hz FR indicates that, when considered relative to the candidate beta oscillation’s frequency, a spike occurred on average just .87 times per cycle. Fig. 5(b) shows a second example of this pattern, in an unusually slowly-spiking GPi unit (mean FR = 7.65 Hz, analyzed duration = 96 s, peak of the alpha-beta oscillation = 15.62 Hz).

Fig. S16 presents two VLa units that illustrate the second pattern, broadly categorized as suspected alpha-beta oscillations accompanied by temporal structure that was not incorporated into our simulations. Such additional structure included both burst firing (that is, transient intervals of elevated spike rates, see both panels (a) and (b)) and prominent 1/f-like trends (panel (b)). The two correction methods’ behavior could differ considerably in these scenarios, with variable outcomes for their relative sensitivity to likely oscillations.

In the first VLa example (Fig. S16(a)), a likely low beta oscillation (peak alpha-beta frequency = 12.70 Hz) appears in tandem with a sporadic tendency for the unit to fire in a highly stereotyped pattern of two-spike bursts. The joint presence of these two patterns is most apparent in the autocorrelation plot (top left panel). Key features of the intra- and extra-burst spiking can be discerned from the ISI PDF (Fig. S17(a)) and joint [ISI_n_,ISI_n+1_] PDF (Fig. S17(c), [45]): ISIs of 2-3 ms occur exclusively between pairs of spikes that are bracketed by much longer ISIs (≥ 14 ms), and all remaining spikes are separated by ISIs that last at least 7 ms (and often much longer). As such, the ISI statistics for this unit are consistent with a description of the spike train as switching between two different states, for which the effective recovery period varies from very short (intra-burst) to much longer (extra-burst).

This two-state characterization lends insight into the PSD outcomes. In the uncorrected spectrum, the beta peak is sunken into a typical distortion-related trough, and the shuffling method achieves satisfactory flattening of the spectrum and raising of the peak. In contrast, the residuals method performs very little distortion correction, due to the 1 ms estimated RP duration (see Fig. 17(a)-(b), *left*) and resulting 1 ms bound on the last-spike effects estimated by the PPM. Such a model should adequately account for RP effects within the two-spike bursts: On simulated, non-bursty spike trains with a 1 ms recovery period and parameters that otherwise roughly matched those for this VLa unit (see Methods), the residuals method exhibited accurate oscillation detection (for *f_osc_* = 12 Hz; *m* = 0.6: hit rate = 100%, FA rate = 7%). The apparent under-correction in this VLa example is likely due to the insufficient removal of the long RP effects that occur outside of the bursts. In the Discussion we propose potential extensions upon the residuals method that may address the need to accommodate burst and non-burst states with differing RP dynamics.

In the second VLa example (Fig. S16(b)), a likely low beta oscillation (peak alpha-beta frequency = 13.67 Hz) appears alongside a somewhat different pattern of spiking, relative to the first example. The ISIs are not split across disconnected zones of the histograms (Fig. 17(a), *right*), but the presence of bursting episodes is suggested by the high density of very short ISI incidence (28.08% of all ISIs ∊ [2,3] ms, compared to a mean of 4.83% for matched, non-bursty synthetic units; see Methods) and sequential co-occurrence (6.02% of all ISI_n_, ISI_n+1_ pairs ∊ [2,3] ms, compared to a mean of 0.25% for the matched synthetic units). Therefore, it is conceivable that this unit’s spiking behavior might also be appropriately described as switching between states of higher FRs (intra-burst) and lower FRs (extra-burst). If so, whether the RP would also be best characterized as varying between the two states is unclear.

These spike pattern observations may again aid interpretation of the PSD results, to some extent. In the uncorrected spectrum, the beta peak sits atop a prominent 1/f-like trend, to which aperiodic burst-associated FR fluctuations could be a contributor [46]. The 1/f-like feature stands out more strongly than any RP distortion that may be present, consistent with the possibility that a very short RP characterizes much of the spike train. The shuffling method does remove the 1/f-like curve, but also removes the beta peak. The residuals method again performs little correction – owing to the 1 ms estimated RP – and therefore retains both the beta peak and 1/f-like trend (with many of the elevated low-frequency points of the PSD also labeled as significant). As alluded to in the Introduction, such 1/f-like features should ideally be modeled and considered an additional component of the baseline against which the statistical significance of candidate oscillatory peaks is determined. Possible strategies for extracting the 1/f-like component – which include both the aforementioned extensions to the residuals method to model bursts, and post-hoc function fits to the corrected PSD – will also be considered further in the Discussion.

#### GPi and VLa Population Results

Fig. 6 illustrates results from the two correction methods for the whole neuronal dataset. The three columns show simplified versions of the spectra (again zoomed in to 0-100 Hz on the *x* axis), in which any points that are both significant and labeled as a local spectral peak are labeled in magenta, and all other significant points are marked as blue. Spectra are plotted according to the order of the neuron’s observed firing rate, with the lowest-FR units at the top.

The population-level patterns mirror those that we highlight with the individual unit examples. First, we found trends consistent with those that our simulations would predict, with respect to overall incidence of alpha-beta oscillation detections, and the variation of these detections with firing rates. Collapsing across the GPi and VLa units, the residuals method reported that a greater proportion of the total units exhibited significant alpha-beta power, compared to the shuffling method (66/170 versus 37/170 units, McNemar’s *χ*^2^ = 19.12, *p* < 1.3e-5). Moreover, for those units for which alpha-beta band oscillations were reported by at least one method, lower firing rates were predictive of an increased likelihood of residuals-only detection (as compared to shuffling-only or joint method detection; *β*_FR_ = - 0.089, *p* < 1.5e-5). Therefore, to the extent that the exclusively residuals-identified power did indeed correspond to true underlying alpha-beta oscillations, the method did fulfill the expectation of enhanced sensitivity, especially in the presence of low firing rates. Note also that the residuals method did identify alpha-beta oscillations over units with a variety of estimated recovery periods (median RP = 3.5 ms, iqr = 6 ms), consistent with the high hit rates seen across the varied RP parameters in the simulations (Fig. S12-S14).

At the population level, the second pattern – non-oscillatory contributors to spike timing – is most visible as a regular occurrence of 1/f-like effects. In Fig. 6, PSDs with strong 1/f-like trends tend to appear as significant points in the lowest frequency bands, starting with the (0,4] Hz delta band, and at times extending well beyond it. Using a simple rule to label likely 1/f-like trends (≥ 2 consecutive significant points within (0,4] Hz; see Methods), we identified a greater incidence of these trends in the residuals-versus shuffling-corrected PSDs (85/170 versus 25/170 units, McNemar’s *χ*^2^ = 54.40, *p* < 1.7e-13). This result is consistent with the shuffling method’s aggressive removal of the 1/f-like trend in Fig. S16(b). Also consistent with the Fig. S16B unit, we can identify additional cases in Fig. 6 for which the residuals method retains a 1/f-like trend that extends into a series of significant alpha-beta points. Therefore, with respect to the above observations of residuals-only alpha-beta detections, an open question does remain regarding the extent to which putative periodic power in these bands would remain significant after removal of the 1/f-like, presumably aperiodic spectral component.

Again consistent with the Fig. S16(b) example, we did observe additional units for which probable bursting may have contributed to 1/f-like trends. For example, for four additional VLa units that met the above delta band rule, > 25% of the ISIs fell within 2-3 ms (range: 35.76% - 42.91%), in spite of overall mean FRs of 5.35 Hz - 8.26 Hz.

## Discussion

### Discussion Overview

To recap, our principal objective was to evaluate a novel approach for removal of the recovery period’s distortion of spike train power spectra, with the ultimate aim of improving the accuracy of oscillation detection. An established ISI shuffling method [1] approaches this problem by extracting the spectral component explained by the spike train’s ISI distribution alone. Because the ISI distribution contains information about not only the recovery period, but also oscillations, the shuffling method may hinder detection of relatively subtle oscillatory spiking, especially in slowly spiking units. As an alternative, we developed a regression-based “residuals” method that also models and removes temporal structure related to the ISIs, but only those ISIs that fall within a historical bound determined by the estimated duration of the recovery period (RP) function. We compared residuals-corrected versus shuffling-corrected PSDs to determine whether the former enhanced sensitivity to oscillations, with this effect predicted to be greatest for low FR spike trains.

The following sections summarize the predicted and less anticipated aspects of our findings, propose future directions for method development, and situate the residuals method in the context of the broader set of tools available for rhythmic spiking analysis.

### Predicted Findings

The two methods were initially evaluated on synthetic spike trains, generated with varying settings for firing rate, oscillatory frequency and modulation strength, and recording duration, and with the RP represented as an exponentially-recovering modulation of spike probability. Collapsing over these variable settings, we found that residuals correction enabled more accurate classification of true versus false positive points in the power spectra. More detailed examination of the varied parameters, and of their impact on the methods’ relative hit and false alarm rates, revealed three general trends. First, and as anticipated, residuals correction primarily benefited from enhanced sensitivity to the ground truth oscillations, especially when firing rates were low, relative to the oscillation frequency. Second, two other predictors of strong relative residuals performance (moderate oscillation modulation (*m*); short spike train duration (*T*)) suggest that this method may be most useful for identifying subtle oscillatory effects that reside close to a threshold level of detectability. Third, the largely sensitivity-driven benefit was gained, for most parameter settings, without exaggeration of false positive oscillation detections.

On a limited subset of high FR, strongly oscillating spike trains, both the residuals and shuffling methods did tend to produce PSDs with very low-power FA peaks; we propose strategies for addressing these FAs farther below.

The largely favorable results for the residuals method generalized across variable recovery period durations (3, 4, 9, and 18 ms), and shapes (i.e., absolute or exponentially-rising, with variable steepness). This persistence of successful detection across RP characteristics was accompanied by relatively accurate estimation of the ground truth RP durations by the ISI distribution-fitting algorithm, with estimation errors rarely exceeding 1-2 ms.

In the real GPi and VLa data – keeping in mind the caveat concerning the unknowability of the ground truth – we did observe additional evidence consistent with our expectations. Focusing on the alpha-beta frequency range that was anticipated in the parkinsonian NHP under consideration, we found that the residuals method reported significant oscillations for a greater fraction of the units than the shuffling method did. This result was anticipated on the basis of the prevalence of low-to-moderate FRs in this dataset (especially amongst the VLa units), and follow-up tests confirmed the relevance of low spike rates for predicting residuals-only oscillation detection. Although we cannot presently determine whether the residuals-flagged oscillations are “real”, it is conceivable that future empirical work could aid in providing corroborating evidence. For example, confidence that an oscillatory drive governs a unit’s spiking could be bolstered by the discovery of matching oscillatory trends in the unit’s synaptic input dynamics (e.g., as tracked by high resolution biosensors of the relevant neurotransmitters [47]).

One might also anticipate that any significant spike oscillations may correspond to similar oscillations in a unit’s membrane voltage. Consistent with this expectation, a recent study in mice [48] observed that striatal cholinergic interneurons that exhibited robust delta rhythms in their spiking (as estimated in the time domain with an ISI-based heuristic) also showed strong delta rhythms in subthreshold membrane voltage (as estimated with wavelets).

A subset of the GPi and VLa spike trains exhibited aperiodic structure that the current implementation of the residuals method was not designed to address. Consequently, the corresponding residuals PSDs should be interpreted with care. Below, we discuss potential strategies for addressing these additional patterns that may arise in empirical spike trains.

### Areas for Future Development

In our synthetic and empirical spike trains, three special cases gave rise to unanticipated outcomes in the corrected PSDs. Below, we briefly review these cases, and the attendant directions for future development.

#### False alarms with fast spiking and strong modulation

When operating on synthetic inputs generated with high base FRs (*p_base_*) and strong oscillatory modulation (*m*), both shuffling and residuals correction produced spectra with elevated false alarm rates (Fig. S8). This trend worsened with increasing spike train duration (*T*), and with variation in the “true” oscillation frequency (*f_osc_*). The nature of the *f_osc_* effect depended on the method in question. For the lowest frequencies (e.g., 7 Hz; Fig. S9), residuals correction commonly yielded false positives, but the shuffling FA rate remained mostly low. As *f_osc_* approached the highest tested settings (e.g., 32 Hz; Fig. S10), shuffling FAs become extremely frequent, while the residuals FA rate moderated somewhat, such that the relative specificity of the two methods flipped. Under the FA-promoting conditions, both methods reported false peaks that tended to fall within the gamma band, and were characterized by very low power that barely exceeded the statistical significance threshold (Fig. S11(a)). The distinctive features of these false alarms suggest that a simple PSD postprocessing step (e.g., imposition of a more stringent significance and/or effect size threshold) may help minimize their appearance in future analyses.

Future development of the residuals method should seek to prevent these elevated false alarms. In the fast spiking, strong modulation cases, a key challenge concerns the heightened correlation between the effect that we are attempting to model (the recovery period) and the unmodeled oscillation effect. This correlation engenders an “omitted variable bias” [49] in RP function estimation. Visualization of representative spike trains (Fig. S11(b)) illustrates the core difficulty: For a strongly oscillating spike train, the spikes and the recovery period regressors will principally cluster under the positive phases of the oscillation cycle. Consequently, the RP regressors are associated not only with the recent occurrence of single spikes, but also with an elevation of the background firing rate above its overall mean. The result is an underestimation of the RP’s suppressive effect, which in turn leads to incomplete removal of the RP distortion effect from the corrected PSD.

Stevenson [49] noted that such biases might be reduced through introduction of variables that may capture suspected unmodeled effects (here, oscillations of unknown frequency), but also acknowledged that such strategies can be assumption-intensive and nontrivial. Such variables are present in those PPMs that estimate spike rhythms entirely in the time domain (e.g., [24–26]). We recap the pros and cons of these time domain methods in the concluding section below (“Conclusions: Tools for Spike Oscillation Analysis.”).

#### Burst firing impacts on the corrected spectra

Spike “bursts” consist of transient intervals of rapid spiking, occurring against a background of a slower-rate steady state [50–52]. In the empirical data, we observed multiple units that displayed likely bursting, as inferred from their ISI statistics (e.g., as in Fig. S16-S17). Previous investigations have reported bursting in the parkinsonian GPi and VLa (see [53] for a review).

Burst firing can introduce two variations in the spike train that may influence the form of the residuals-corrected PSD. First, by definition, bursting creates increases in the background FR, which may occur aperiodically. Our current residuals implementation does not remove such variation, which may in turn contribute to 1/f-like trends in the resulting PSD (consistent with Fig. S17(b); see also [46]). Second, bursting could alter the recovery period, as suggested by the Fig. S17(a) VLa example. Our current model cannot accommodate distinct burst and non-burst recovery periods, thereby preventing the accurate removal of all RP-associated variance. In the VLa example, the model estimated and removed the short, within-burst RP, resulting in insufficient correction for the long, between-bursts RP.

Future development could conceivably expand the current residuals method to capture burst- and non-burst states. Previous work has proposed discrete state-space models designed to infer burst states in point process time series [54, 55] and could inform such an approach. Armed with an algorithm that labels spike train timepoints with their likelihood of pertaining to burst or non-burst intervals, one could attempt to model the two states’ base FRs and RPs with separate sets of delta functions, with these functions weighted by the inferred probabilities that either state is active at each timepoint (i.e., 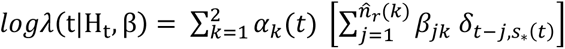, where *k* indexes over the states, and the **⍺*_k_*(*t*) denote the complementary state probabilities). Note that adoption of this strategy would require *a priori* estimates of the two states’ RP durations (*n̂*_*r*_(*k*)), and therefore the development of a dual-RP estimation algorithm.

#### Aperiodic spectral components

Several GPi and VLa units exhibited prominent 1/f-like trends in both the uncorrected and residuals spectra. “1/f-like” refers to spectral components that are well-approximated by a 1/f^χ^ function (where *χ* is the “aperiodic exponent”), to which researchers commonly add a “knee” parameter to accommodate a potential bend in the curve [56]. Strong 1/f-like effects are ubiquitous in neural activity spectra (and especially EEG/LFP PSDs) and represent the frequency domain manifestation of aperiodic trends. Removal of these trends will be necessary for rigorous interpretation of the suprathreshold points (e.g., the green points in Fig. S16(b)) in the residuals-corrected PSDs.

Two general strategies may help achieve this removal. First, as noted above, aperiodic burst spiking may contribute to the 1/f-like component [46]. Therefore, effective removal of burst-related variance at the regression stage could ultimately attenuate some 1/f-like effects. However, bursts represent just one of the many diverse underlying dynamics that may give rise to 1/f-like trends [57]. Therefore, complementary procedures will still be needed to remove any remaining aperiodic effects.

The second strategy entails the standard postprocessing step of removing a fitted 1/f^χ^ function (with a possible knee component) from the PSD. This step would assume that residuals correction has removed sufficient RP-related distortion from the spectrum. The routine introduced by [12] offers one option for achieving a robust curve fit to the PSD. This method additionally fits a series of Gaussians to any narrowband spectral peaks, and those Gaussians that sufficiently exceed the aperiodic baseline are inferred to represent likely oscillations.

### Adaptation to Time-Frequency Analysis

In its present form, the residuals method assumes the presence of stable oscillatory components. Adaptation of the method to the analysis of time-varying oscillations may be feasible, depending on two factors: the unit’s spike rate, and the stability of its RP function. Vigorous spiking and a stable RP present the optimal scenario. In this case, one could reasonably apply the PPM strategy that we described here, and submit the residuals time series to the preferred time-frequency analysis method (e.g., short-time Fourier Transform (STFT), Wavelet Transform, or bandpass filtering followed by the Hilbert Transform).

If spiking is slow, then insufficient data may be available to detect transient oscillations, even with adequate RP correction. Simulations may aid in estimating the minimal spike rate required for sensitivity to any hypothesized effects.

If the RP function is likely to vary over time, the pattern of the suspected variation should inform the adaptation approach. If one assumes that the RP varies across transitions between latent discrete states, but remains stable within states, then it may be reasonable to first apply a multi-state PPM (as we have described for bursting) and then submit the residuals to time-frequency analysis. Alternatively, if one assumes both fast spiking and RP stability within each segment of an STFT analysis, one might conceivably execute the full residuals routine (RP estimation/PPM/PSD) on each segment independently. More challenging data features (e.g., complicated RP variation patterns) will require the development of methods that go beyond the basic approaches that we have laid out here.

If time-frequency analyses are attempted, appropriate postprocessing should be applied to extract aperiodic trends. For example, Wilson et al. [58] recently proposed a time-resolved method for aperiodic and periodic trend decomposition, which extends upon the stationarity-assuming method of Donoghue et al. [12].

### Conclusions: Tools for Spike Oscillation Analysis

Throughout this report, we have referenced a set of available tools that both estimate spike oscillations and control for recovery period effects. Here, we briefly summarize the major features of each method, with an eye to the tradeoffs to consider when deciding which approach to adopt for a given project. We divide these tools into those that operate entirely in the time domain, and those that return outputs in the frequency domain.

#### Time domain methods

We further split the time domain methods into two subcategories. The first subcategory encompasses all Poisson GLM-based PPMs that may capture both recovery period and oscillation effects (e.g., [25, 26]). As described in the Methods (“Clarifications and Relationship to Pre-existing PPM Approaches”), these PPMs typically model spike history effects with a series of indicator functions of varying width. A set of narrow, short-lag regressors estimate recovery period and bursting effects, and wider, longer-lag regressors are positioned to estimate periodic effects. For example, spike counts summarized over the −50 to −40 ms bin may be used to estimate 20-25 Hz rhythms [25]. Note that the fundamental differences between such methods and the current residuals implementation are relatively modest. The principal distinctions concern the choice of indicator regressors to include, and the estimation of oscillatory trends through either strategically spaced regressors or export of the residuals to spectral analysis.

Depending on a researcher’s goals, the replacement of the PSD with binned history effects may stand out as a limitation of these alternative GLM methods. The bin-based oscillation estimates sacrifice the sharp frequency resolution of a standard DFT-produced spectrum, and represent a departure from the typical basis set of sinusoids. These characteristics may complicate projects that aim to draw comparisons with previous spectral data, or derive precise oscillation frequency information. However, when these limitations are acceptable, these GLM methods can offer some advantages. For example, by incorporating oscillation terms alongside the short-lag regressors, these models may mitigate the omitted variable bias in RP estimation.

The second subcategory consists of the Latent Oscillatory Spike Train (LOST) model of [24]. LOST is a distinct, Bayesian PPM, which shares some key features with the regression-based models, but also differs in significant ways. The shared features include short-lag history terms (implemented through splines) and other covariate regressors (e.g., related to task events) that could be easily incorporated into a standard GLM. The core difference entails the modeling of oscillations. LOST represents underlying oscillatory drives as continuous latent states, which are constrained by priors, but may vary over time with respect to their precise modulation strengths and center frequencies. These latent oscillatory components in turn shape the ongoing spike probability. This approach therefore offers a number of benefits, especially for those users who seek a principled accounting for modest fluctuations in the properties of the rhythmic drive. Moreover, as with the above-mentioned GLM methods, LOST’s simultaneous estimation of short-lag and oscillatory trends may reduce any omitted variable bias effects. Note that LOST does constrain the number of distinct oscillatory effects that the user can simultaneously estimate, and may be best suited for small datasets (due to both runtime demands and manual fine-tuning requirements).

#### Frequency domain methods

The shuffling and residuals methods form the frequency domain category. These methods offer the advantage of a corrected power spectrum estimate.

We reiterate that the shuffling method remains a reasonable approach for RP distortion correction in moderate-to-high-FR spike trains. In addition, shuffling appeared to handle some instances of aperiodic structure (e.g., 1/f-like trends, and the Fig. S16(a) bursting example) more effectively than the current residuals implementation did. The principal tradeoff for these strengths is reduced sensitivity to oscillations in sparse spike trains. Note also that the shuffling method’s run time may grow very long as the spike train duration and count of shuffling iterations increases, and that this method shares the residuals method’s vulnerability to biases introduced by strong oscillations (as indicated by Fig. S10-S11).

The residuals method is an especially useful option when a power spectrum is required, spike rates are low, and when other factors, such as modest modulation strength and short recording duration, challenge oscillation detection. In the implementation we present here, the method draws upon an accessible and flexible GLM framework. Moreover, the generation of a corrected time series output – and not only corrected oscillation estimates – expands the set of analyses that the underlying framework might ultimately support, including common time-frequency analyses.

## Acknowledgments

We thank Lisa Nieman-Vento for her contributions to animal care.

## Data Availability Statement

Both the experimentally-acquired single-unit data and the analyzed simulation outputs are available on Zenodo at https://doi.org/10.5281/zenodo.8313070. Code sufficient to reproduce all figures and tables, and the simulations and analyses that informed them, is available on Github: https://github.com/kc13/residuals_spectral_analysis. The most recent Github release (1.1.0) has been issued the DOI 10.5281/zenodo.10867519 by Zenodo. The Github repository also stores copies of the single-unit data, and synthetic datasets that are represented in a postprocessed state.

## Supporting Information

### Supporting Figures

**Fig. S1.**
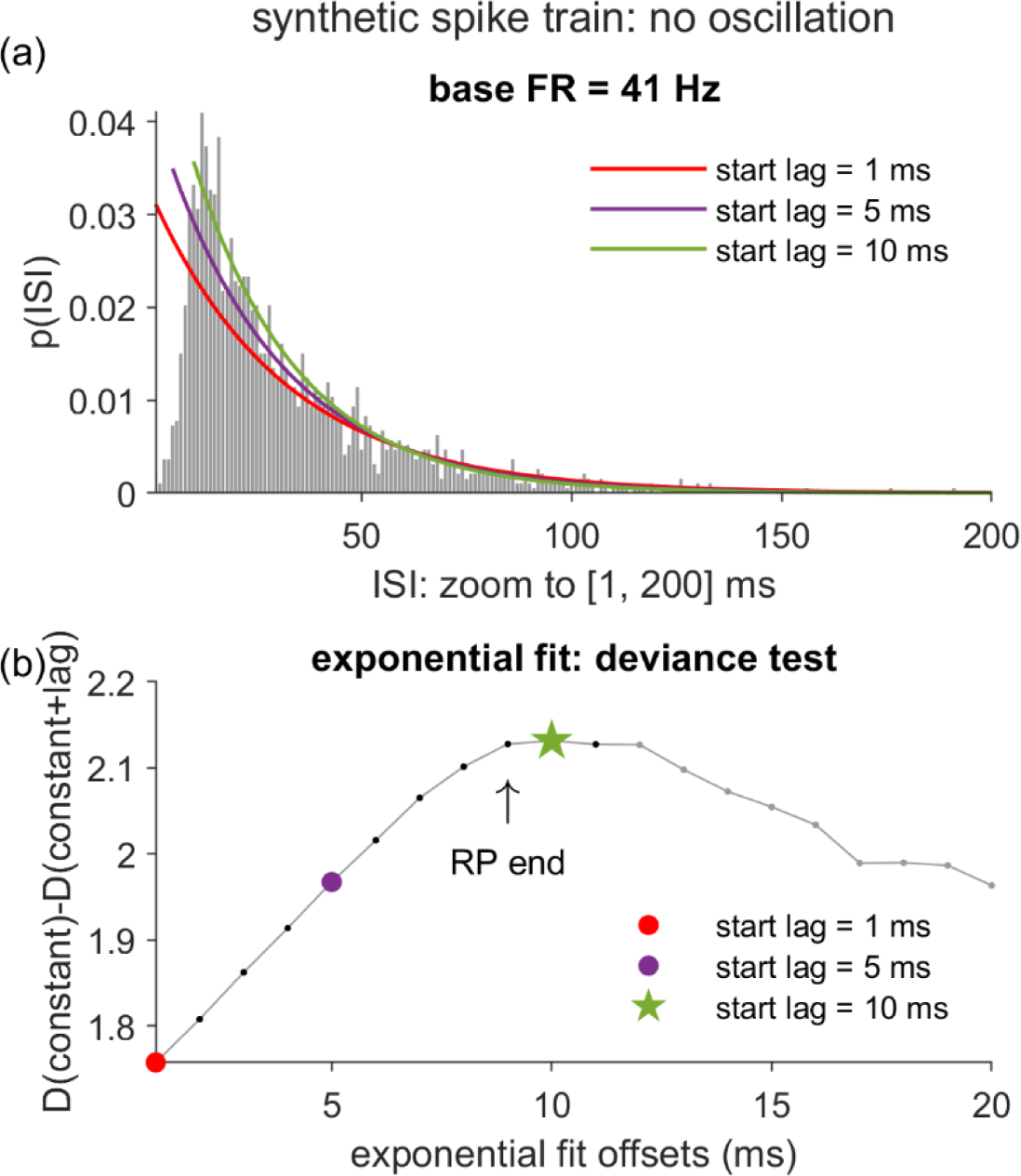
Estimation of recovery period duration: No oscillation example. (a) Illustration of the procedure for obtaining an estimate of the RP duration (*n̂*_*r*_), as applied to a synthetic spike train with no oscillation (modulation strength *m* = 0). A series of right-shifted exponential curves are fit to the ISI distribution, left-anchored to starting positions advanced in 1 ms steps (with 3 sample iterations highlighted in the figure). (b) Plot of the deviance difference statistic, ΔD, as a function of the first 20 starting positions of the exponential fits. D(constant), D(constant+lag) = deviance measures for the intercept-only and intercept+exponential curve models, respectively. ΔD tracks the goodness of fit contributed by the exponential curve. The *n̂*_*r*_ estimate is set equal to the post-spike lag immediately preceding the first local maximum in the ΔD plot.

**Fig. S2.**
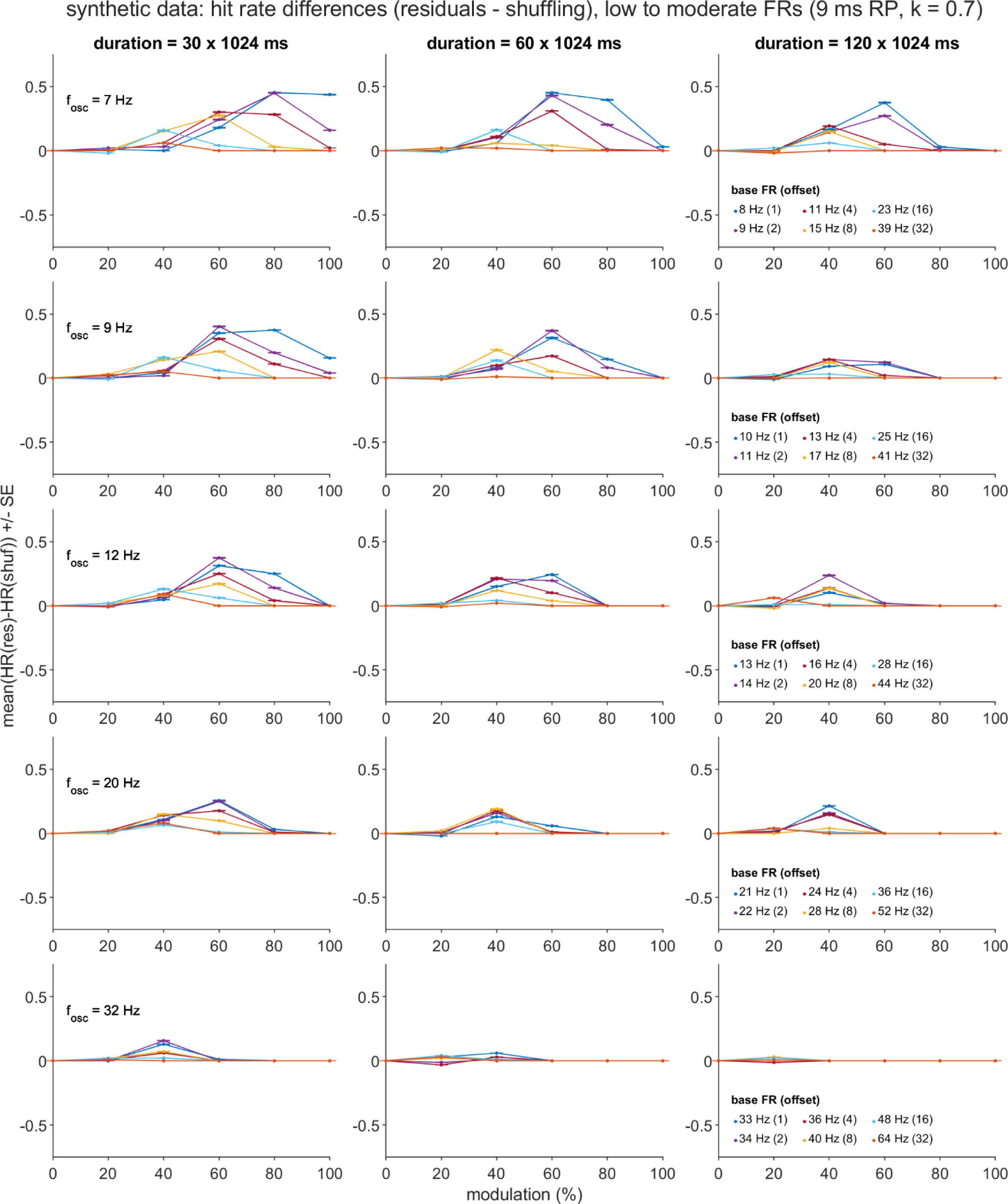
Residuals (res) - shuffling (shuf) difference in hit rates (D_HR_) over the varied parameters of the primary synthetic dataset. Means and standard errors (SE) reflect summaries over the hit rates (HR) computed for each of 1000 subsamples of the original, primary synthetic dataset, which was generated using low-to-moderate firing rates (FR) and a 9 ms relative recovery period (RP; *k* = steepness parameter). See Methods and the Figure 4 caption for details regarding the definition of a hit and the subsampling procedure. Plots depict all 540 unique combinations of oscillation frequency (*f_osc_*, rows), simulation duration (*T*, columns), oscillation modulation strength (*m*, *x* axes) and base FR - oscillation frequency offset (*p_base_offset_*, lines).

**Fig. S3.**
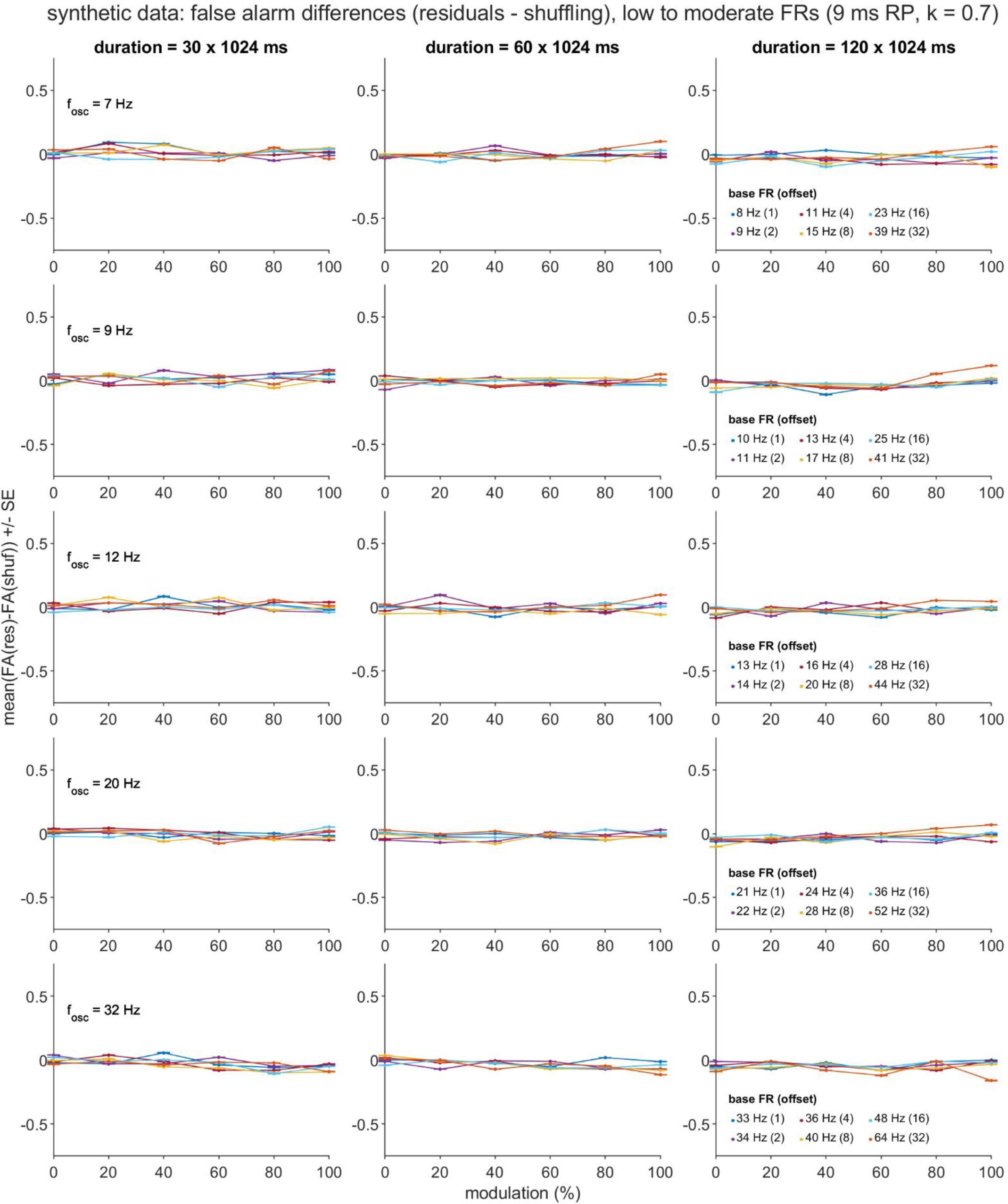
Residuals - shuffling difference in false alarm rates (D_FA_) over the varied parameters of the primary synthetic dataset. Means and standard errors reflect summaries over the false alarm rates (FA) computed for each of 1000 subsamples of the original dataset. See the Figure 4 caption for details regarding the definition of a false alarm. Abbreviations, plotting conventions, and the subsampling procedure follow from those described for Fig. S2 and Fig. 4.

**Fig. S4.**
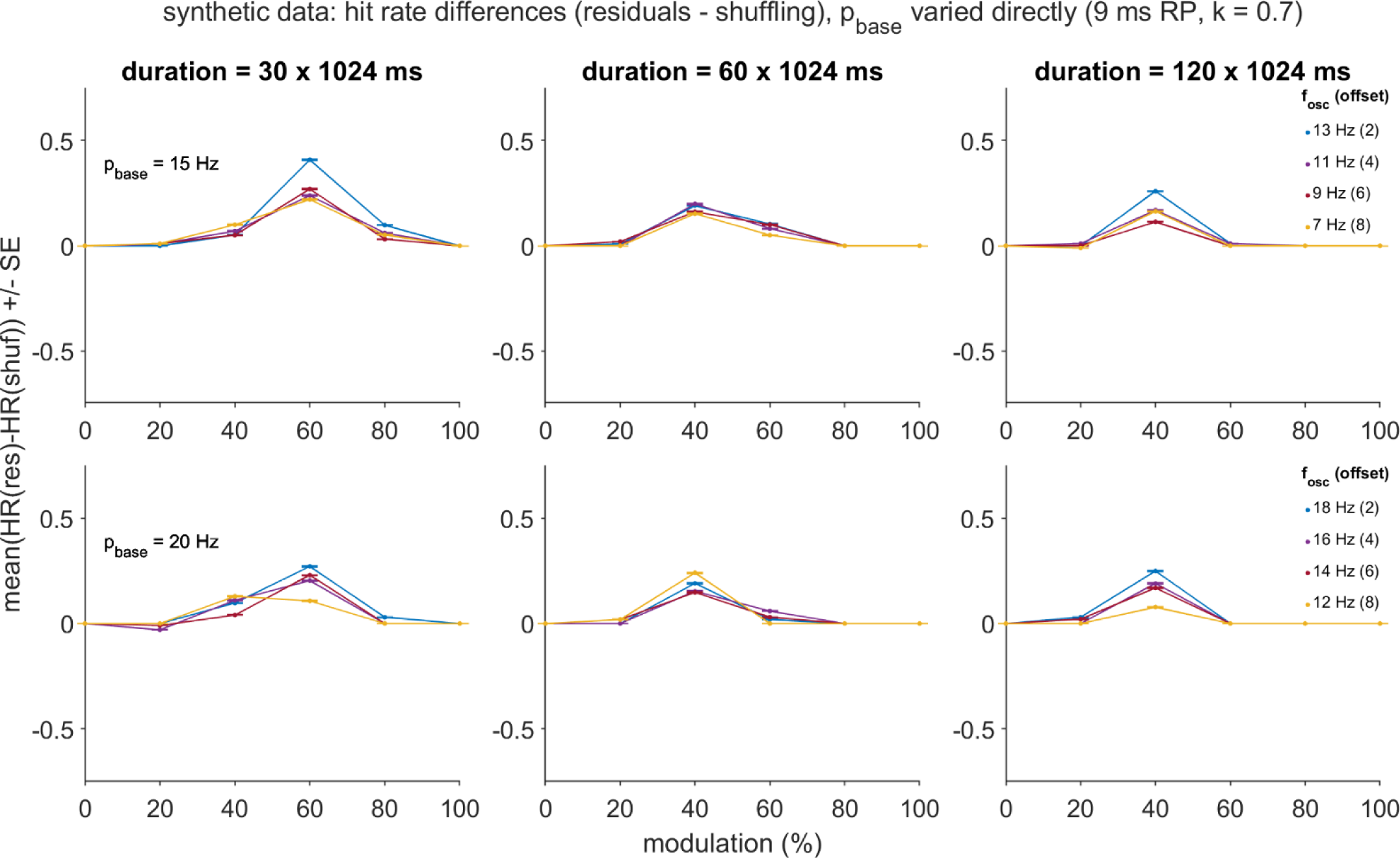
Residuals - shuffling difference in hit rates over the varied parameters of the dataset formed with direct base firing rate manipulation. Means and standard errors reflect summaries over the hit rates computed for each of 1000 subsamples of a dataset in which the *p_base_* (i.e., base FR) and *p_base_offset_* parameters were varied directly (as opposed to *p_base_offset_* and *f_osc_*). Plots depicts all 144 unique combinations of *p_base_* (rows), simulation duration (*T*, columns), oscillation modulation strength (*m*, *x* axes), and oscillation frequency (*f_osc_*, lines). All other abbreviations follow from those described for Fig. S2. See the Methods for a description of the subsampling procedure for this dataset.

**Fig. S5.**
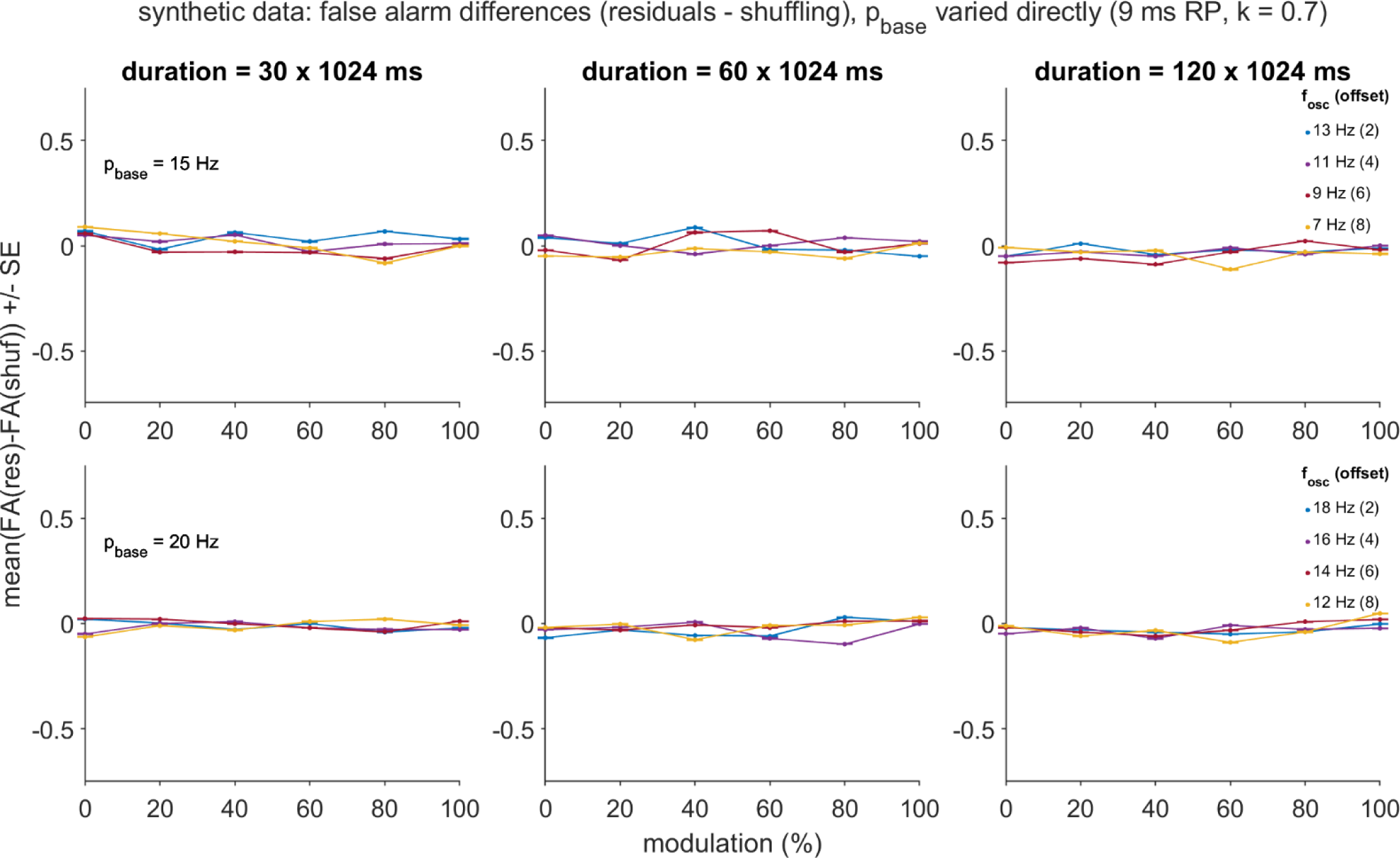
Residuals - shuffling difference in false alarm rates over the varied parameters of the dataset formed with direct base firing rate manipulation. Means and standard errors reflect summaries over the false alarm rates computed for each of 1000 subsamples of a dataset in which the *p_base_* and *p_base_offset_* parameters were varied directly. All other abbreviations, the plotting conventions, and the subsampling procedure follow from those described for Fig. S3-S4.

**Fig. S6.**
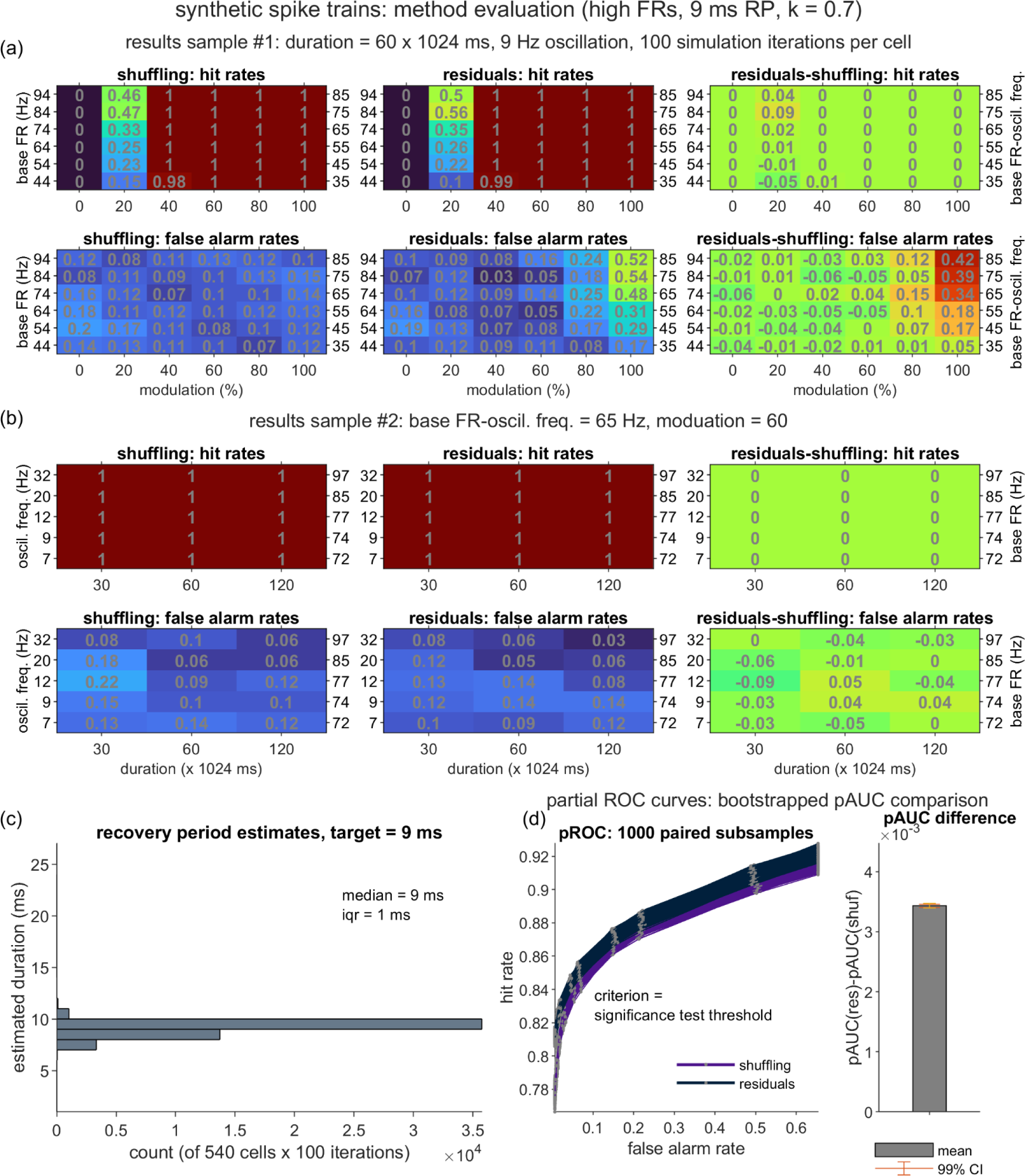
Performance of the shuffling and residuals methods over a synthetic dataset of high firing rate (FR) spike trains. Panels (a)-(d): Plotting conventions, hit and false alarm definitions, and analysis procedures are identical to those described for the primary dataset depicted in Fig. 4. Relative to the primary dataset, this high FR dataset differed in the use of greater p_base_offset_ values (see “base FR - oscil. freq.” tick labels in Panel (a)).

**Fig. S7.**
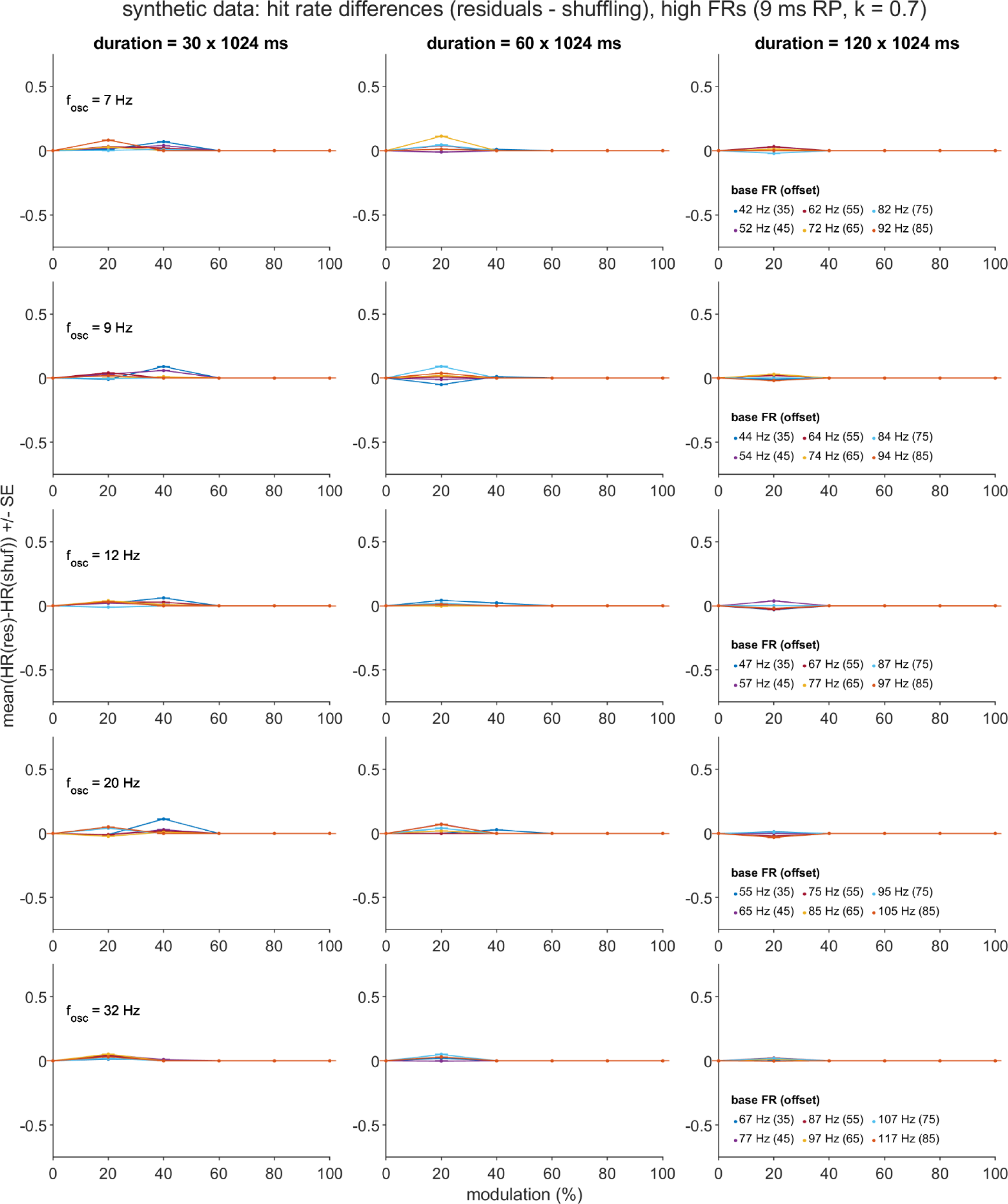
Residuals - shuffling difference in hit rates over the varied parameters of the dataset of high firing rate spike trains. Means and standard errors reflect summaries over the hit rates computed for each of 1000 subsamples of the original, high FR dataset (see Fig. S6 for details). Abbreviations and plotting conventions follow from those described for Fig. S2.

**Fig. S8.**
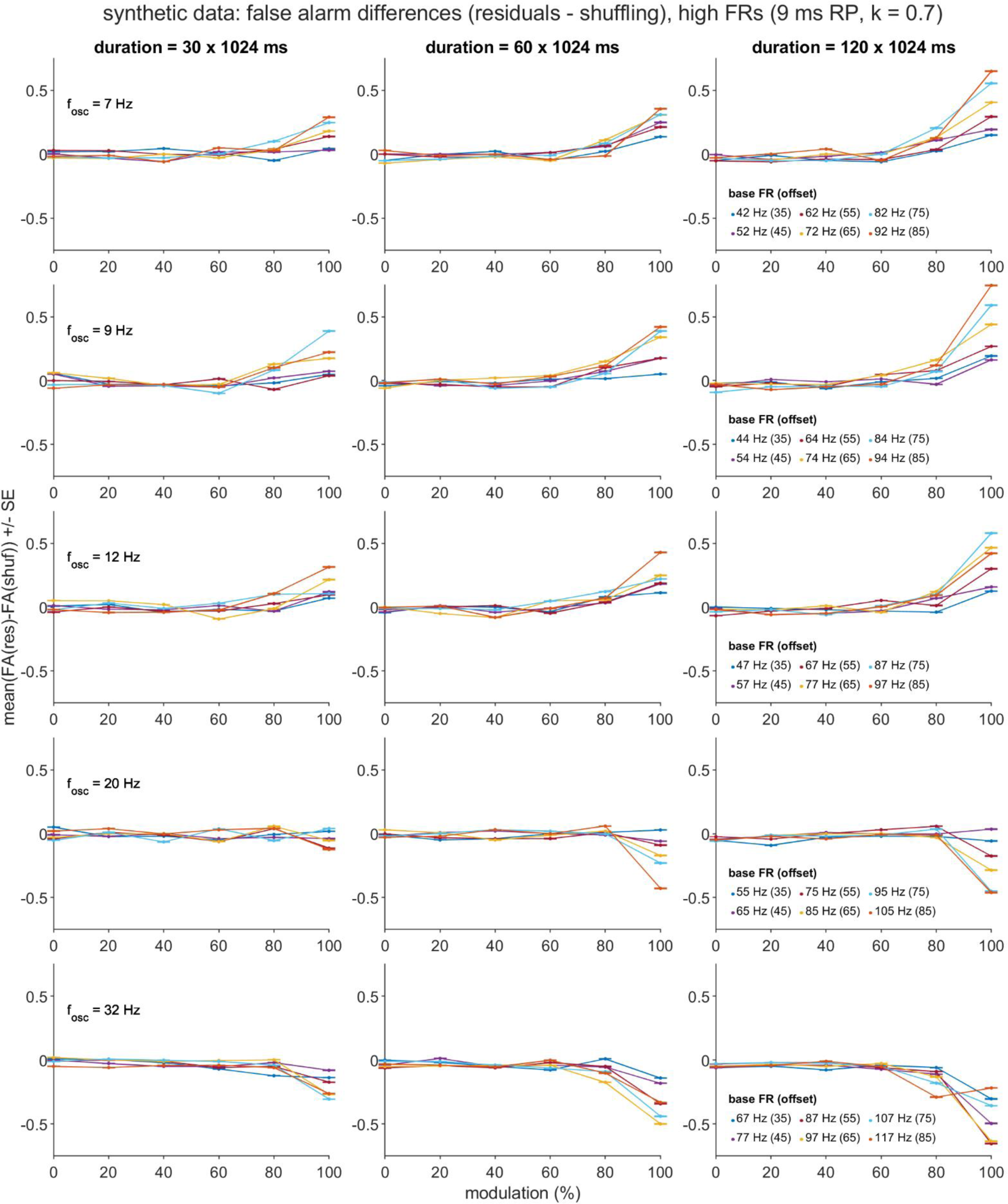
Residuals - shuffling difference in false alarms over the varied parameters of the dataset of high firing rate spike trains. Means and standard errors reflect summaries over the false alarm rates computed for each of 1000 subsamples of the original, high FR dataset (see Fig. S6 for details). Abbreviations and plotting conventions follow from those described for Fig. S3.

**Fig. S9.**
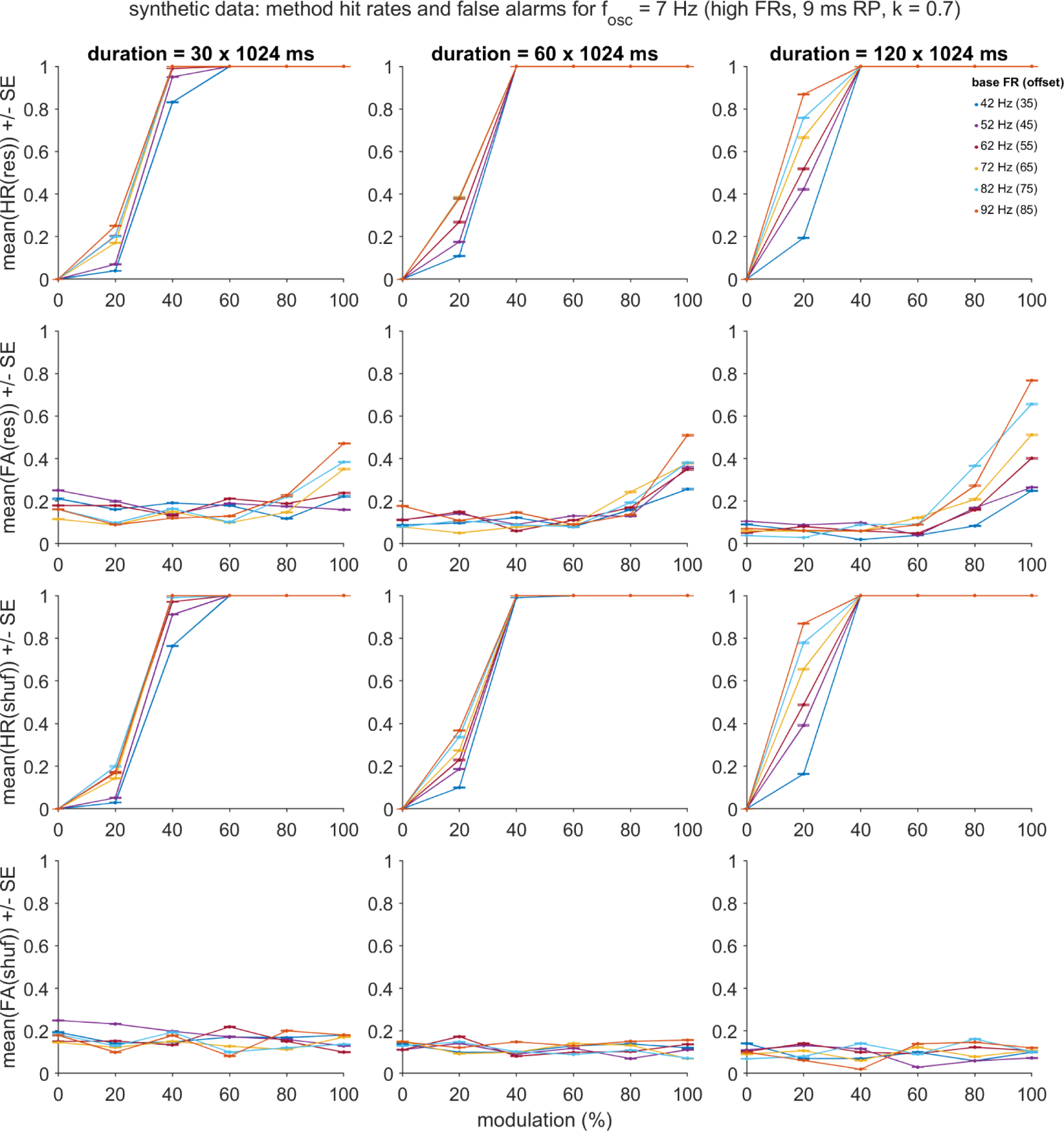
Residuals and shuffling hit and false alarm rates for the dataset of high firing rate spike trains, with the oscillation frequency (*f_osc_*) fixed at 7 Hz. Means and standard errors reflect summaries over the hit and false alarm rates computed for each of 1000 subsamples of the original, high FR dataset. Plots depict all 108 unique combinations of simulation duration (*T*, columns), oscillation modulation strength (*m*, *x* axes) and base FR - oscillation frequency offset (*p_base_offset_*, lines) for the cases in which *f_osc_* = 7 Hz. Abbreviations follow from those described for Figs. S2-S3.

**Fig. S10.**
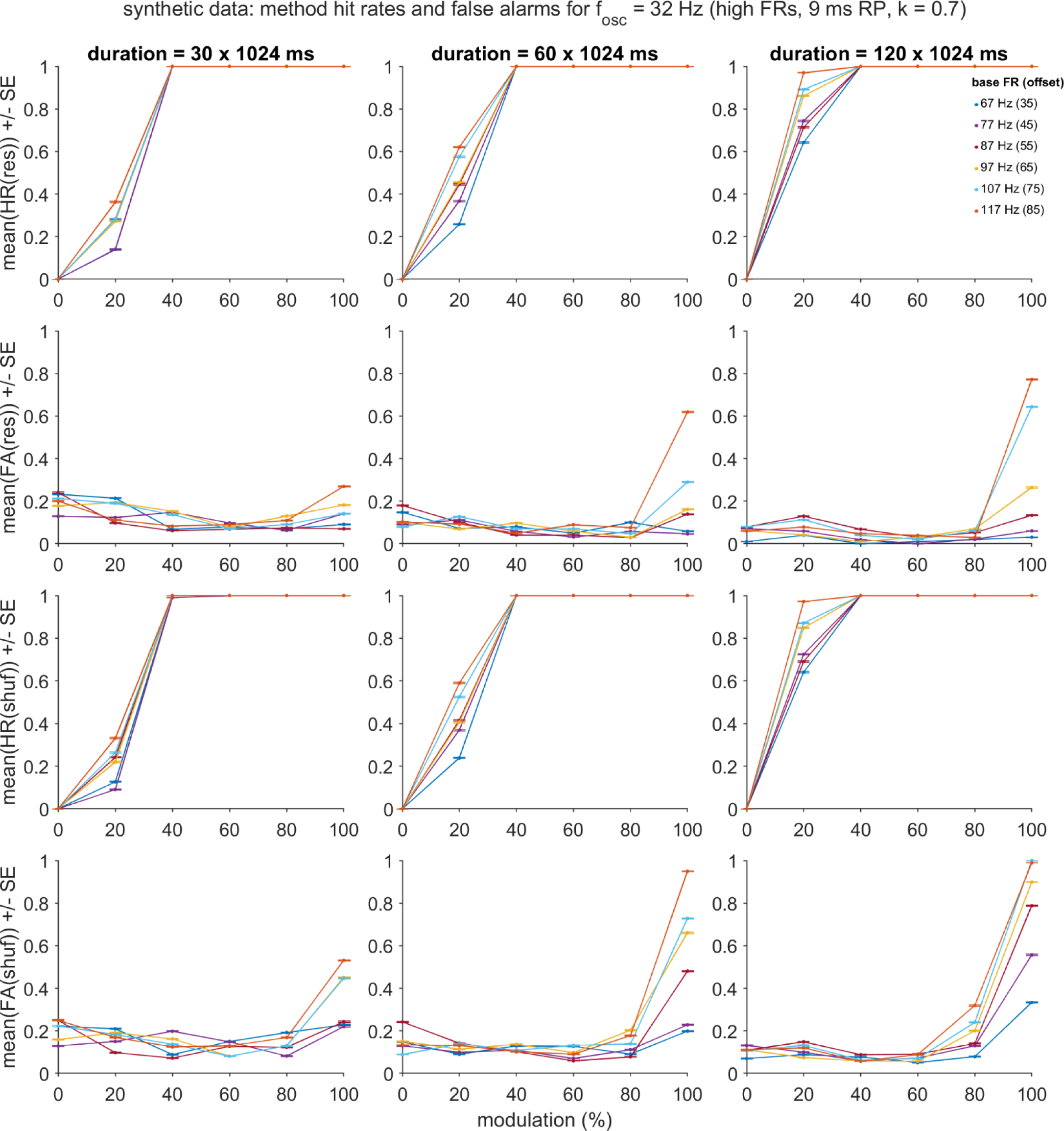
Residuals and shuffling hit and false alarm rates for the dataset of high firing rate spike trains, with the oscillation frequency (*f_osc_*) fixed at 32 Hz. Abbreviations and plotting conventions follow from those described for Fig. S2-S3 and S9.

**Fig. S11.**
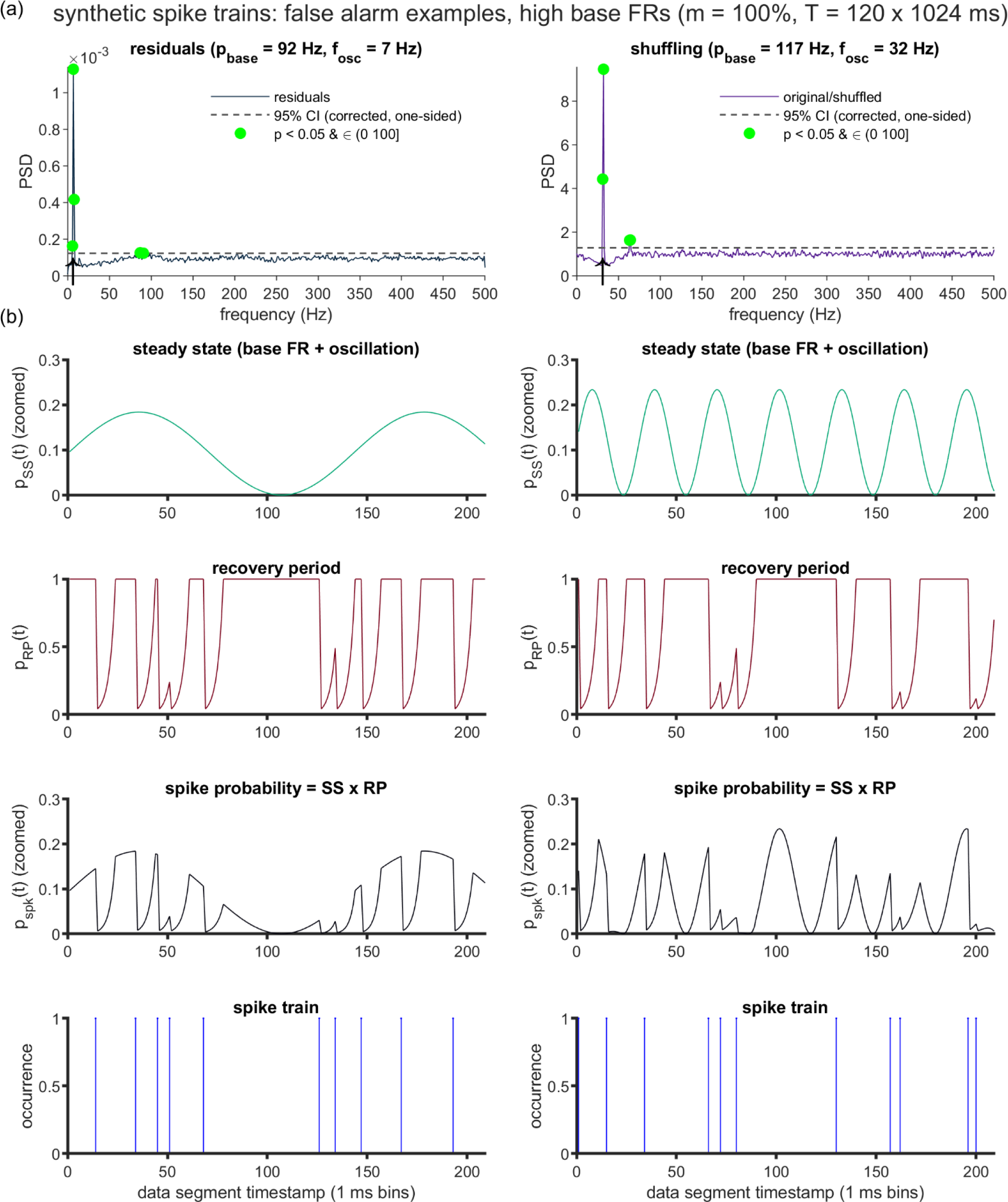
Example power spectra and spike train segments for high firing rate cases that produced pronounced false alarms following residuals or shuffling correction. Two synthetic spike trains were generated to illustrate conditions that yielded especially high false alarm rates under residuals and shuffling correction, respectively. Both spike trains utilized the maximal modulation strength (*m* = 1.0), duration (*T* = 120 × 1024 ms), and base FR - oscillation frequency offset (*p_base_offset_* = 85 Hz) settings from the original high FR dataset, and the default RP parameters (duration *n_r_* = 9 ms, steepness *k* = 0.7). Oscillation frequency was set at either the lowest setting from the full high FR dataset (7 Hz, to illustrate residuals FAs) or the highest setting (32 Hz, to illustrate shuffling FAs). (a) Corrected power spectral density (PSD) functions generated by the residuals method (left) and shuffling method (right). Statistical testing and plotting conventions follow from those described for Fig. 2-3. (b) Illustration of the initial 209 ms of the synthetic spike trains (fourth row) from which the PSDs in (a) were computed, and the components of the rate function that governed their generation (first-third rows; see Methods, Eq. 1 for full details). Term definitions: *p_ss_*(t) = steady state spiking probability, *p_RP_*(t) = recovery period spiking probability, *p_spk_*(t) = spike probability, reflecting the product of the steady state and recovery period components.

**Fig. S12.**
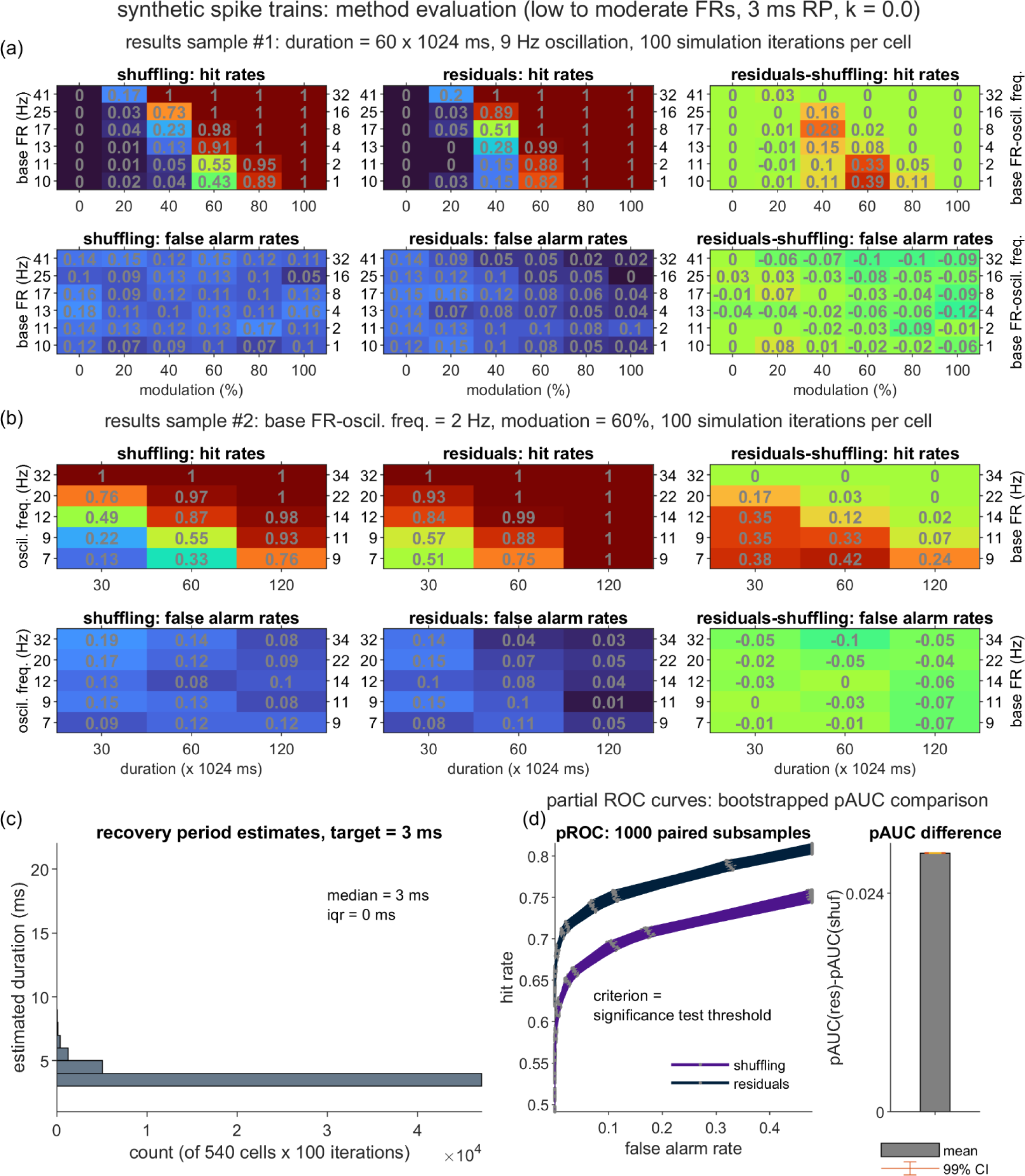
Performance of the shuffling and residuals methods over a synthetic dataset generated using a short, absolute recovery period. Panels (a)-(d): Plotting conventions, hit and false alarm definitions, and analysis procedures are identical to those described for the primary dataset depicted in Fig. 4. Relative to the primary dataset, this secondary dataset differed in the use of an absolute, as opposed to relative RP (k = 0), and a shorter RP duration (nr = 3 ms).

**Fig. S13.**
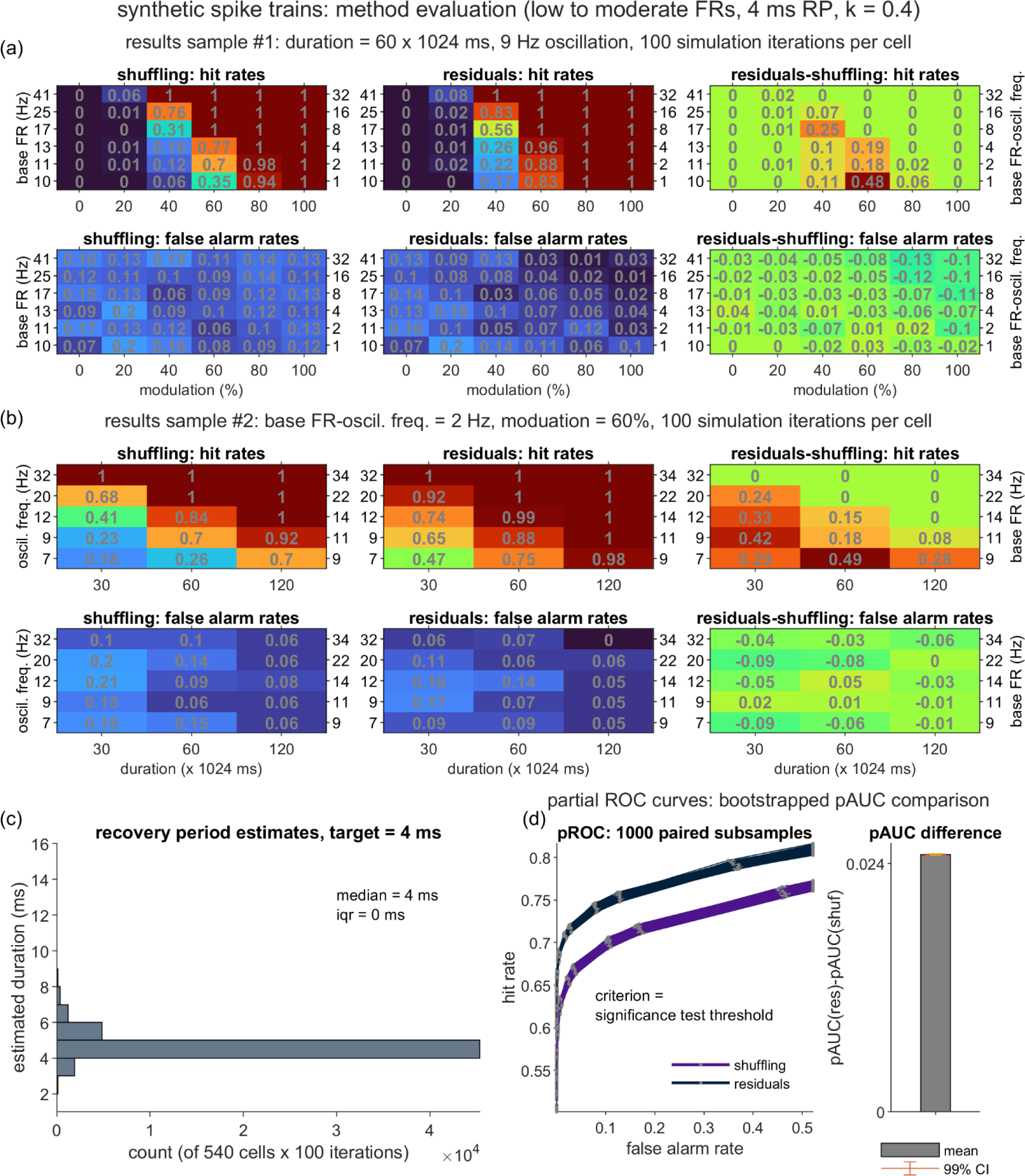
Performance of the shuffling and residuals methods over a synthetic dataset generated using a shortened relative recovery period. Panels (a)-(d): Plotting conventions, hit and false alarm definitions, and analysis procedures are identical to those described for the primary dataset depicted in Fig. 4. Relative to the primary dataset, this secondary dataset differed in the use of a shortened and steeper relative RP (*n_r_* = 4 ms, *k* = 0.4).

**Fig. S14.**
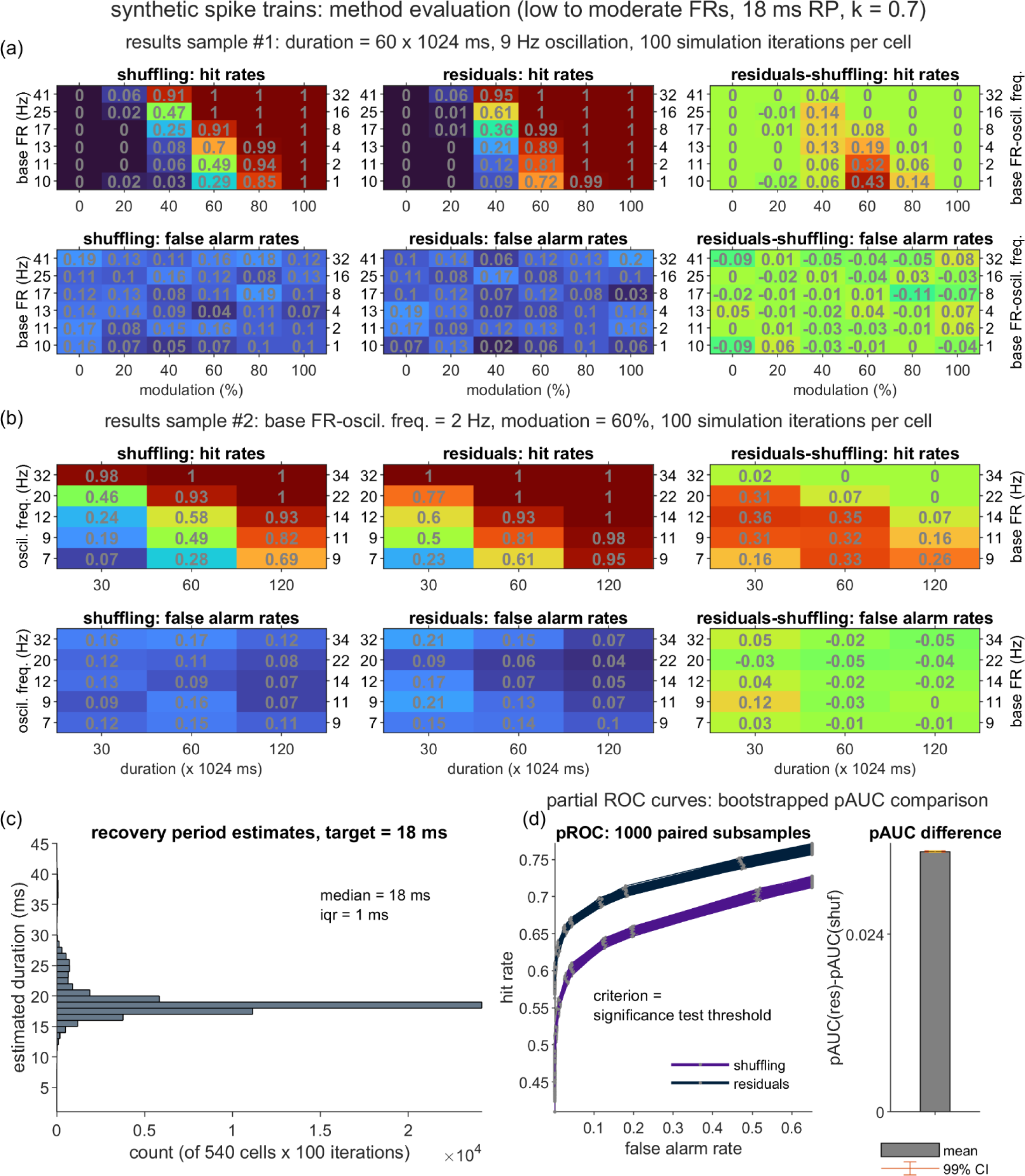
Performance of the shuffling and residuals methods over a synthetic dataset generated using a lengthened relative recovery period. Panels (a)-(d): Plotting conventions, hit and false alarm definitions, and analysis procedures are identical to those described for the primary dataset depicted in Fig. 4. Relative to the primary dataset, this secondary dataset differed in the use of a longer relative RP (nr = 18 ms, k = 0.7).

**Fig. S15.**
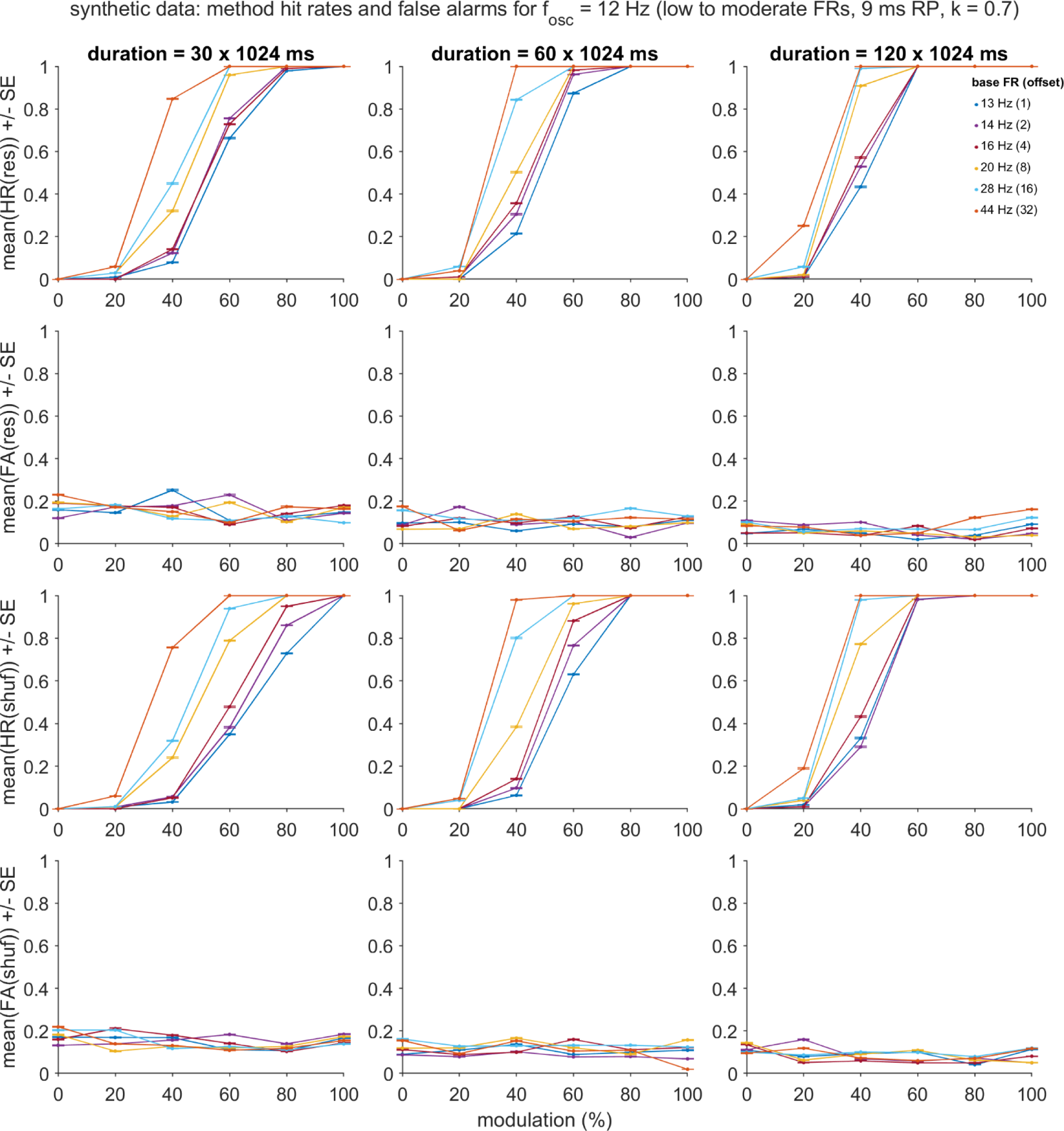
Residuals and shuffling hit and false alarm rates for the primary synthetic dataset, with the oscillation frequency (*f_osc_*) fixed at 12 Hz. Abbreviations and plotting conventions follow from those described for Fig. S9-S10.

**Fig. S16.**
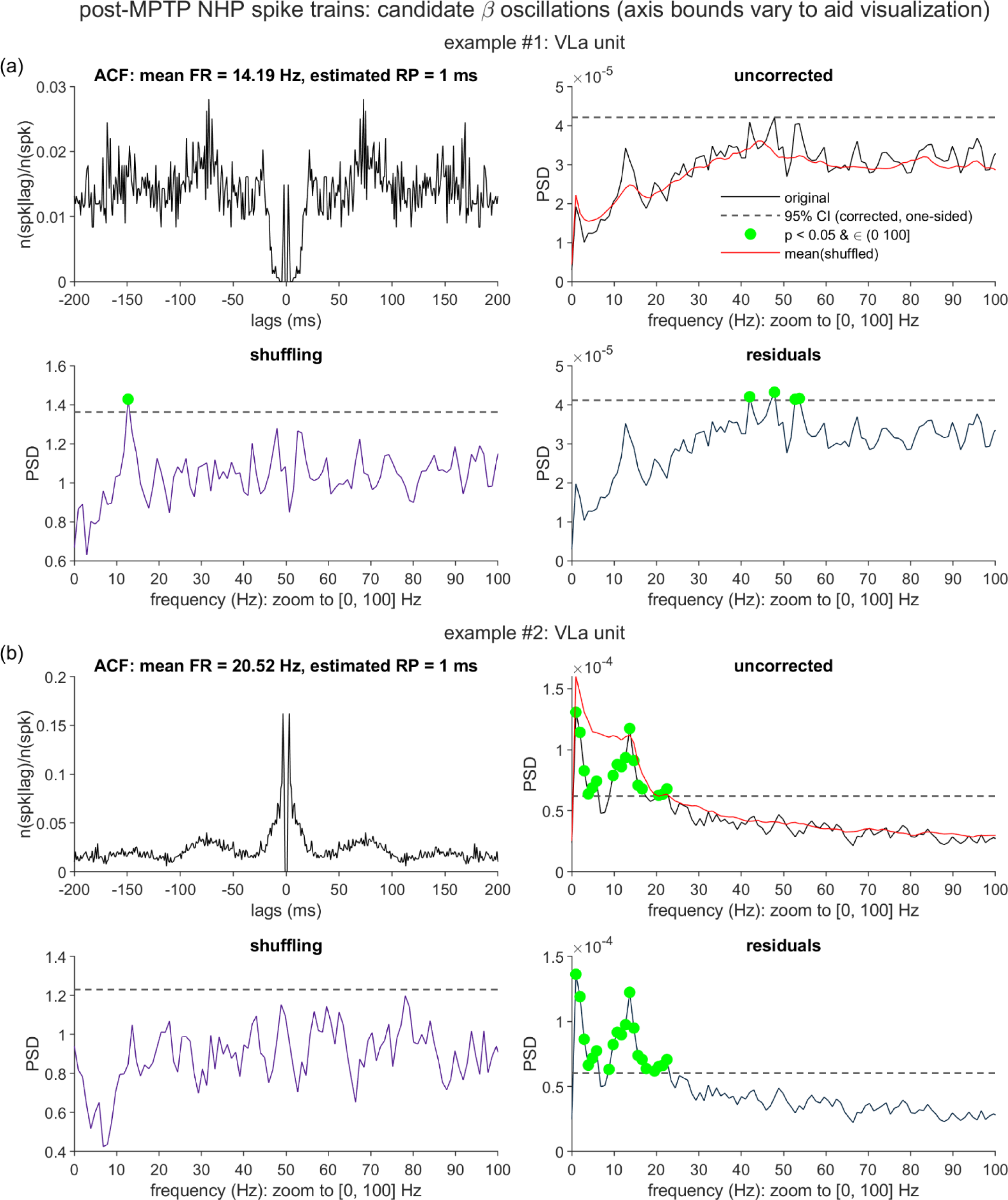
Comparison of the shuffling and residuals output for two units with putative beta oscillations and non-oscillatory features. Spike trains originated from two ventrolateral anterior thalamus (VLa) units, which had been recorded from the same parkinsonian non-human primate (NHP) that contributed to Fig. 5-6. Panels (a)-(b): Abbreviations and the statistical and plotting conventions follow from those described for Fig. 5.

**Fig. S17.**
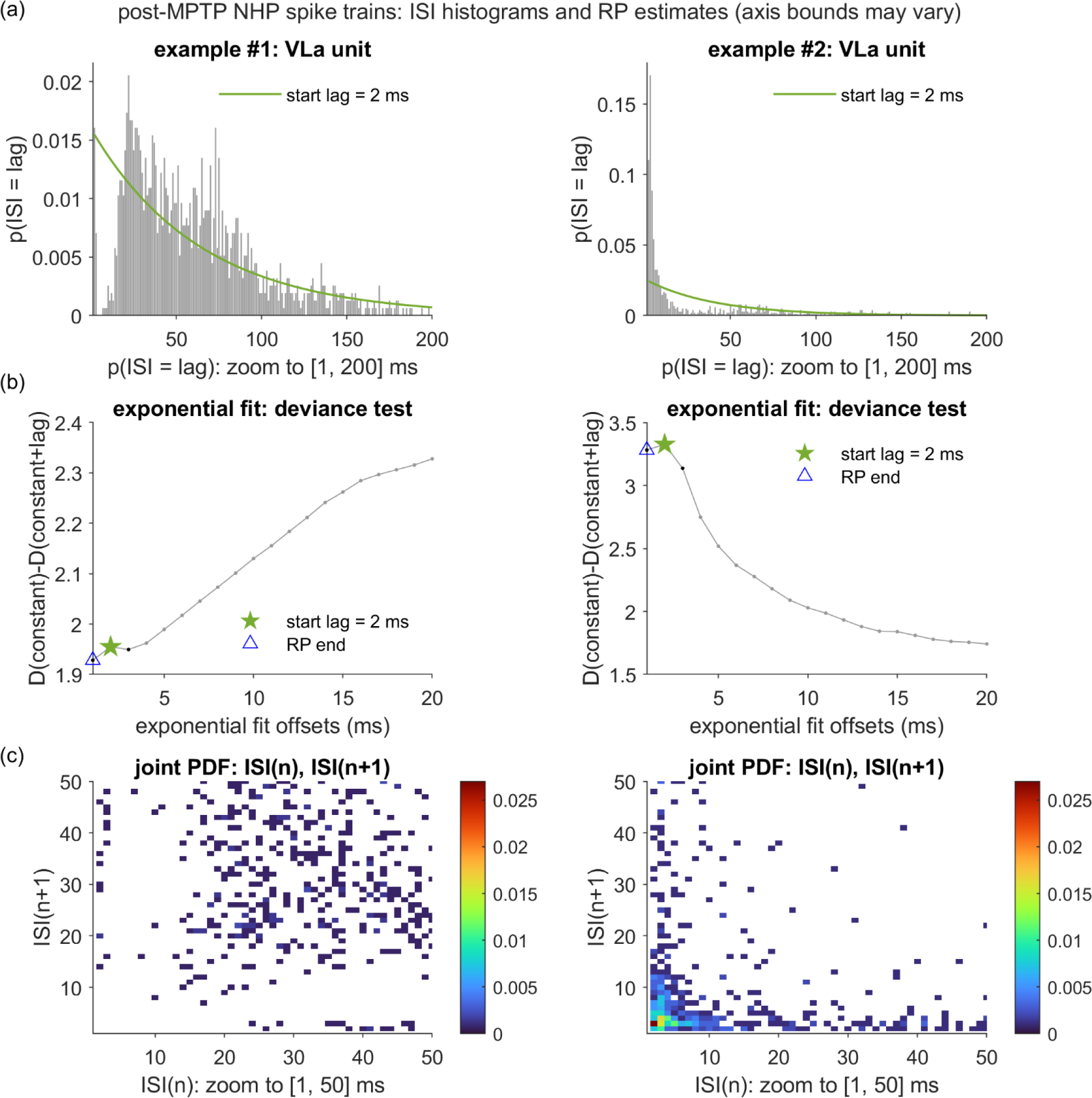
Inter-spike interval distributions and RP duration estimation for the two VLa units with suspected oscillatory and non-oscillatory features. The left and right columns depict the same two empirical spike trains as were presented in Fig. S16(a)-(b). (a) Illustration of the procedure for obtaining the estimated RP duration (*n̂*_*r*_) for the two spike trains. See Fig. 3(a)-(b) and the Methods for full details. For each unit, the figure highlights the exponential curve with the left anchor position (“start lag”) that returned the first local maximum in the curve goodness-of-fit plot. (b) Plots of the goodness-of-fit measure (the deviance difference statistic, ΔD) as a function of the first 20 starting positions of the exponential fits. Each *n̂*_*r*_ estimate (blue triangles) was set equal to the post-spike lag immediately preceding the first local maximum in the corresponding ΔD plot (green stars). (c) Joint probability density function (PDF) for all consecutive [ISI_n_, ISI_n+1_] inter-spike interval pairs. All remaining abbreviations follow from those described in Fig. 3 and Fig. 5.

### Supporting Tables

**Table S1.** Parametric Effects on Residuals-Shuffling Hit Rates Low-to-Moderate Firing Rates (9 ms RP, k = 0.7)

| Effect | Beta | SE | t | p |
| --- | --- | --- | --- | --- |
| Intercept | 0.084 | 0.00017 | 494.17 | -- |
| T | -0.00086 | 4.6e-06 | -188.48 | -- |
| $f_{osc}$ | -0.0049 | 1.9e-05 | -263.45 | -- |
| $p_{base\_offset}$ | -0.0046 | 1.6e-05 | -292.25 | -- |
| m | -0.17 | 0.0012 | -143.68 | -- |
| $m^2$ | -0.48 | 0.0016 | -296.41 | -- |
| T x $f_{osc}$ | 3.4e-05 | 5e-07 | 68.25 | -- |
| T x $p_{base\_offset}$ | 3.5e-05 | 4.2e-07 | 82.42 | -- |
| $f_{osc}$ x $p_{base\_offset}$ | 0.00027 | 1.7e-06 | 155.21 | -- |
| T x m | -0.0024 | 3.1e-05 | -77.28 | -- |
| $f_{osc}$ x m | -0.0028 | 0.00013 | -21.75 | 7.6e-105 |
| $p_{base\_offset}$ x m | 0.00057 | 0.00011 | 5.29 | 1.2e-07 |
| T x $m^2$ | 0.0049 | 4.4e-05 | 112.53 | -- |
| $f_{osc}$ x $m^2$ | 0.03 | 0.00018 | 168.36 | -- |
| $p_{base\_offset}$ x $m^2$ | 0.029 | 0.00015 | 191.37 | -- |
| m x $m^2$ | 1.1 | 0.0081 | 132.67 | -- |
| T x $f_{osc}$ x $p_{base\_offset}$ | -9.9e-07 | 3e-08 | -33.48 | 2e-245 |
| T x $f_{osc}$ x m | 8.4e-05 | 1.1e-06 | 73.92 | -- |
| T x $p_{base\_offset}$ x m | 6.4e-05 | 9.5e-07 | 66.88 | -- |
| $f_{osc}$ x $p_{base\_offset}$ x m | 0.00017 | 3.9e-06 | 44.01 | -- |
| T x $f_{osc}$ x $m^2$ | -0.00015 | 4.8e-06 | -32.06 | 3.3e-225 |
| T x $p_{base\_offset}$ x $m^2$ | -0.00016 | 4e-06 | -40.27 | -- |
| $f_{osc}$ x $p_{base\_offset}$ x $m^2$ | -0.0015 | 1.7e-05 | -90.54 | -- |
| T x m x $m^2$ | 0.013 | 0.00022 | 60.29 | -- |
| $f_{osc}$ x m x $m^2$ | 0.0075 | 0.00088 | 8.54 | 1.3e-17 |
| $p_{base\_offset}$ x m x $m^2$ | -0.012 | 0.00074 | -16.60 | 7e-62 |

RP = recovery period; T = duration;
$f_{osc}$ = oscillation frequency;
$p_{base\_offset}$ = offset of base firing rate from $f_{osc}$ ;
m = oscillation modulation
strength;
-- = outside range of double representation

**Table S2.** Parametric Effects on Residuals-Shuffling False Alarms Low-to-Moderate Firing Rates (9 ms RP, k = 0.7)

| Effect | Beta | SE | t | p |
| --- | --- | --- | --- | --- |
| Intercept | -0.023 | 0.00017 | -138.04 | -- |
| T | -0.00034 | 4.5e-06 | -75.60 | -- |
| $f_{osc}$ | -0.0012 | 1.8e-05 | -64.95 | -- |
| $p_{base\_offset}$ | -4.3e-05 | 1.5e-05 | -2.81 | 0.0049 |
| m | -0.0026 | 0.0011 | -2.28 | 0.023 |
| $m^2$ | 0.095 | 0.0016 | 59.43 | -- |
| T x $f_{osc}$ | 1.2e-05 | 4.9e-07 | 24.87 | 1.9e-136 |
| T x $p_{base\_offset}$ | 1.1e-05 | 4.1e-07 | 27.19 | 1.2e-162 |
| $f_{osc}$ x $p_{base\_offset}$ | 4.3e-06 | 1.7e-06 | 2.54 | 0.011 |
| T x m | 0.001 | 3e-05 | 34.38 | 9.8e-259 |
| $f_{osc}$ x m | -0.0031 | 0.00012 | -25.25 | 1.4e-140 |
| $p_{base\_offset}$ x m | 0.0055 | 0.0001 | 52.64 | -- |
| T x $m^2$ | 0.00053 | 4.3e-05 | 12.32 | 6.9e-35 |
| $f_{osc}$ x $m^2$ | -0.0027 | 0.00017 | -15.26 | 1.4e-52 |
| $p_{base\_offset}$ x $m^2$ | 0.0031 | 0.00015 | 21.28 | 2.1e-100 |
| m x $m^2$ | 0.031 | 0.0079 | 3.93 | 8.6e-05 |
| T x $f_{osc}$ x $p_{base\_offset}$ | -1e-06 | 2.9e-08 | -34.66 | 7.9e-263 |
| T x $f_{osc}$ x m | 2.3e-05 | 1.1e-06 | 20.65 | 1e-94 |
| T x $p_{base\_offset}$ x m | 2.1e-05 | 9.3e-07 | 22.26 | 9.5e-110 |
| $f_{osc}$ x $p_{base\_offset}$ x m | -0.00025 | 3.8e-06 | -65.86 | -- |
| T x $f_{osc}$ x $m^2$ | 2.1e-05 | 4.7e-06 | 4.39 | 1.1e-05 |
| T x $p_{base\_offset}$ x $m^2$ | -1.5e-05 | 3.9e-06 | -3.72 | 0.0002 |
| $f_{osc}$ x $p_{base\_offset}$ x $m^2$ | -0.00043 | 1.6e-05 | -26.54 | 4.8e-155 |
| T x m x $m^2$ | -0.0034 | 0.00021 | -16.24 | 2.8e-59 |
| $f_{osc}$ x m x $m^2$ | 0.0067 | 0.00086 | 7.73 | 1e-14 |
| $p_{base\_offset}$ x m x $m^2$ | -0.027 | 0.00073 | -37.01 | 2.5e-299 |

RP = recovery period; T = duration;
$f_{osc}$ = oscillation frequency;
$p_{base\_offset}$ = offset of base firing rate from $f_{osc}$ ;
m = oscillation modulation
strength;
-- = outside range of double representation

**Table S3.** Parametric Effects on Residuals-Shuffling Hit Rates Low-to-Moderate Firing Rates (9 ms RP, k = 0.7)

| Effect | Beta | SE | t | p |
| --- | --- | --- | --- | --- |
| Intercept | 0.1 | 0.0003 | 341.42 | -- |
| T | -0.00099 | 8e-06 | -124.45 | -- |
| $p_{base}$ | -0.0052 | 0.00012 | -43.21 | -- |
| $p_{base\_offset}$ | -0.008 | 0.00013 | -59.81 | -- |
| m | -0.44 | 0.002 | -217.82 | -- |
| $m^2$ | -0.62 | 0.0029 | -217.13 | -- |
| T x $p_{base}$ | 8.9e-05 | 3.2e-06 | 27.85 | 3.5e-170 |
| T x $p_{base\_offset}$ | 4e-05 | 3.6e-06 | 11.29 | 1.5e-29 |
| $p_{base}$ x $p_{base\_offset}$ | 0.00036 | 5.3e-05 | 6.66 | 2.7e-11 |
| T x m | -0.004 | 5.4e-05 | -72.62 | -- |
| $p_{base}$ x m | -0.018 | 0.00082 | -22.29 | 7.4e-110 |
| $p_{base\_offset}$ x m | 0.013 | 0.00091 | 14.57 | 5e-48 |
| T x $m^2$ | 0.0079 | 7.7e-05 | 103.32 | -- |
| $p_{base}$ x $m^2$ | 0.036 | 0.0011 | 31.79 | 6.4e-221 |
| $p_{base\_offset}$ x $m^2$ | 0.053 | 0.0013 | 41.28 | -- |
| m x $m^2$ | 2.7 | 0.014 | 193.84 | -- |
| T x $p_{base}$ x $p_{base\_offset}$ | -7.2e-06 | 9.2e-07 | -7.81 | 5.8e-15 |
| T x $p_{base}$ x m | 2.2e-05 | 7.3e-06 | 3.03 | 0.0025 |
| T x $p_{base\_offset}$ x m | 0.00023 | 8.1e-06 | 28.19 | 3e-174 |
| $p_{base}$ x $p_{base\_offset}$ x m | -4.1e-05 | 0.00012 | -0.34 | 0.74 |
| T x $p_{base}$ x $m^2$ | -0.00045 | 3.1e-05 | -14.64 | 1.8e-48 |
| T x $p_{base\_offset}$ x $m^2$ | -0.00062 | 3.4e-05 | -17.96 | 5.3e-72 |
| $p_{base}$ x $p_{base\_offset}$ x $m^2$ | -0.0019 | 0.00051 | -3.77 | 0.00016 |
| T x m x $m^2$ | 0.024 | 0.00038 | 64.06 | -- |
| $p_{base}$ x m x $m^2$ | 0.11 | 0.0057 | 20.26 | 4e-91 |
| $p_{base\_offset}$ x m x $m^2$ | -0.084 | 0.0063 | -13.21 | 7.7e-40 |

RP = recovery period; T = duration;
$p_{base}$ = base firing rate;
$p_{base\_offset}$ = offset of base firing rate from $f_{osc}$ ;
m = oscillation modulation strength;
-- = outside range of double representation

**Table S4.**
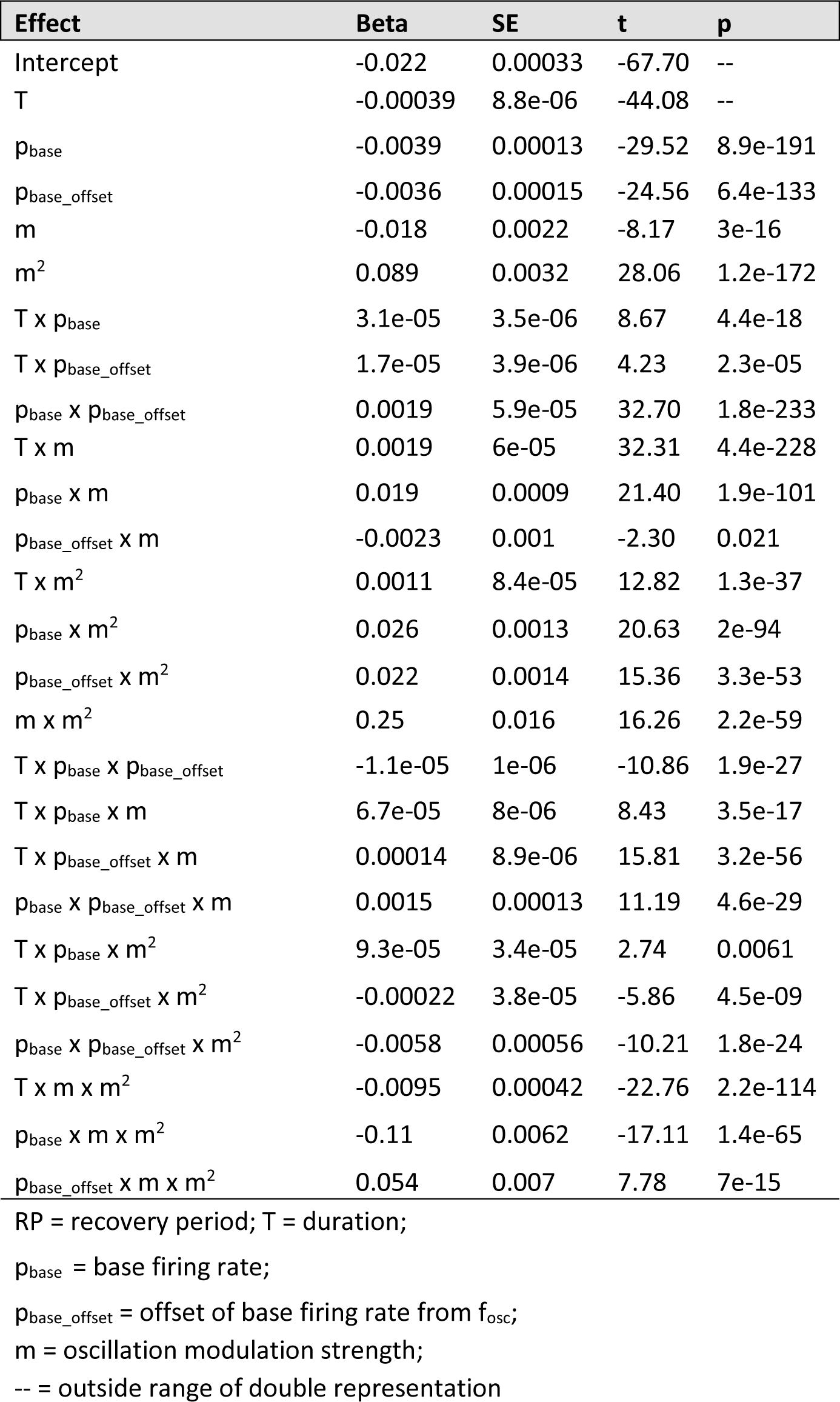
Parametric Effects on Residuals-Shuffling False Alarms Low-to-Moderate Firing Rates (9 ms RP, k = 0.7)

| Effect | Beta | SE | t | p |
| --- | --- | --- | --- | --- |
| Intercept | -0.022 | 0.00033 | -67.70 | -- |
| T | -0.00039 | 8.8e-06 | -44.08 | -- |
| $p_{\text{base}}$ | -0.0039 | 0.00013 | -29.52 | 8.9e-191 |
| $p_{\text{base\_offset}}$ | -0.0036 | 0.00015 | -24.56 | 6.4e-133 |
| m | -0.018 | 0.0022 | -8.17 | 3e-16 |
| $m^2$ | 0.089 | 0.0032 | 28.06 | 1.2e-172 |
| T x $p_{\text{base}}$ | 3.1e-05 | 3.5e-06 | 8.67 | 4.4e-18 |
| T x $p_{\text{base\_offset}}$ | 1.7e-05 | 3.9e-06 | 4.23 | 2.3e-05 |
| $p_{\text{base}}$ x $p_{\text{base\_offset}}$ | 0.0019 | 5.9e-05 | 32.70 | 1.8e-233 |
| T x m | 0.0019 | 6e-05 | 32.31 | 4.4e-228 |
| $p_{\text{base}}$ x m | 0.019 | 0.0009 | 21.40 | 1.9e-101 |
| $p_{\text{base\_offset}}$ x m | -0.0023 | 0.001 | -2.30 | 0.021 |
| T x $m^2$ | 0.0011 | 8.4e-05 | 12.82 | 1.3e-37 |
| $p_{\text{base}}$ x $m^2$ | 0.026 | 0.0013 | 20.63 | 2e-94 |
| $p_{\text{base\_offset}}$ x $m^2$ | 0.022 | 0.0014 | 15.36 | 3.3e-53 |
| m x $m^2$ | 0.25 | 0.016 | 16.26 | 2.2e-59 |
| T x $p_{\text{base}}$ x $p_{\text{base\_offset}}$ | -1.1e-05 | 1e-06 | -10.86 | 1.9e-27 |
| T x $p_{\text{base}}$ x m | 6.7e-05 | 8e-06 | 8.43 | 3.5e-17 |
| T x $p_{\text{base\_offset}}$ x m | 0.00014 | 8.9e-06 | 15.81 | 3.2e-56 |
| $p_{\text{base}}$ x $p_{\text{base\_offset}}$ x m | 0.0015 | 0.00013 | 11.19 | 4.6e-29 |
| T x $p_{\text{base}}$ x $m^2$ | 9.3e-05 | 3.4e-05 | 2.74 | 0.0061 |
| T x $p_{\text{base\_offset}}$ x $m^2$ | -0.00022 | 3.8e-05 | -5.86 | 4.5e-09 |
| $p_{\text{base}}$ x $p_{\text{base\_offset}}$ x $m^2$ | -0.0058 | 0.00056 | -10.21 | 1.8e-24 |
| T x m x $m^2$ | -0.0095 | 0.00042 | -22.76 | 2.2e-114 |
| $p_{\text{base}}$ x m x $m^2$ | -0.11 | 0.0062 | -17.11 | 1.4e-65 |
| $p_{\text{base\_offset}}$ x m x $m^2$ | 0.054 | 0.007 | 7.78 | 7e-15 |

RP = recovery period; T = duration;
$p_{\text{base}}$ = base firing rate;
$p_{\text{base\_offset}}$ = offset of base firing rate from $f_{\text{osc}}$ ;
m = oscillation modulation strength;
-- = outside range of double representation

**Table S5.** Parametric Effects on Residuals-Shuffling Hit Rates High Firing Rates (9 ms RP, k = 0.7)

| Effect | Beta | SE | t | p |
| --- | --- | --- | --- | --- |
| Intercept | 0.0011 | 6.2e-05 | 18.48 | 3.4e-76 |
| T | -4.8e-05 | 1.7e-06 | -28.85 | 6.6e-183 |
| $f_{osc}$ | -0.00011 | 6.8e-06 | -15.83 | 1.9e-56 |
| $p_{base\_offset}$ | -0.00019 | 3.6e-06 | -51.56 | -- |
| m | -0.018 | 0.00042 | -42.45 | -- |
| $m^2$ | 0.045 | 0.00059 | 75.82 | -- |
| T x $f_{osc}$ | 2.6e-06 | 1.8e-07 | 14.60 | 2.8e-48 |
| T x $p_{base\_offset}$ | 5.1e-06 | 9.7e-08 | 52.69 | -- |
| $f_{osc}$ x $p_{base\_offset}$ | 5.1e-06 | 4e-07 | 12.96 | 2.2e-38 |
| T x m | 0.00056 | 1.1e-05 | 49.35 | -- |
| $f_{osc}$ x m | 0.00094 | 4.6e-05 | 20.44 | 8.3e-93 |
| $p_{base\_offset}$ x m | 0.0017 | 2.5e-05 | 69.09 | -- |
| T x $m^2$ | -0.00045 | 1.6e-05 | -28.05 | 5.7e-173 |
| $f_{osc}$ x $m^2$ | 0.0013 | 6.5e-05 | 20.03 | 3e-89 |
| $p_{base\_offset}$ x $m^2$ | 0.0018 | 3.5e-05 | 51.76 | -- |
| m x $m^2$ | -0.015 | 0.0029 | -5.27 | 1.4e-07 |
| T x $f_{osc}$ x $p_{base\_offset}$ | -1.5e-08 | 6.8e-09 | -2.22 | 0.026 |
| T x $f_{osc}$ x m | 5.5e-08 | 4.1e-07 | 0.13 | 0.89 |
| T x $p_{base\_offset}$ x m | 4.5e-06 | 2.2e-07 | 20.60 | 3.3e-94 |
| $f_{osc}$ x $p_{base\_offset}$ x m | -7.2e-06 | 9e-07 | -8.00 | 1.3e-15 |
| T x $f_{osc}$ x $m^2$ | -2.5e-05 | 1.7e-06 | -14.19 | 1.1e-45 |
| T x $p_{base\_offset}$ x $m^2$ | -6e-05 | 9.3e-07 | -64.42 | -- |
| $f_{osc}$ x $p_{base\_offset}$ x $m^2$ | -2.8e-05 | 3.8e-06 | -7.26 | 4e-13 |
| T x m x $m^2$ | -0.0017 | 7.8e-05 | -22.31 | 3.2e-110 |
| $f_{osc}$ x m x $m^2$ | -0.0078 | 0.00032 | -24.22 | 1.6e-129 |
| $p_{base\_offset}$ x m x $m^2$ | -0.013 | 0.00017 | -74.32 | -- |

RP = recovery period; T = duration;
$f_{osc}$ = oscillation frequency;
$p_{base\_offset}$ = offset of base firing rate from $f_{osc}$ ;
m = oscillation modulation strength;
-- = outside range of double representation

**Table S6.** Parametric Effects on Residuals-Shuffling False Alarms High Firing Rates (9 ms RP, k = 0.7)

| Effect | Beta | SE | t | p |
| --- | --- | --- | --- | --- |
| Intercept | -0.014 | 0.00022 | -63.65 | -- |
| T | 1.5e-05 | 6e-06 | 2.53 | 0.011 |
| $f_{osc}$ | -0.00052 | 2.4e-05 | -21.37 | 3e-101 |
| $p_{base\_offset}$ | 0.00024 | 1.3e-05 | 18.24 | 2.5e-74 |
| m | 0.12 | 0.0015 | 76.88 | -- |
| $m^2$ | 0.23 | 0.0021 | 107.57 | -- |
| T x $f_{osc}$ | -5.4e-06 | 6.5e-07 | -8.25 | 1.6e-16 |
| T x $p_{base\_offset}$ | -6.7e-06 | 3.5e-07 | -19.16 | 8.6e-82 |
| $f_{osc}$ x $p_{base\_offset}$ | 1.3e-05 | 1.4e-06 | 9.33 | 1e-20 |
| T x m | 4.2e-05 | 4.1e-05 | 1.04 | 0.3 |
| $f_{osc}$ x m | -0.0089 | 0.00017 | -53.48 | -- |
| $p_{base\_offset}$ x m | 0.002 | 8.9e-05 | 22.75 | 1.6e-114 |
| T x $m^2$ | 0.0005 | 5.7e-05 | 8.83 | 1e-18 |
| $f_{osc}$ x $m^2$ | -0.079 | 0.00023 | -336.62 | -- |
| $p_{base\_offset}$ x $m^2$ | 0.0057 | 0.00013 | 45.72 | -- |
| m x $m^2$ | -0.11 | 0.011 | -10.32 | 5.9e-25 |
| T x $f_{osc}$ x $p_{base\_offset}$ | -6.5e-07 | 2.4e-08 | -26.43 | 7.2e-154 |
| T x $f_{osc}$ x m | -0.00026 | 1.5e-06 | -178.70 | -- |
| T x $p_{base\_offset}$ x m | 2.8e-05 | 7.9e-07 | 35.75 | 1.9e-279 |
| $f_{osc}$ x $p_{base\_offset}$ x m | -0.00059 | 3.2e-06 | -181.70 | -- |
| T x $f_{osc}$ x $m^2$ | -0.00069 | 6.2e-06 | -110.94 | -- |
| T x $p_{base\_offset}$ x $m^2$ | 0.00015 | 3.3e-06 | 45.24 | -- |
| $f_{osc}$ x $p_{base\_offset}$ x $m^2$ | -0.0016 | 1.4e-05 | -116.58 | -- |
| T x m x $m^2$ | 0.006 | 0.00028 | 21.19 | 1.3e-99 |
| $f_{osc}$ x m x $m^2$ | -0.15 | 0.0012 | -129.27 | -- |
| $p_{base\_offset}$ x m x $m^2$ | 0.0062 | 0.00062 | 10.05 | 9.2e-24 |

RP = recovery period; T = duration;
$f_{osc}$ = oscillation frequency;
$p_{base\_offset}$ = offset of base firing rate from $f_{osc}$ ;
m = oscillation modulation strength;
-- = outside range of double representation

